# Quantifying pathogen spread across scales from pairwise genetic distance distributions

**DOI:** 10.64898/2026.11.18.767662

**Authors:** Noémie Lefrancq, Richard A. Neher, Tanja Stadler

**Affiliations:** Department of Biosystems Science and Engineering, ETH Zurich, Basel, Switzerland; SIB Swiss Institute of Bioinformatics, Lausanne, Switzerland; Biozentrum, University of Basel, Basel, Switzerland

## Abstract

Pathogen spread is structured across space, demographic groups, and host species. Yet quantifying movement between these states remains a central challenge of genomic epidemiology, particularly with uneven sampling. Here, we developed a scalable framework that quantifies symmetric transition rates from pairwise genetic distance distributions. In simulations, our framework recovered transition rates across diverse demographic histories and sampling schemes, and remained robust to uneven sampling intensities. Applied to SARS-CoV-2, it quantified international spread associated with human mobility and local transmission across age groups associated with social contacts. Analyses of influenza, respiratory syncytial virus, and MERS-CoV extended these results across pathogens and host systems. Together, these results establish a general framework for accurate and scalable inference of pathogen spread across scales.

## Main

Transmission dynamics of infectious diseases are inherently structured. Individual behaviour, population organisation and environmental context shape how pathogens move between states such as regions, demographic groups, and host species. Quantifying transition rates between such states is central to understanding epidemic dynamics, identifying drivers of spread and designing effective interventions. Yet, these rates are difficult to measure directly, as surveillance data are frequently incomplete or biased across groups, making it challenging to distinguish true differences in transmission from differences in observation (*1*).

Pathogen genomic data provide a complementary approach (*2*). As pathogens spread, they accumulate mutations that record their ancestry. Genetic differences among sampled infections, therefore, reflect the history of pathogen divergence and movement between states through time. This evolutionary signal underpins modern genomic epidemiology and phylodynamics (*3*). Most commonly, transitions between discrete states are modelled along the branches of reconstructed phylogenetic trees using continuous-time Markov chain models (*4, 5*). Alternatively, structured coalescent models (*6*) and multi-type birth-death models (*7*) incorporate population structure directly into the tree-generating process. These types of approaches have provided important insights into, for example, spatial spread (*8*) and mixing between populations (*9*). However, they face two major limitations. First, heterogeneous sampling across states can bias estimated transition rates (*10*). Second, explicit phylogenetic reconstruction and structured phylodynamic inference are computationally intensive, limiting scalability to the very large genomic datasets now routinely generated.

These limitations motivate methods that extract signals of transitions between states that are robust to uneven sampling, while remaining computationally tractable (*11, 12*). Pairwise genetic distance distributions can provide such signals, as they capture information about both time since shared ancestry and the movement of lineages across states. These distributions are constructed from genetic divergence between pairs of sequences, rather than counts of transitions between states, making them robust to uneven sampling. They can also be efficiently computed from aligned sequence data or genetic distance-based phylogenies. However, existing approaches have mostly used these distributions to describe qualitatively transmission between states, rather than to estimate transition rates.

Here, we introduce a scalable framework to quantify backwards-in-time symmetric transition rates between discrete population states directly from pairwise genetic distance distributions. We first use simulations to assess the performance of our approach, its robustness to demographic and sampling intensity, and its limits concerning directional inference. We then apply it to SARS-CoV-2 genomes to quantify large-scale spread between countries and fine-scale spread between age groups, and compare these estimates with human mobility and social contact patterns. Finally, we broaden the analysis to influenza A/H1N1pdm, influenza A/H3N2, RSV-A, RSV-B and Middle East respiratory syndrome coronavirus (MERS-CoV) genomes to quantify spread across age groups and host species.

### Signal of spread in pairwise genetic distance distributions

Individuals close together within a transmission chain are expected to be infected by genetically similar pathogens (Figure 1A). Pairwise genetic distance distributions across states therefore reflect both past transmission dynamics and transitions between states. For a given demographic history, transitions between states (*e*.*g*. migration between locations, transmission across age groups) shape how pairwise genetic distance distributions differ between sequences sampled from the same or different states (Figure 1B). At low transition rates, closely related sequences are concentrated within the same state, yielding different within- and between-state distributions (Figure 1C). As transition rates increase, these distributions become increasingly similar. The ratio of these distributions (see Methods, Eqn. 4) summarises this structure (Figure 1D) (*11, 12*). Critically, normalizing the distributions separately for each pair type makes them robust to unequal sampling across states, as they capture the distance distribution within each pair type rather than differences in pair counts (Figure S1). However, the same transition rate can generate different pairwise genetic distance distributions depending on past demographic histories, for example whether populations are dominated by recently or deeply diverged lineages. This highlights that a method to infer transition rates from these distributions must account for past within-state transmission dynamics (see Methods, Figure S2).

**Figure 1:**
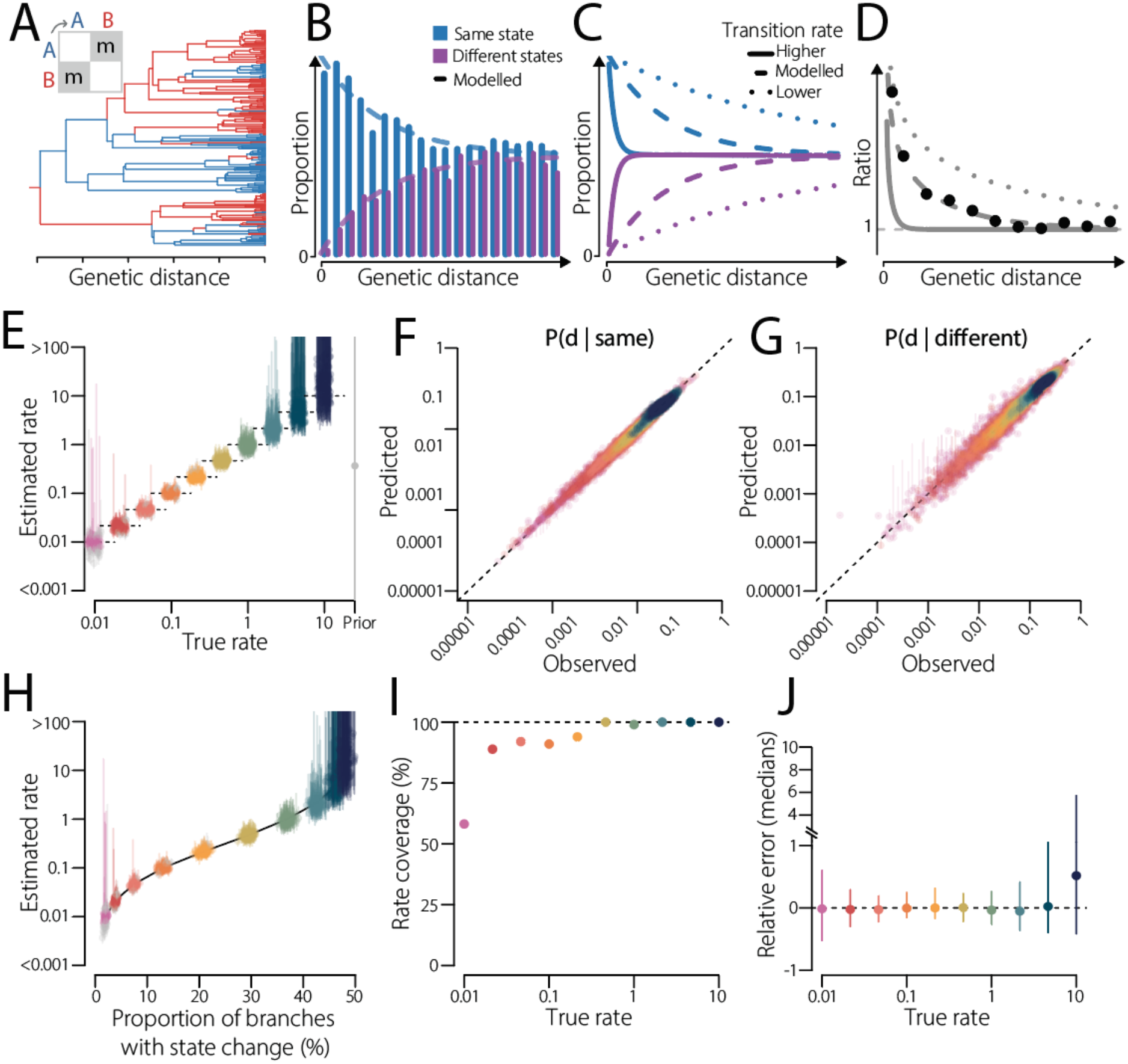
Migration signal in pairwise genetic distance distributions. **A**. Example of a phylogeny simulated under a structured coalescent model, with 2 states and one migration rate *m*between them. **B**. Example of pairwise genetic distance distributions for sequences from the same state (blue) and different states (purple), with the distributions predicted under the model (dashed lines). **C**. Pairwise distance distributions predicted by the model for a higher (solid), the modelled (dashed) and a lower (dotted) migration rate between states. **D**. Ratio of the two example distributions (dots) to summarise the differences, and ratios predicted by the model for the same three migration rates (lines). **E**. Estimated migration rates by our model on simulated data versus true rates. For each migration rate, 100 trees were simulated. Dashed lines represent the true rate. Estimates that do not cover the true rate are colored in grey. **F**. Observed versus predicted pairwise genetic distance distribution for sequences from the same state, colored by rate. **G**. Same as F, but for different states. **H**. Estimated rates as a function of the average proportion of branches with state change in each simulated phylogeny. The black line represents the overall average across all phylogenies. Estimates that do not cover the true rate are colored in grey. **I**. Percentage of simulations for which the true rate was covered by the credible intervals. The dashed line represents 100%. **J**. Median relative error across all simulations, for each true rate. The dashed line represents no relative error. In E-J, dots and lines are colored according to the true rate of the simulation. See methods for details on the simulations. Example of model fits are presented in Figure S3.

### Estimating the rate of transition between states from pairwise genetic distance distributions

We developed a continuous-time Markov chain model that links backwards-in-time migration between states to the distribution of pairwise genetic distances between states, while accounting for the broader genetic structure in the dataset (see Methods for details). We assume that sequence pairs contribute independently to the likelihood and that demographic and migration processes can be separated. We examine the implications of both approximations below.

We first assessed the performance of our approach on phylogenies simulated under a two-state structured coalescent model with symmetric migration and equal population dynamics (Figure 1A, Figure S3). We found that our framework consistently fitted the data and captured the migration rates (Figure 1E-G). The model fits and credible interval widths of the estimated rates are consistent with the phylogenetic signal (Figure 1F-H). At low migration rates, the signal depends strongly on where the few transitions between states occurred, leading to greater variability across datasets and a reduced credible interval coverage. At high rates, lineages mix before genetic differences accumulate, diluting the signal of state structure, yielding large credible intervals (Figure 1H). This shows that the range of possible migration rates that can be estimated depends on the genetic resolution of the data. Over the 1000 trees we simulated here, 95.9% of fits reached convergence (effective sample size >200), with 92.6% covering the true rates, ranging from 58.1% for low rates, to 100% for high rates (Figure 1I). The estimated rates are overall unbiased, with relative errors reflecting the strength of the phylogenetic signal in each dataset (Figure 1J).

### Robustness to transmission histories and sampling intensities

Our framework accounts for population dynamics without explicitly estimating or coupling them to migration. To assess whether variation in transmission history affects model estimates, we simulated phylogenies under fixed symmetric migration rates between two states across a wide range of scenarios: constant, exponentially growing, seasonal, and unequal population dynamics (Figure 2 A-B and S4). These demographic histories altered coalescent times and, consequently, the pairwise distance distributions generated by the same migration rate. Nevertheless, our method recovered the true migration rates and remained unbiased, with the credible interval width reflecting phylogenetic signal. These results support the appropriateness of our approximation that demographic and migration processes can be treated separately across a range of population dynamics.

**Figure 2:**
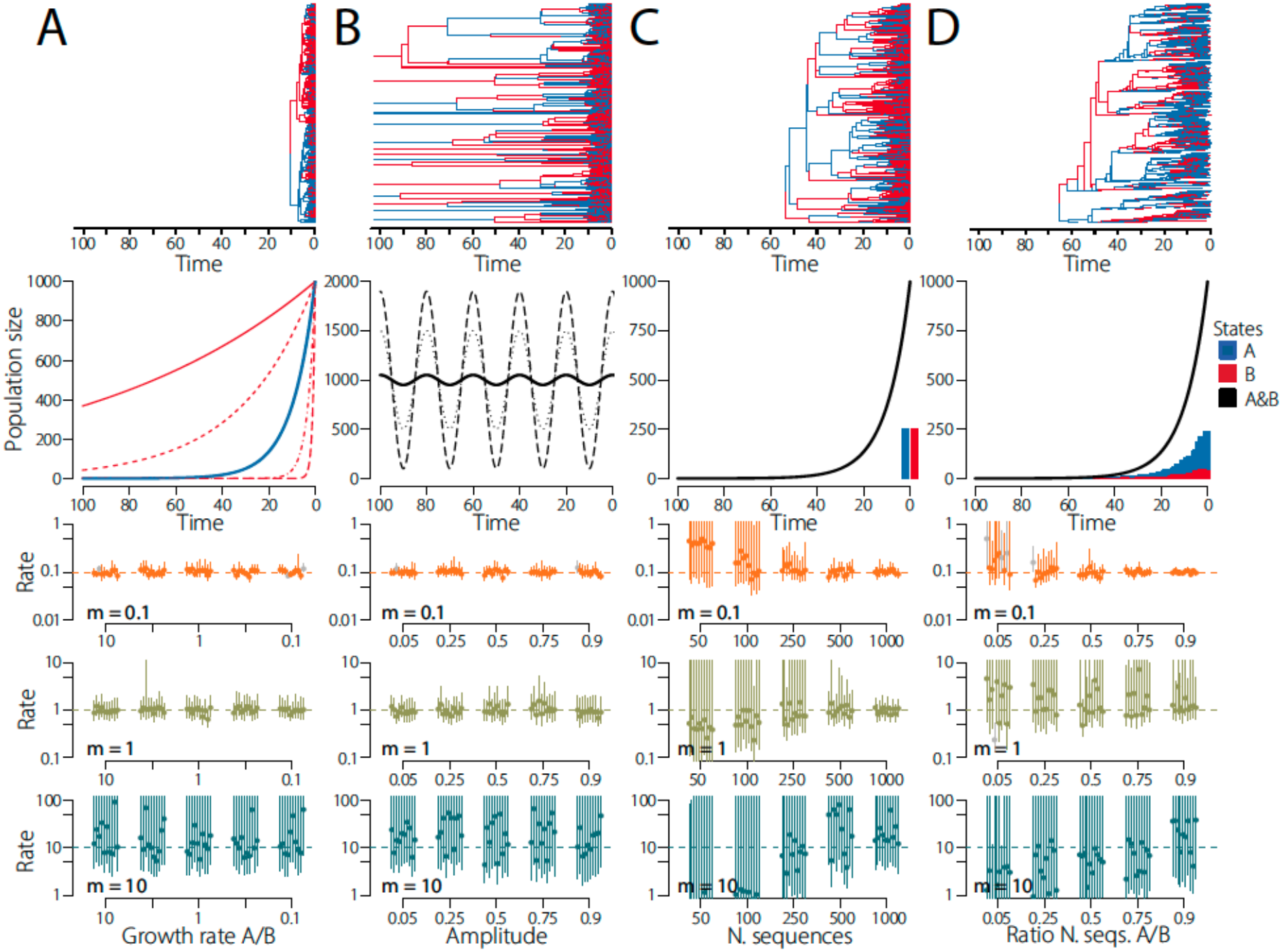
Robustness to different transmission histories and different sampling schemes. **A-B**. Impact of different transmission intensities across demes. A. Exponentially growing effective population size. The effective population size of state A is growing at a rate of 0.1 per unit of time (blue), and we vary the growth rate of the effective population size of state B (red). B. Periodically varying effective population size. Both states A and B have the same effective population size, and we vary the amplitude across simulations. **C-D**. Impact of different sampling schemes. C. Both states are equally sampled at one time point, and we vary the number of sequences sampled across simulations. D. States are sampled at a constant rate through time, proportional to the effective population size. Across simulations, state A is always sampled at the same rate, while we vary the sampling rate of state B, effectively introducing a sampling bias. The first row of panels presents examples of simulated phylogenies in each scenario. The second row presents the different effective population sizes used (lines, blue: state A, red: B, black: both states). Bars indicate examples of the number of sampled sequences by state. The third to fifth rows present the migration rates estimated by our approach, for each scenario and for three different true migration rates (0.1, 1 and 10 migrations per unit of time). Dots indicate median estimates, and bars indicate 95% credible intervals. The dashed lines represent true rates. Estimates that do not cover the true rates are coloured in grey. See methods for details about the simulations. Additional scenarios are presented in Figures S4-5.

We next evaluated robustness to sampling heterogeneity, a key feature of pathogen genomic datasets. We considered several sampling schemes, including sampling at a single time point, at multiple discrete time points, continuously through time, and proportional to incidence (Figure 2C and S5). We found that varying the total number of sequences or the sampling regime generally did not introduce systematic bias. Nonetheless, in the smallest datasets, estimates were noisier and more strongly influenced by the prior. Although pairwise-distance calculation scales quadratically with the number of sequences, inference depends only on the number of bins and states and therefore remains efficient for large datasets.

We further tested the effect of uneven sampling across states by varying the number of sequences obtained from each state. Such imbalance did not bias inferred migration rates (Figure 2D), in contrast to standard ancestral state reconstruction tools (Figure S6) (*10*). This is consistent with our method relying on the structure of pairwise distance distributions rather than on sampling frequencies.

### Estimating symmetric and asymmetric transition rate matrices

Beyond two-state systems, our framework can in principle estimate symmetric matrices of transition rates across multiple states. We therefore investigated how reliably individual rates can be recovered in practice. Under symmetric migration and equal population dynamics, our approach recovered the individual rates in three, four and five state simulations, including rates spanning two orders of magnitude (Figure S7 and S8). However, unequal demography can bias estimates in three or more state systems (see Methods and Figure S9). Migration rates between weakly connected states are particularly sensitive to coalescence in other states, where smaller effective population sizes shortened genetic distances and could lead to overestimated migration rates (Figure S9). Estimates between strongly connected states remained accurate. This confounding is consistent with our method modelling migration separately from demographic processes, as in discrete trait phylogeographic models (*e*.*g*., (*4*)). In contrast, unequal sampling intensity did not introduce systematic bias (Figure S10).

Finally, we examined the identifiability of asymmetric migration. We found that the overall movement between states can be estimated, while not its direction. We exemplify this limitation in the case of two states: pairwise distance distributions depend on the directional rates only through symmetric combinations and remain unchanged when the rates are exchanged (see Methods). In simulations with the same total migration rate, ranging from symmetric to one-way migration, symmetric models recovered the average of the two rates, while asymmetric models estimated similar rates in both directions with wide credible intervals, regardless of the true asymmetry (Figure S11). This limitation extends to larger systems, where distinct asymmetric migration matrices can produce nearly identical pairwise distance distributions (see Methods and Figure S12). Our approach is therefore best suited to estimating symmetric transition rates and overall movement between states, with directional inference requiring external information.

### Quantifying pathogen spread between countries

We next asked whether our approach could recover international patterns of pathogen movement at scale. We applied our framework to 15,991 SARS-CoV-2 whole genomes from NCBI, sampled from July 2024 to June 2025 in seven countries: Denmark, Japan, Slovakia, Switzerland, Thailand, the United Kingdom, and the United States (Table S1). Despite marked heterogeneity in both the number of genomes available from each country (Figure 3A) and in their temporal distribution (Figure 3B), we found a clear signature of spatial spread (Figure 3C). Overall, the proportion of genomes within countries that are identical was 78.7 (95%CI: 34.0-189.5) times higher than that of different countries. This ratio decreased to one for pairs of sequences separated by 4-5 mutations, consistent with transient geographic clustering.

**Figure 3:**
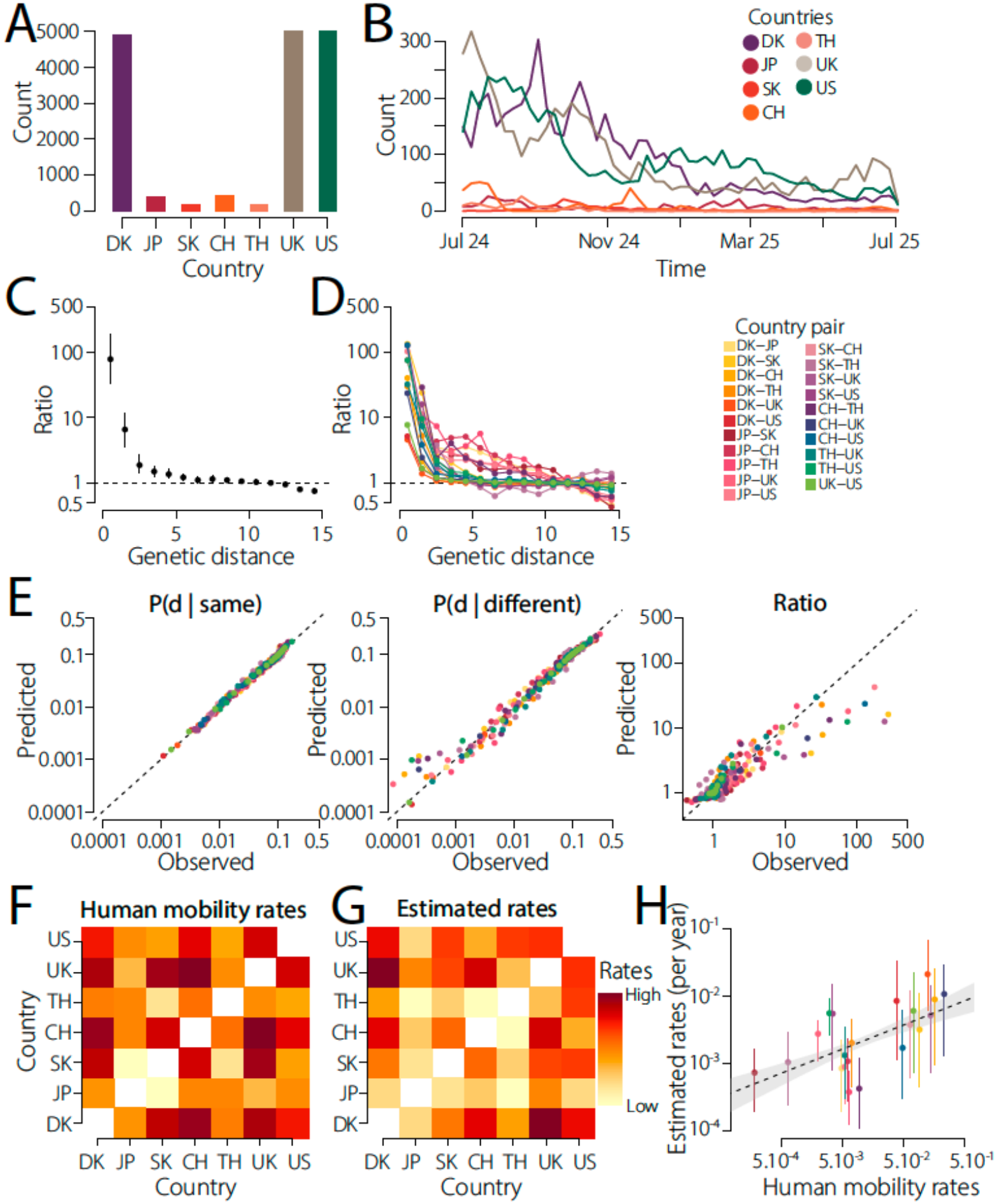
Quantifying SARS-CoV-2 spread between countries. **A**. Number of sequences per country. **B**. Number of sequences through time, by country. **C**. Ratio of pairwise genetic distance distributions for sequences from the same country *vs*. different countries, all country pairs combined. **D**. Same as C, but detailed by country pair. In C and D, the dashed lines denote a ratio of one (*i*.*e*. no difference), dots denote the median, and bars denote the 95% confidence intervals obtained by bootstrap. **E**. Model fit of the pairwise genetic distance distributions for sequences from the same country (left), different countries (centre) and ratio (right), coloured by country pair. The dashed line represents identity. Model fits by country pair are presented in Figure S13. **F**. Human mobility rates between countries as estimated by (*25*). **G**. Estimated transition rates per year between countries by our approach. **H**. Pairwise comparison between human mobility rates per year (x axis) and estimated transition rates per year by our approach (y axis). Estimates were rescaled in units of time using a SARS-CoV-2 substitution rate of 5.10-4 subs/site/year (*26, 27*). Dots indicate median estimates, and bars indicate 95% credible intervals, coloured by country pair. The dashed line represents a log-log regression, and the shaded area the confidence region, accounting for the matrix structure. Individual transition rates and comparison to other covariates are presented in Figure S14. The phylogeny coloured by country is presented in Figure S20.

We further observed a consistent geographical signal across country pairs (Figure 3D), while also detecting differences that revealed heterogeneity in connectivity between countries. For example, we found that for Denmark-UK, the proportion of identical genomes within countries was (95%CI: 3.2-6.6) times higher than that of across countries, but for Thailand-US, this ratio was 75.8 (95%CI: 17.0-584.4). Fitting our model to the pairwise distance distributions between countries, we reconstructed the observed distributions, and their ratio across country pairs (Figure 3E and S13).

The resulting estimates uncovered substantial heterogeneity in connectivity among country pairs (Figure 3F-G, and S14A). We found that Denmark-UK was the most connected pair of countries, with an estimated transition rate of 0.02 per year (95%CrI: 0.006-0.066), while Japan-Thailand was the least connected pair, with an estimated rate of 0.0004 per year (95%CrI: 0.0001-0.0007). These estimates were positively associated with independent measures of human mobility between countries (Figure 3H, permutation test, p value = 0.02), consistent with other studies (*13, 14*). We also found an association with the number of genomes available for each country pair (Figure S14B, p value = 0.01), however sensitivity analyses showed that estimates remained stable when varying the number of sequences sampled per location (Figure S15), supporting the robustness of our approach to uneven sampling. Estimates were not significantly associated with geographic distance between countries (Figure S14C, p value = 0.07).

### SARS-CoV-2 fine-scale transmission structure across age groups

We next investigated whether the same framework could resolve finer-scale social structure (*12*). As age strongly shapes contact patterns for respiratory infections, we applied our framework to 1,228 SARS-CoV-2 genomes from the National Reference Center (NRC) of Respiratory Viruses in Lyon, France, collected between July 2024 and June 2025, for which age-group information was available (Figure 4A and B). We used five age groups: 0–19 years, 20–39 years, 40–59 years, 60–79 years and 80 years and above. Despite substantial heterogeneity in sampling over time and across ages, pairwise distance distributions again revealed a clear signature of structured spread (Figure 4C). Overall, we found that the proportion of identical genomes within age groups was 2.2 (95%CI: 1.4-3.8) times higher than that of between age groups.

**Figure 4:**
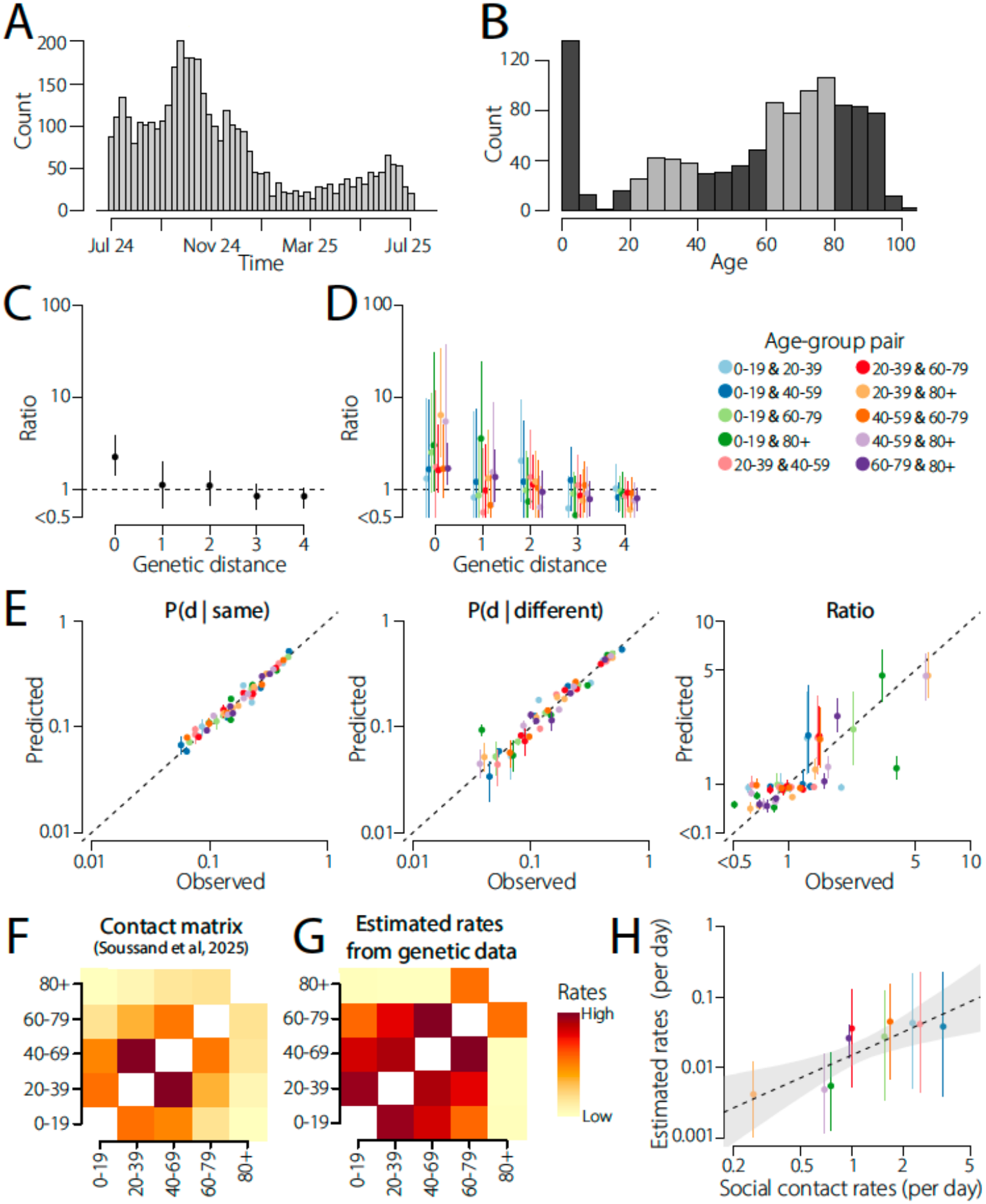
Quantifying SARS-CoV-2 spread between age groups. **A**. Number of SARS-CoV-2 sequences through time from Auverge-Rhône-Alpes region, France. **B**. Number of sequences with age information available by age. Shades of grey indicate the different age groups used. **C**. Ratio of pairwise genetic distance distributions for sequences from the same age group *vs* different age groups, all age groups combined. **D**. Same as C, but detailed by age-group pair. In C and D, the dashed lines denote a ratio of 1 (*i*.*e*. no difference), dots denote the median, and bars denote the 95% confidence intervals obtained by bootstrap. **E**. Model fit of the pairwise genetic distance distributions for sequences from the same age group (left), different age groups (centre) and ratio (right), coloured by age-group pair. The dashed line represents identity. **F**. Contact matrix between age groups, representing the number of contacts per day between individuals from different age groups, as estimated by (*28*). **G**. Estimated transition rates per day between age groups by our approach. **H**. Pairwise comparison between social contact rates (x axis) and estimated transition rates by our approach (y axis). Estimates were rescaled in units of time using a SARS-CoV-2 substitution rate of 5.10-4 subs/site/year (*26, 27*). Dots indicate median estimates, and bars indicate 95% credible intervals, coloured by age-group pair. The dashed line represents a log-log regression, and the shaded area the confidence region, accounting for the matrix structure. Comparison to other covariates is presented in Figure S16-17. The phylogeny coloured by age group is presented in Figure S21.

However, this ratio decayed much more rapidly with genetic distance and approached one within a 1-2 mutations (Figure 4C), consistent with a much faster SARS-CoV-2 spread locally between age groups than between countries worldwide. Across age-group pairs, the pattern was consistent but varied in magnitude, indicating heterogeneous connectivity among age classes (Figure 4D). For example, we found that for the age-group pair 20-39 and 80+, the proportion of identical genomes within age group was 6.4 (95%CI: 2.3-33.2) times higher than that of across age groups, but the age-group pair 0-19 and 20-39, this ratio was 1.3 (95%CI: 0.5-9.6). This pattern indicates that children and the oldest adults are less tightly connected in transmission than children and younger adults.

Fitting our model to these pairwise distance distributions, we reconstructed the observed distributions, and their ratio across age-group pairs (Figure 4E). The inferred rates varied substantially among age groups, as expected from independent estimates of social contact rates (Figure 4F-G). We found that 20-39 and 80+ were the least connected, with an estimated transition rate of 0.004 per day (95%CrI: 0.001-0.012), while 40-59 and 60-79 were the most connected, with 0.04 per day (95%CrI: 0.007-0.15). Estimates were positively associated with the independently obtained social contact rates (Figure 4H, p value = 0.02, Figure S16), but not with sampling intensity (Figure S17).

### Beyond SARS-CoV-2: measurable patterns of spread of respiratory viruses across age groups

To test whether the age-group signal captured by our framework extends beyond SARS-CoV-2, we applied our approach to four other respiratory viruses surveilled by the NRC of Respiratory Viruses in Lyon, France: H1N1pdm, H3N2, RSV-A, and RSV-B. We used the same age groups for influenza. For RSV, due to limited data, we used three age groups: 0–19 years, 20–59 years, 60 and above (Figure 5). Across pathogens, we observed marked differences in the strength of measurable age structure. Fitting our model to these pairwise distance distributions, we reconstructed the observed distributions, their ratio across age-group pairs, and estimated the connectivity between age groups for each pathogen (Figure S18).

**Figure 5:**
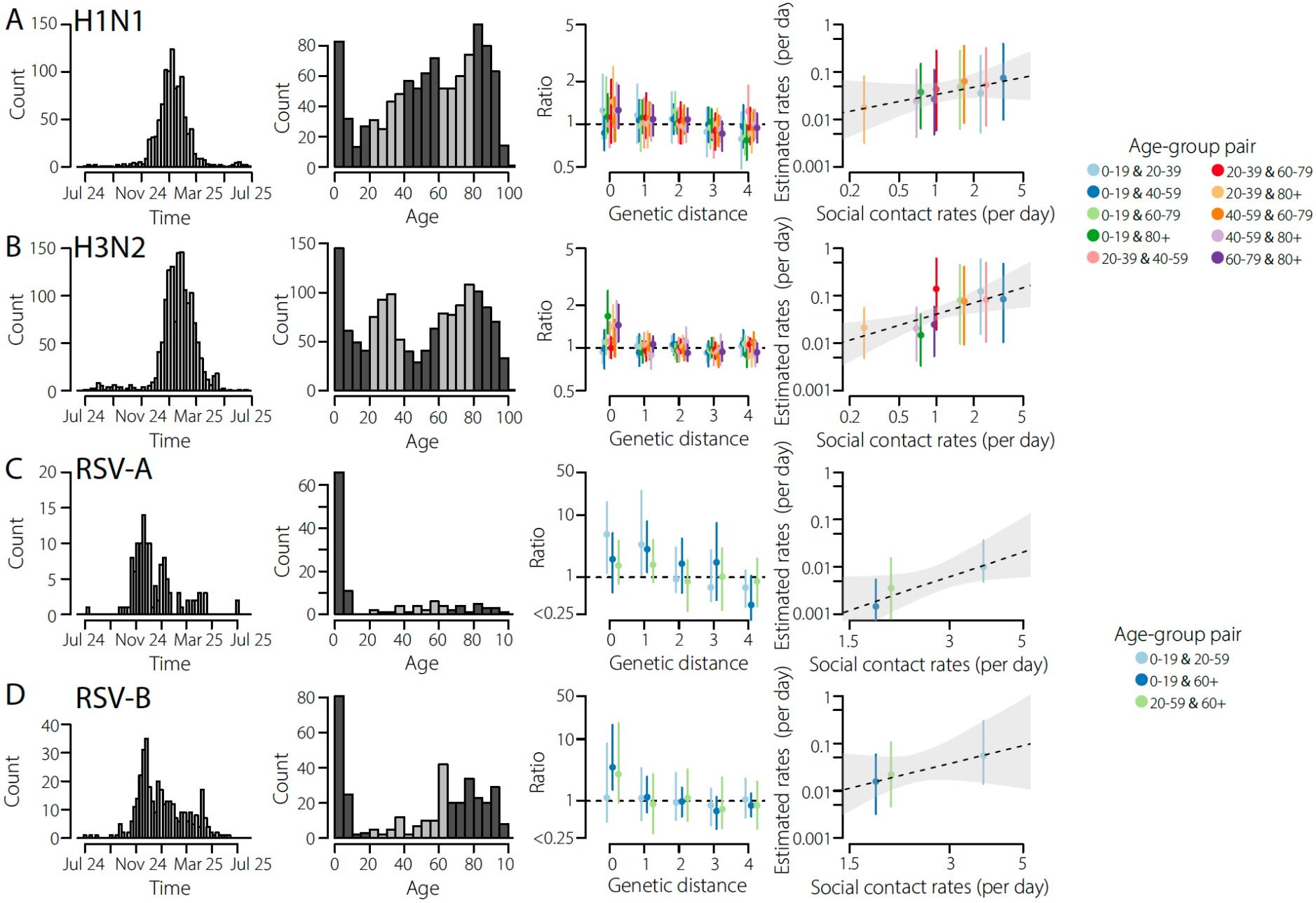
Signal of spread across age groups for H1N1, H3N2, RSV-A and RSV-B. We present signals of age group transmission from sequences of influenza H1N1 (**A**), influenza H3N2 (**B**), RSV-A (**C**) and RSV-B (**D**), from Auverge-Rhône-Alpes region, France. The panels in the first column presents the number of sequences through time, for each pathogen. The second column presents the number of sequences with age information available by age, for each pathogen. Shades of grey indicate the different age groups used. The third column displays the ratio of pairwise genetic distance distributions for sequences from the same age group *vs* different age groups, coloured by age groups. The dashed line denotes a ratio of 1 (i.e. no difference). Dots denote the median, and bars the 95% confidence intervals obtained by bootstrap. The fourth column presents the pairwise comparison between social contact rates (x axis) as estimated by (*28*) and estimated transition rates by our approach (y axis). Estimates were rescaled in units of time using the substitution rate for each pathogen (see Methods). Dots indicate median estimates, and bars indicate 95% credible intervals. The dashed line represents a log-log regression, and the shaded area the confidence region, accounting for the matrix structure. Comparison to sampling intensity per age group is presented in Figure S17. Model fits for each pathogen are present in Figure S18. Phylogenies coloured by age group for each pathogen are presented in Figure S22-23.

While SARS-CoV-2 showed the clearest evidence of age-structured spread (Figure 4D), H1N1pdm and H3N2 displayed weak structure, with nearly all ratio confidence intervals overlapping with one (Figure 5A-B). Nonetheless, inferred H3N2 rates were significantly associated with social contacts (p value = 0.025), while rates for H1N1pdm were not (p value = 0.17, Figure 5A-B). RSV-A and RSV-B showed more heterogeneous patterns and greater uncertainty, likely reflecting the more limited data available (Figure 5C-D). With only three age groups, the permutation test lacked sufficient resolution to assess the association with social contacts. Across all pathogens, our estimated rates were not associated with age-group sampling intensity (Figure S17). Overall, the framework revealed substantial variation in measurable age structure among respiratory viruses and provided a common basis for comparing spread across age groups.

### MERS-CoV spread between camels and humans

Finally, we investigated whether the framework could quantify pathogen spread between hosts in a setting where spread is known to be strongly asymmetric. We applied our framework to a dataset of 178 MERS-CoV genomes collected in the Arabian Peninsula from camels and humans (98 and 80 genomes, respectively) during the 2012–2015 outbreak (*15*). Previous studies showed frequent camel-to-human transmission, but very limited human-to-camel transmission (*7, 15*). Although our model cannot identify the direction of transmission, assuming no human-to-camel transmission allows the estimated overall transition rate to be interpreted as camel-to-human transmission. The resulting rate is consistent with previous estimates (Figure S19). This illustrates the ability of the framework to quantify movement between populations even when the underlying transmission process is strongly asymmetric.

## Discussion

We introduce a framework to quantify pathogen spread directly from pairwise genetic distance distributions within and between states. It addresses a gap between full phylogenetic methods, which can reconstruct spread in detail but are computationally intensive or sensitive to uneven sampling (*10*), and more scalable pairwise distance-based approaches, which have mostly been used qualitatively (*11, 12, 16, 17*).

Across simulations, the method recovered migration rates across diverse demographic histories and sampling schemes. In empirical datasets, inferred rates were associated with independent measures of human mobility and social contact, supporting their interpretation as biologically meaningful quantitative summaries of transmission. MERS-CoV further showed that connectivity can be quantified even when transmission is strongly asymmetric.

The range of transition rates that can be estimated depends on the genetic resolution of the data, which is shaped by evolution rate, transmission dynamics and sampling intensity (*18*). Notably, the informative timescale varies among pathogens: typically weeks to months for rapidly evolving RNA viruses, but years or longer for more slowly evolving bacteria. Among the respiratory viruses analysed here, SARS-CoV-2 showed the clearest age-group structure, influenza weaker but detectable structure, and RSV more limited or uncertain structure. Although these differences likely reflect variation in the timescales of molecular evolution and transmission, the estimated patterns of connectivity between age groups were broadly consistent across pathogens. This consistency suggests that inferred rates capture meaningful transmission structure and could be used to investigate the contributions of contact patterns, age-specific susceptibility, and transmission intensity to pathogen spread.

A key feature of our framework is that it used the overall pairwise distance distribution to capture demographic and genealogical structure while modelling how migration redistributes this structure across states. This approach avoids explicit modelling of the full population history, as done in structured coalescent models (*6*) or multi-type birth-death models (*7*), while still retaining enough information to estimate transition rates. In practice, this greatly expands the scale of datasets that can be analysed and makes the method attractive for large genomic surveillance. Because inference relies on pairwise genetic structure rather than on raw sampling frequencies, it also remains robust to uneven sampling across states, a common feature of modern pathogen genomic datasets.

## Limitations

Our framework infers transition rates from pairwise genetic distance distributions rather than phylogenetic trees, meaning we ignore part of the phylogenetic information contained in the sequences. This improves scalability but limits identifiability. In particular, migration direction cannot be inferred. The method is therefore best suited to estimating symmetric or effectively averaged connectivity patterns. However, some of this information about asymmetry is preserved in phylogenetic structure and could be further exploited. Furthermore, our framework makes the simplifying assumption that pairs of sequences are independent observations, even though they are linked through the same underlying transmission process. This simplification is necessary to keep inference tractable and should be reasonable in large populations, but may become less accurate in smaller trees or populations.

Relying on these pairwise genetic distance distributions implies that differences in sample numbers across states do not themselves bias inference, however, preferential sampling of particular lineages or outbreaks may distort these distributions and influence estimated rates. Thus, as with any phylogenetic methods quantifying migration rates, datasets need to be carefully assessed regarding preferential sampling biases. Finally, the method assumes that the baseline pairwise distance distribution captures demographic effects independently of migration. This approximation performs well across a broad range of settings, and is expected to hold when migration occurs faster than coalescence, but may break down when the two processes occur on similar timescales and are then coupled.

## Conclusions

Our results demonstrate that pairwise genetic distance distributions provide a robust signal to quantify pathogen spread across states. Our method avoids full phylogenetic reconstruction, remains computationally efficient, and scales to large and unevenly sampled genomic datasets. Across simulated and empirical applications, we show that transition rates can be estimated across spatial, demographic and host scales, establishing pairwise genetic distance distributions as a scalable basis for genomic epidemiological inference.

## Acknowledgements

We gratefully acknowledge all data contributors, *i.e*., the Authors and their Originating laboratories responsible for obtaining the specimens, and their Submitting laboratories for generating the genetic sequence and metadata and sharing via the GISAID Initiative, on which this research is based. We thank the National Reference Center of Respiratory Viruses in Lyon, France for sharing pathogen genomic sequences and useful discussions.

## Authors contributions

Conceptualization: N.L.; Methodology: N.L., R.A.N. and T.S.; Data curation, Visualization: N.L.; Funding acquisition: N.L. and T.S., Writing – original draft: N.L., Writing – review & editing: N.L., R.A.N. and T.S.

## Data, code, and materials availability

The code supporting the analysis presented in the manuscript is available at https://github.com/noemielefrancq/Quantifying_pathogen_spread_pairwise_genetic_distance_distributions. All the sequences isolates used in this study are listed in Tables S1-7, and available on NCBI (see Table S1 and S7) or GISAID(*19*), with EPISET IDS: EPI_SET_261110ge(*20*), EPI_SET_261110qd(*21*), EPI_SET_261110xe(*22*), EPI_SET_261110mh(*23*) and EPI_SET_261110rs(*24*).

## List of Supplementary Materials

### Materials and Methods

Supplementary Text

Supplementary Text S1: Relationship between the co-location approximation and the structured coalescent

Supplementary Text S2: Details on the identifiability of asymmetric migration matrices

### Supplementary Figures

Figure S1: Pairwise distance distributions are robust to subsampling

Figure S2: Examples of the impact of population dynamics on pairwise distance ratios

Figure S3: Examples of model fits across four migration rates

Figure S4: Additional sensitivity analyses under different population dynamics Figure S5: Additional sensitivity analyses under different sampling schemes Figure S6: Comparison of our method with estimates from ace

Figure S7: Fitting symmetric three-state migration matrices Figure S8: Fitting four- and five-state migration matrices Figure S9: Unequal demography in multi-deme systems

Figure S10: Fitting a three-deme matrix in the presence of uneven sampling

Figure S11: Non-identifiability of asymmetric migration and estimation of the average migration rate in a two-deme population

Figure S12: Theoretical signal of asymmetric migration and non-identifiability of individual rates in a three-deme population

Figure S13: Model fits by country pair for SARS-CoV-2 spread between countries

Figure S14: Estimated SARS-CoV-2 transition rates between countries and comparison with covariates Figure S15: Robustness of estimated transition rates between countries to subsampling of the SARS-CoV-2 dataset

Figure S16: Comparison of estimated SARS-CoV-2 transition rates across age groups using different contact matrices

Figure S17: Comparison of estimated transition rates across age groups with sampling intensity for SARS-CoV-2, H1N1, H3N2, RSV-A, and RSV-B

Figure S18: Model fits for spread across age groups in H1N1, H3N2, RSV-A, and RSV-B Figure S19: Estimating the transmission rate between camels and humans in MERS-CoV Figure S20: SARS-CoV-2 phylogeny of sequences from seven countries, coloured by country

Figure S21: SARS-CoV-2 phylogeny of sequences from Auvergne-Rhône-Alpes, France, coloured by age group

Figure S22: Influenza A/H1N1 and H3N2 phylogenies of sequences from Auvergne-Rhône-Alpes, France, coloured by age group

Figure S23: RSV-A and RSV-B phylogenies of sequences from Auvergne-Rhône-Alpes, France, coloured by age group

Figure S24: Approximation versus true P(d | diff) distributions Figure S25: Likelihood profiles across migration rates

### Supplementary tables

Table S1 (csv): Isolates of SARS-CoV-2 for the geographical spread analysis

Table S2 (pdf): EPISET of isolates of SARS-CoV-2 for the age-group transmission analysis Table S3 (pdf): EPISET of isolates of H1N1 influenza for the age-group transmission analysis Table S4 (pdf): EPISET of isolates of H3N2 influenza for the age-group transmission analysis Table S5 (pdf): EPISET of isolates of RSV-A for the age-group transmission analysis

Table S6 (pdf): EPISET of isolates of RSV-B for the age-group transmission analysis Table S7 (csv): Isolates of MERS-CoV

