## Supplementary materials for "Quantifying pathogen spread across scales from pairwise genetic distance distributions"

**Affiliations**

**List of Supplementary Materials.**

Materials and Methods

Supplementary Text
Supplementary Text S1: Relationship between the co-location approximation and the structured coalescent
Supplementary Text S2: Details on the identifiability of asymmetric migration matrices

Supplementary Figures
Figure S1: Pairwise distance distributions are robust to subsampling
Figure S2: Examples of the impact of population dynamics on pairwise distance ratios
Figure S3: Examples of model fits across four migration rates
Figure S4: Additional sensitivity analyses under different population dynamics
Figure S5: Additional sensitivity analyses under different sampling schemes
Figure S6: Comparison of our method with estimates from ace
Figure S7: Fitting symmetric three-state migration matrices
Figure S8: Fitting four- and five-state migration matrices
Figure S9: Unequal demography in multi-deme systems
Figure S10: Fitting a three-deme matrix in the presence of uneven sampling
Figure S11: Non-identifiability of asymmetric migration and estimation of the average migration rate in a two-deme population
Figure S12: Theoretical signal of asymmetric migration and non-identifiability of individual rates in a three-deme population
Figure S13: Model fits by country pair for SARS-CoV-2 spread between countries
Figure S14: Estimated SARS-CoV-2 transition rates between countries and comparison with covariates
Figure S15: Robustness of estimated transition rates between countries to subsampling of the SARS-CoV-2 dataset
Figure S16: Comparison of estimated SARS-CoV-2 transition rates across age groups using different contact matrices
Figure S17: Comparison of estimated transition rates across age groups with sampling intensity for SARS-CoV-2, H1N1, H3N2, RSV-A, and RSV-B

Figure S18: Model fits for spread across age groups in H1N1, H3N2, RSV-A, and RSV-B

Figure S19: Estimating the transmission rate between camels and humans in MERS-CoV

Figure S20: SARS-CoV-2 phylogeny of sequences from seven countries, coloured by country

Figure S21: SARS-CoV-2 phylogeny of sequences from Auvergne-Rhône-Alpes, France, coloured by age group

Figure S22: Influenza A/H1N1 and H3N2 phylogenies of sequences from Auvergne-Rhône-Alpes, France, coloured by age group

Figure S23: RSV-A and RSV-B phylogenies of sequences from Auvergne-Rhône-Alpes, France, coloured by age group

Figure S24: Approximation versus true P(d | diff) distributions

Figure S25: Likelihood profiles across migration rates

Supplementary tables

Table S1 (csv): Isolates of SARS-CoV-2 for the geographical spread analysis

Table S2 (pdf): EPISET of isolates of SARS-CoV-2 for the age-group transmission analysis

Table S3 (pdf): EPISET of isolates of H1N1 influenza for the age-group transmission analysis

Table S4 (pdf): EPISET of isolates of H3N2 influenza for the age-group transmission analysis

Table S5 (pdf): EPISET of isolates of RSV-A for the age-group transmission analysis

Table S6 (pdf): EPISET of isolates of RSV-B for the age-group transmission analysis

Table S7 (csv): Isolates of MERS-CoV

**Materials and methods.**

**Overview**

We developed a method to estimate the rate of pathogen spread between demes directly from these distance distributions. The approach links genetic divergence to backward-in-time movement between demes through a continuous-time Markov chain (CTMC) model, capturing how deme connectivity shapes the observed patterns of pairwise distributions.

**Pairwise distance distributions across demes**

We consider a pathogen population subdivided into $K$ discrete demes (*e.g.* locations, traits). Given a set of sequences sampled across demes $i$ and $j$, we compute the distribution of pairwise genetic distances, stratified according to the sampling demes of each pair: $ii$, $jj$ and $ij$.

Let $d$ denote the pairwise genetic distance between two sequences. We note that here distances are in units of genetic distance, which can be rescaled to time if the substitution rate is known. We compute the distribution of pairwise distances between sequences sampled in a pair of demes $i$ and j as:

$$\widehat{D}(d \mid i,j) = \frac{\widehat{N}(d \mid i,j)}{\widehat{N}(i,j)}$$

[Eqn. 1]

Where $\widehat{N}(d \mid i,j)$ and $\widehat{N}(i,j)$ denote the number of pairs where one sequence is sampled in $i$, and the other one in $j$, either by distance $d$ or irrespective of the distance. In practice, these quantities are estimated using histograms of pairwise distances, which are discretised into $H$ bins.

For a given pair of demes $i$ and $j$, we define the within-deme distance distribution as the average of the two deme-specific distributions:

$$\widehat{D}(d \mid same, i,j) = \frac{1}{2}\widehat{D}(d \mid i,i) + \frac{1}{2}\widehat{D}(d \mid j,j)$$

[Eqn. 2]

This gives equal weight to the distance distribution within each deme, so that $\widehat{D}(d \mid same, i,j)$ represents the average within-deme genetic structure. The between-deme distribution is simply:

$$\widehat{D}(d \mid diff, i,j) = \widehat{D}(d \mid i,j)$$

[Eqn. 3]

As each pair distribution is normalised by its number of sequence pairs, these distributions are not affected by differences in sampling intensity between demes. In Figure S1 we present examples of these distributions in the presence of strong sampling bias and show that their expectation remains unchanged.

To facilitate comparison between the two distributions $\widehat{D}(d \mid same, i,j)$ and $\widehat{D}(d \mid diff, i,j)$, we define the ratio $\widehat{R}_{ij}(d)$:

$$\widehat{R}_{ij}(d) = \frac{\widehat{D}(d \mid same, i,j)}{\widehat{D}(d \mid diff, i,j)}$$

[Eqn. 4]

This quantity measures the relative enrichment of same-deme pairs at a given genetic distance, compared to different-deme pairs. It is helpful to visualise the signal present in the data (*i.e.* difference between $\widehat{D}(d \mid same, i,j)$ and $\widehat{D}(d \mid diff, i,j)$). Values of $\widehat{R}_{ij}(d) > 1$ indicate that pairs at distance $d$ are more likely to be observed within the same demes than between demes, and therefore reflect stronger within-deme clustering. Conversely, values of $\widehat{R}_{ij}(d) \simeq 1$ indicate no observed structure.

Closely related quantities have previously been used to characterise clustering and spatial structure in pathogen genomic data (*11*, *12*).

Confidence intervals for $\hat{R}_{ij}(d)$ are obtained using a bootstrap approach. Sequences are resampled with replacement independently within each deme, while preserving the original number of sequences sampled from each deme, over 200 resampling events, and recalculating the statistic each time. The 95% confidence intervals are given by the 2.5 and 97.5% quantiles from the resultant distribution.

We use $\hat{R}_{ij}(d)$ to visualise the structure in the pairwise distance distributions. Inference is performed below directly on the distance distributions.

**Model-based pairwise distance distributions**

Here we fit model-based $P(d \mid ij, Q)$ to the observed distributions $\hat{D}(d \mid i, j)$ to estimate the migration rates between demes without explicitly modelling the underlying demographic history.

We model migration backwards in time using a continuous-time Markov chain with rate matrix $Q$, where $Q_{ij}$ describes the per-lineage rate of migration from deme $i$ to deme $j$. These rates can be translated to forward-in-time rates, using $Q_{ij} = \frac{\pi_j}{\pi_i} Q_{ji}{}^{forward}$, where $\pi_i$ denotes the share of the population that is in deme $i$. Forward and backward rate matrices are equal for a stationary, reversible lineage-state process, but not in general. Here, we parameterise $Q$ directly in backward time, similarly to structured-coalescent-based models (*29*, *30*).

**Structured-coalescent motivation.** In a structured population, pairwise distance distributions are defined by two interacting processes: migration between states and demography within states. We start by writing how those two processes interact formally, under a structured coalescent model. Let $c_k = \frac{1}{N_{e\,k}}$ be the coalescence rate in deme $k$ and let $q_{kk}(t|\, ij)$ denote the probability that a pair sampled at present in deme $i$ and $j$ has not yet coalesced and that its two ancestral lineages occupy the same deme $k$ at distance $t$ in the past. In the case of a two-deme system, their dynamics are defined by:

$$\frac{d\,q_{ii}}{dt} = -(2Q_{ij} + c_i)q_{ii} + Q_{ji}q_{ij}$$
$$\frac{d\,q_{jj}}{dt} = -(2Q_{ji} + c_j)q_{jj} + Q_{ij}q_{ij}$$
$$\frac{d\,q_{ij}}{dt} = 2Q_{ij}q_{ii} + 2Q_{ji}q_{jj} - (Q_{ij} + Q_{ji})q_{ij}$$

[Eqn. 5]

In general, for any number of demes, the dynamics of the probabilities $q_{kk}(t|\, ij)$ are described by a system of ordinary differential equations parameterized by migration backward-in-time and the coalescence rates (see Supplementary Text S1). From these probabilities, we can write the coalescence time density, which determines the corresponding pairwise evolutionary-distance distribution:

$$P(t \mid ij) = \sum_k c_k\, q_{kk}(t|\, ij)$$

[Eqn. 6]

From the densities, one can in principle identify both the migration and population dynamics parameters. Taking the first-order expansions around $t = 0$, and assuming a constant effective population size, we have:

$$P(t \mid ii) = c_i - c_i t(2\,Q_{ij} + c_i) + o(t)$$
$$P(t \mid jj) = c_j - c_j t(2\,Q_{ji} + c_j) + o(t)$$
$$P(t \mid ij) = t(c_i\,Q_{ji} + c_j Q_{ij}) + o(t)$$

[Eqn. 7]

This illustrates the information contained in the pairwise distance distributions. Under constant effective population sizes and at short distances, the within-deme densities inform the coalescence rates, while their slopes depend on outward backward migration. The between-deme density depends on both migration and coalescence. In particular, taking the ratio between the average within-deme and between-deme densities gives, at short distances:

$$R_{ij}(t) = \frac{P(t \mid ii) + P(t \mid jj)}{2P(t \mid ij)} \approx \frac{c_i + c_j}{2t(\,c_i\,Q_{ji} + c_j Q_{ij})}$$

[Eqn. 8]

For symmetric migration, *i.e.* $Q_{ij} = Q_{ji}$, this simplifies to:

$$R_{ij}(t) \approx \frac{1}{2t\,Q_{ij}}$$

[Eqn. 9]

Thus, the ratios retain information about migration while reducing the dependency on population dynamics (Figure 1D). This provides the intuition for the migration signal present in the data. Our inference focuses on the underlying pairwise genetic distance distributions and models how migration shapes their structure across demes.

Jointly estimating both the migration rates and the population dynamics is challenging, as one needs assumptions about the population dynamics through time. Instead, here, we introduce the simplifying assumption that migration and coalescence can be decoupled, and estimate the migration rates between demes without explicitly modelling the underlying demographic history. Specifically, we use an observed pairwise genetic distance distributions as an empirical reference to account for demographic effects, while estimating migration rates from the relative differences between within- and between-deme distributions.

**Inference model for pairwise genetic distance distributions.** For a pair of demes $i$ and $j$, we model the between-deme pairwise genetic distance distribution using a reference distribution describing the underlying demographic structure of the population together with a migration model describing how this structure is distributed across demes.

We define the reference pairwise distance distribution in the population, $P_{ref,i,j}(d|Q)$, in which demes $i$ and $j$ contribute equally, by:

$$P_{ref,i,j}(d|Q) = \frac{1}{4}P(d \mid i,i,Q) + \frac{1}{4}P(d \mid j,j,Q) + \frac{1}{2}P(d \mid i,j,Q)$$

[Eqn. 10]

This distribution captures the underlying demographic structure of the two demes, while remaining independent of their relative sampling intensity.

We then use the migration model to relate the reference distribution to the distance distribution for pairs sampled from demes $i$ and $j$. We have:

$$P(d \mid ij, Q) = \frac{P(ij \mid d, Q)\,P_{ref,i,j}(d|Q)}{P(ij \mid \,Q)}$$

[Eqn. 11]

Where $P(ij \mid d, Q)$ is the probability that a pair at genetic distance $d$ is sampled from demes $i$ and $j$.

Under the full structured coalescent, this probability depends jointly on migration and on deme-specific coalescence rates (as shown above). In our approximation, we instead assume that the information provided by genetic distance about the sampling demes can be captured by the probability of ancestral co-location under the migration process alone. For two sequences separated by a genetic distance $d$ and sampled at the same time, we approximate their most recent common ancestor lies at a distance $d/2$ of each sequence. By definition, the two ancestral lineages can coalesce only if they occupy the same deme. Let $A_d$ denote the event that the two ancestral lineages are co-located at distance $d/2$. We therefore approximate:

$$P(ij \mid d, Q) \approx P(ij \mid A_d, Q)$$

[Eqn. 12]

This relationship is an approximation as the demographic processes within each deme can impact the distribution of pairs across demes. We analyse this approximation and compare it to a structured coalescent in Supplementary Text S1.

We can re-write this probability as:

$$P(ij \mid A_d, Q) = \frac{P(A_d \mid ij, Q)\, P(ij|Q)}{P(A_d \mid Q)}$$

[Eqn. 13]

We define $cr_{ij}(d,Q)$, the co-location ratio:

$$cr_{ij}(d,Q) = \frac{P(A_d \mid ij, Q)}{P(A_d \mid Q)}$$

[Eqn. 14]

Combining Eqns. 11-14, we finally obtain our key equation:

$$\tilde{P}(d \mid ij, Q) = P_{ref,i,j}(d|Q)\, cr_{ij}(d,Q)$$

[Eqn. 15]

The reference distribution $P_{ref,i,j}(d|Q)$ is unknown. We use instead the empirical distribution $\widehat{D}_{ref,i,j}(d)$ to approximate it:

$$\widehat{D}_{ref,i,j}(d) = \frac{1}{4}\widehat{D}(d \mid i,i) + \frac{1}{4}\widehat{D}(d \mid j,j) + \frac{1}{2}\widehat{D}(d \mid i,j)$$

The remaining terms are obtained from the migration process. Let $T(d) = exp(Qd)$, where $T_{il}(d)$ denote the probability that a lineage sampled in deme $i$ is in deme $l$ after tracing it a genetic distance $d$ backwards in time. Assuming that the two ancestral lineages migrate independently:

$$P(A_d \mid ij, Q) = \sum_{l=1}^{K} T_{i,l}\left(\frac{d}{2}\right) T_{j,l}\left(\frac{d}{2}\right)$$

[Eqn. 16]

Then, $P(A_d \mid Q)$ can be computed by:

$$P(A_d \mid Q) = \sum_{r \in \{ii,jj,ij\}} w_r\, P(A_d \mid r, Q)$$

[Eqn. 17]

Where $w_r$ are the weights of pair type $r$ in the reference population. By construction, these are fixed (Eqn. 10), therefore:

$$P(A_d \mid Q) = \frac{1}{4}P(A_d \mid ii, Q) + \frac{1}{4}P(A_d \mid jj, Q) + \frac{1}{2}P(A_d \mid ij, Q)$$

[Eqn. 18]

Thus, the model uses the observed pairwise genetic distance distribution to account non-parametrically for demographic and genealogical structure, while migration determines how this structure is distributed across demes. We note that here we model migration backwards in time, in units of genetic distance, which can be rescaled to time if the substitution rate is known.

In practice, we normalise $\tilde{P}(d \mid ij, Q)$ to model the observed between-deme pairwise distance distribution $\hat{D}(d \mid diff, i, j)$, as our approximation does not in general sum to one over all distances. The relationship between this approximation and the full structured coalescent is derived in Supplementary Text S1. We show in Figure S24 how our approximation captures the relationship between genetic distance distributions and migration rates across a range of scenarios, including demes of different effective population sizes, with only a small error.

**Model implementation.**

For a population subdivided into $K$ demes, we fit our model to each unordered pair of demes $i, j$. For each pair, we construct the reference distribution $\hat{D}_{ref,i,j}(d)$ from $ii$, $jj$ and $ij$ distance distributions (Eqn. 10), and derive $\tilde{P}(d \mid ij, Q)$ using Eqn. 15. All deme pairs are fitted jointly to estimate the migration rates parameterising $Q$.

Pairwise genetic distances are discretised into $H$ bins, of which we fit the first $H* \leq H$ bins. Indeed, at short distances, between-deme pairs are less abundant relative to the reference distribution, and this carries the information on migration. At larger distances, the between-deme and reference distributions converge, carrying no migration information. We therefore set $H*$ to the last bin before $\hat{R}_{ij}(h) < 1$, with a minimum of five bins. All deme pairs use the same $H*$. The co-location ratios are computed for each bin by midpoint-rule numerical integration over the bin, using $N_{grid}$ evaluation points per bin.

For each pair of demes $(i, j)$, we fit both the shape of the between-deme distance histogram and the total number of between-deme pairs falling within the fitted bins $H*$. We first normalise the model quantities $\tilde{P}(d \mid ij, Q)$ over these bins:

$$D_{model}(h \mid ij, Q) = \frac{\tilde{P}(h \mid ij, Q)}{\sum_{h'=1}^{H*} \tilde{P}(h' \mid ij, Q)}$$

[Eqn. 19]

We model the distribution $\hat{N}(h \mid diff, i, j)$ using a Dirichlet-Multinomial likelihood:

$$(\hat{N}(h \mid diff, i, j))_{h=1,\dots,H*} \sim DirichletMultinomial\left(\sum_{h'=1}^{H*} \hat{N}(h' \mid diff, i, j), (D_{model}(h \mid ij, Q))_{h=1,\dots,H*}, \frac{1-\alpha}{\alpha}\right)$$

[Eqn. 20]

And model the total count with a negative binomial likelihood:

$$\sum_{h'=1}^{H*} \hat{N}(h' \mid diff, i, j) \sim NegBin\left(\hat{N}(diff, i, j) \sum_{h'=1}^{H*} \tilde{P}(h \mid ij, Q), \frac{1-\alpha}{\alpha}\right)$$

[Eqn. 21]

Where $\alpha \in (0,1)$ is the parameter controlling overdispersion relative to a pure multinomial (smaller $\alpha$ corresponds to a larger effective concentration $\frac{1-\alpha}{\alpha}$).

For the one-rate models, we place weakly informative priors on the log migration rate: $log(m) \sim Cauchy(-1,1)$, truncated to $[-15,15]$, to avoid numerical issues. For matrices with multiple rates, we parametrise the rates using a global-local decomposition, to help model convergence. We parametrise the $M$ log migration rates parameterising $Q$ to:

$$log\, m_i = m_{global\,mean} + \epsilon_i\,, \sum_{i=1}^{M} \epsilon_i = 0$$

[Eqn. 22]

With $m_{global\,mean} \sim Cauchy(-1,1)$, truncated to $[-15,15]$, being the global mean log-rate, and $\epsilon_i \sim N(0,\delta)$ the deviations. The deviations are constrained to sum to zero, which removes the additive non-identifiability between $m_{global\,mean}$ and $\epsilon_i$. The dispersion parameter $\delta$ is by default set to 1; smaller values shrink individual rates toward the global mean, which aids convergence when $K$ is large and data are sparse. The dispersion parameter prior is set to $\alpha \sim Beta(1,99)$, which concentrates prior mass near low overdispersion.

The model is implemented in Stan and fitted via the cmdstanr R interface (*31*). Posterior inference was performed with 3 chains of typically 1,000 warm-up and 1,000 sampling iterations each. Some complex models might need more iterations, for example when the number of demes is large. Convergence is assessed using the effective sample sizes and the automatic checks from cmdstanr. To perform model comparison, we used the Watanabe–Akaike information criterion (WAIC) implemented in the loo package (*32*, *33*).

The algorithm directly uses the histograms computed from the data, therefore, its computational cost scales with the number of bins, and not the number of sequences, enabling the analysis of large datasets. Pre-computating the pairwise distance histograms can be challenging for the datasets containing hundreds of thousands of sequences. In this case, subsampling the dataset to thousands of sequences and/or using efficient algorithms (*e.g.* pairsnp (*34*)) help manage this cost.

Examples of likelihood profiling in Figure S25 for different migration rates show the sensitivity of our approach.

**Identifiability of migration matrices**

As described above, we model the probability that the ancestors of sequences sampled from demes $k_i$ and $k_j$ are co-located at a distance $d$ in the past (Eqn. 16):

$$P(A_d \mid i,j,Q) = \sum_{l=1}^{K} T_{i,l}(\frac{d}{2})T_{j,l}(\frac{d}{2})$$

With the transition matrix $T(d) = exp\,(Qd)$ where $Q$ is the backward-in-time rate matrix parameterised by the migration rates $Q_{ij}$ to be estimated, and $d$ the pairwise genetic distance. We denote by $T(d)^{\top}$ the transposed matrix of $T(d)$.

**Symmetric migration matrices**. When migration is symmetric, $Q = Q^{\top}$, and therefore $T(d) = T(d)^{\top}$. The co-location probability can then be written as:

$$P(A_d \mid i,j,Q) = [e^{Qd/2}e^{Qd/2}]_{ij} = [e^{Qd}]_{ij}$$

Therefore, using Eqn. 15, one can fit $\tilde{P}(d \mid ij,Q)$ across deme pairs to determine $Q$. It follows that symmetric migration matrices are structurally identifiable by our model for any number of states.

**Asymmetric migration matrices**. When migration is asymmetric, identifiability is more complex. As shown above, the pairwise-distance histograms identify the product $T(d/2)T(d/2)^\top$, rather the directional transition matrix $T(d)$ itself.

In a two-state system, upon exchanging the labels of the two states, migration is reversed, and the co-location probabilities $P(A_d \mid i, i, Q)$ and $P(A_d \mid j, j, Q)$ are swapped, but $P(A_d \mid i, j, Q)$ remains unchanged. As both within-state probabilities enter the reference distribution with equal weights, the between-state distribution alone cannot identify migration direction. Although asymmetric migration affects the within-state distance distributions, these depend on deme-specific demographic histories, leading to confounding between migration asymmetry and differences in effective population sizes. Identifying directional rates would therefore require modelling the within-state distributions under additional demographic assumptions. Nonetheless, fitting a symmetric model broadly recovers the average migration rate between the two demes (Figure S11, see Supplementary Text S1).

For three or more states, reversing the direction of migration is no longer equivalent to relabelling the states, so the argument above does not apply. However, the pairwise distance distributions still only depend on the product $T(d/2)T(d/2)^\top$, through asymmetry can be lost, as we do not observe intermediary states. To demonstrate that asymmetric migration matrices are not always identifiable, we show that we can construct distinct $Q$ matrices that produce the same pairwise distances distributions. In general, all rate matrices $Q$ can be written as:

$$Q = S + K$$

where $S = \frac{Q + Q^\top}{2}$ is symmetric and $K = \frac{Q - Q^\top}{2}$ is anti-symmetric.

If we impose for $S$ and $K$ to commute, *i.e.* $SK = KS$, then we can write (See Supplementary text S2 for details):

$$T(d/2)T(d/2)^\top = e^{Qd/2}e^{Q^\top d/2} = e^{S\,d}$$

[Eqn. 23]

Thus yielding, under the condition that $S$ and $K$ commute, pairwise distance distributions that do not depend on $K$, but only depend on $S$, which is symmetric. Therefore, we have class of matrices $Q$, defined by:

$$Q = S + K; \qquad \text{where } S \text{ and } K \text{ commute}, \quad S = \frac{Q + Q^\top}{2} \text{ and } K = \frac{Q - Q^\top}{2}$$

that cannot be distinguished from pairwise-distance distributions alone. This shows that asymmetric matrices are not generally identifiable.

Additionally, as the class above is relatively restricted due to the condition that $S$ and $K$ need to commute, we also show numerical examples outside of this class. In Figure S12 we show the case of two asymmetric matrices, not from this class, that produce near identical pairwise distances distributions. Nonetheless, particular three or more states asymmetric matrices may still be identifiable.

**Simulation study**

We simulated phylogenies with ReMaster (*35*) under different models to test whether our framework was able to capture known migration rates between demes. To simplify simulations, we assume that evolutionary time is equal genetic distance.

*Two-deme simulations.* In our base scenario, we simulate under a structured coalescent, between two populations A and B. Each population has an effective population size of 1,000 at present, and has been growing exponentially at a rate of 0.1 per unit of time ($N_{eA}(t) = N_{eB}(t) = 1000 * exp(-0.1t)$). The migration rate between population A and B is set constant in each simulation ($m_{AB} = m_{BA}$). We tested rates ranging between 0.01 and 10 movements per individual per unit of time (10 rates tested in total). For each migration rate, we simulated 100 trees, and sampled 1,000 tips from each population A and B at time 0 (present time). This yielded trees of 2,000 tips each, for a total of 1000 phylogenies for this base scenario.

To explore the influence of different population size dynamics between demes, we simulated phylogenies with migration rate between demes of 0.1, 1 and 10 movements per individual per unit of time, and the following population dynamics:

- *Exponential:* We set the effective population size of A to $N_{eA}(t) = 1000 * exp(-0.1t)$, and set the effective population size of B to $N_{eB}(t) = 1000 * exp(-r_B t)$, with $r_B$ ranging from 0.01 to 1 per unit of time (Figure 2A).
- *Constant:* We set the effective population size of A to 1000, and set the effective population size of B to 100, 500, 1000, 5000 or 10000, in each scenario (Figure S4A).
- *Seasonal, varying amplitude:* We set the effective population size of A and B to $N_{eA}(t) = N_{eB}(t) = 1000\,(1 + a\ cos(2\pi t\ /\ P + \phi))$, where $a$ is the amplitude of the seasonality, $P$ is the period of the seasonality and $\phi$ is an offset. We keep the period $P$ to 20 units of time, the offset $\phi$ to 0, and vary $a$ to values ranging from 0.05 to 0.9 (Figure 2B). For example, an amplitude of 0.05 produces a population size that varies between 950 and 1050, while an amplitude of 0.9 produces a population size that varies between 100 and 1900.
- *Seasonal, out of time:* Similarly to above, we set the effective population size of A and B to be seasonal, but with $\phi_A = 0$ and $\phi_B = \pi$, yielding out of time effective population sizes. We vary $a$ to values ranging from 0.05 to 0.9 (Figure S4B).
- *Seasonal, varying period:* Similarly to above, we set the effective population size of A and B to be seasonal. We keep the amplitude $a$ to 0.75, the offset $\phi$ to 0, and vary the period $P$ to values ranging from 1 to 50 units of time (Figure S4C).
- *Seasonal and constant:* Similarly to above, we set the effective population size of A to $N_{eA}(t) = 1000$. For B, we set its effective population size to be seasonal, with varying amplitudes $a$ ranging from 0.05 to 0.9 (Figure S4D).

For each scenario, we simulate 10 trees, and sample 1,000 tips from each population A and B at time 0 (present time).

To explore the influence of different sampling schemes, we simulated phylogenies with exponentially growing effective population sizes $N_{eA}(t) = N_{eB}(t) = 1000 * exp(-0.1t)$, similarly to the base scenario. We use migration rates between demes of 0.1, 1 and 10 movements per individual per unit of time. Compared to the base scenario, where all tips are sampled at present, we explore the following sampling schemes:

- *Sampling at one time point:* we sample each population A and B equally at time 0 (present), with either 50, 100, 250, 500, or 2000 tips (Figure 2C).
- *Sampling at discrete time points:* we sample each population A and B equally every 10 unit of time for 60 units of time, with either 25, 50, 100, 150, or 200 tips each time (Figure S5A).
- *Constant number through time:* we sample each population A and B in the 60 last time units at a constant rate through time, irrespective of their effective population size. We set the expected sampling rates to values ranging from 1 to 20 per unit of time (Figure S5B).

- *Constant proportion through time:* we sample each population A and B in the 60 last time units at a rate proportional to their population size. We sample each population A and B at expected rates of 1 to 20% of their population size per unit of time (Figure S5C).
- Constant proportion through time, biased between demes: as above, we sample each population A and B at a rate proportional to their population size. However, here we sample each population with at different intensities. We set the expected sampling rate of population A to 20%, and the expected sampling rate of population B to of 1-20% of its population size, per unit of time (Figure 2D).

For each scenario, we simulated 10 trees.

On the biased-sampling scenarios above, we compare the estimations of our approach to that of a commonly used model: *ace*, from the package *ape* in R (*36*) (Figure S6). We used *ace* with the default options, and compared the migration rates estimated by both approaches.

To verify that our model was also suitable to capture backward-in-time migration rates in forward-in-time simulations, we simulated phylogenies under a multi-type birth-death model. We simulate a process starting from one individual from one population (randomly chosen from either state A or B). The forward-in-time rate of movement between demes per unit of time was set to 0.1, 1 or 10. The birth rate was set to 0.6 and death rate to 0.5 per unit of time. We set the sampling rate of individuals in population A to 0.5, and the sampling rate of individuals in population B to values ranging from 0.025 to 0.5, without removal. For each scenario, we simulated 10 trees (Figure S5D). To compare our model backward-in-time migration rates with the forward-in-time rates used in these simulations, we assume that the two demes represent equal shares of the population through time, *i.e.* ($\pi_A(t) = \pi_B(t)$). Under symmetric migration, the corresponding forward- and backward-in-time migration rates are therefore equal.

To test whether our framework can recover directional migration, we simulated asymmetric two-deme scenarios. We used the same exponential-growth setup as the base scenario ($N_{eA}(t) = N_{eB}(t) = 1000 * exp(-0.1t)$), and kept the total migration rate constant ($m_{AB} + m_{BA} = 0.1$) while varying its asymmetry. We considered three migration matrices: symmetric ($m_{AB} = m_{BA} = 0.05$ ), mildly asymmetric ($m_{AB} = 0.075, m_{BA} = 0.025$), and strongly asymmetric, with migration strictly one-way ($m_{AB} = 0.1$, $m_{BA} = 0$), with lineages moving from A to B backwards in time. We sampled 1,000 tips from each population at time 0 (present) and simulated 10 trees per condition. We fitted each dataset with the equal-rate (ER) and all-rates-different (ARD) parameterisations of the rate matrix (Figure S11).

*Multi-deme simulations.* Finally, we tested our framework on simulated data with a higher number of demes. We extended the base scenario to three demes A, B and C, each with an effective population size of 1,000 at present growing exponentially at rate 0.1 per unit of time ($N_{eA}(t) = N_{eB}(t) = N_{eC}(t) = 1000 * exp(-0.1t)$). For each condition, we sampled 1,000 tips from each population at time 0 (3,000 tips per tree) and simulated 10 trees. We fitted each dataset with the equal-rate (ER), symmetric (SYM) and all-rates-different (ARD) parameterisations of the rate matrix, comparing them by WAIC. We considered three symmetric migration matrices:

- a one-rate matrix, with equal pairwise rates ($m_{AB} = m_{AC} = m_{BC} = m$), with $m$ ranging from 0.01 to 10 movements per individual per unit of time (10 rates tested, Figure S7A).

- a two-rate matrix, with $m_{AB} = m_{AC}$ held fixed at 0.1 while $m_{BC}$ ranged from 0.01 to 10 (Figure S7B).
- a three-rate matrix, with all three pairwise rates different but symmetric, differing by 2-, 4- or 10-fold ($m_{AB} = 0.1$, $m_{AC} = 0.1/\{2,4,10\}$ and $m_{BC} = 0.1 \times \{2,4,10\}$) (Figure S7C).

We further extended the simulations to four and five demes, with the same population dynamics and sampling (1,000 tips per deme at time 0, 10 trees). We used symmetric matrices with distinct pairwise rates spanning 100-fold: 0.002, 0.01, 0.02, 0.04, 0.1 and 0.2 for four demes, and ten rates between 0.002 and 0.2 for five demes (Figure S8).
To assess the impact of unequal demography with more than two demes, we simulated four demes with effective population sizes of 10,000 at present, growing exponentially at rate 0.1 per unit of time, and six pairwise rates (0.002, 0.01, 0.02, 0.05, 0.1 and 0.2). The rates were either distributed across pairs (non-hub), or assigned so that deme C had the three highest rates (hub). In each case, the effective population size of C was equal to, ten times larger than (100,000) or ten times smaller than (1,000) that of the other demes (Figure S9).

To assess the impact of unequal sampling with more than two demes, we used the three-rate matrices above and sampled 1,000 tips from A and B, and 0 (unsampled), 50, 100, 250, 500 or 1,000 tips from C, simulating 10 trees per condition (Figure S10).

**Geographical spread of SARS–CoV-2 between countries**
**Genetic data.** We selected SARS-CoV-2 genomes available from NCBI, sampled from July 1, 2024, to June 30, 2025, with a complete collection date, isolated in humans, and a minimum length of 29,000 base pairs (equivalent to 97% of the Wuhan-Hu-1/2019 genome length). We downloaded the dataset on April 7, 2026. This yielded 119,999 genomes from 23 countries. Nextclade was used to align all the sequences, in batches of 5,000 sequences, to the Wuhan-Hu-1/2019 genome (*37*). We then selected viruses from countries with at least 100 sequences with good QC scores: Denmark, Japan, Slovakia, Switzerland, Thailand, the UK and the US. For the US and the UK, which had 12,270 and 80,431 sequences, respectively, we randomly selected 5,000 each. This yielded a final dataset of 15,991 sequences, from 7 countries (Denmark: 4895, Japan: 374, Slovakia: 150, Switzerland: 420, Thailand: 152, UK: 5000, US: 5000). The complete list of isolates used and their accession numbers are available in Table S1.

We used IQ-Tree3 (*38*) to reconstruct a maximum likelihood genetic distance-based phylogeny, with the *--pathogen-force* option, which selected CMAPLE as the search algorithm (optimized for highly similar sequences). We used the default DNA substitution model (GTR model). We computed the pairwise distance matrix between all viruses using the *cophenetic.phylo* function in *ape,* rescaled by the SARS-CoV-2 reference genome length (29,903) and saved the output as a symmetric sparse matrix, discarding pairwise genetic distances greater than 20 substitutions.
**Model.** We used our framework to estimate the symmetric migration rate matrix between countries. We computed the histograms on distances ranging from 0 to 15, in steps of 1, for pairs of sequences sampled within 1 month from each other (*i.e.* $\sim$4 generation intervals), which we stratified by pair of country. The model was fitted to the histograms computed from the full dataset. We ran our model using 3 chains of 2,000 warm-up and 2,000 sampling iterations each. We rescaled the migration rates obtained in units of time using a substitution rate of $5.10^{-4}\ subs/site/year$, which matches the within variant mutation rate relevant for comparisons of sequences at short distances (*26*, *27*).

**Mobility data.** We used the Global Transnational Mobility Dataset 2.0 (*25*) from 2022 as a proxy for mobility between countries, normalised by the population size of each country, extracted from the World Bank website (*39*). This matrix $T$ gives the per-capita number of cross-border trips $T_{ij}$ a person in country $i$ makes to a country $j$ per year. These rates are asymmetric by definition. To compare with backward-in-time symmetric migration estimates from our method, we transformed the matrix to a symmetric one by replacing the raw rates by:

$$T_{ij}{}^{sym} = T_{ji}{}^{sym} = \frac{N_i T_{ij} + N_j T_{ji}}{N_i + N_j}$$

Where $N_i$ is the population size of country $i$.

**Transmission of respiratory viruses between age groups**

**Genetic data.** We selected SARS-CoV-2, H1N1pdm, H3N2, RSV–A and RSV-B genomes from the National Reference Center of Respiratory Viruses in Lyon, France, with a complete collection date from July 1, 2024, to June 30, 2025, age-group information was available and from the Auverge-Rhône-Alpes region in France. For SARS-CoV-2, RSV-A and RSV-B we used whole genomes, while for H1N1pdm and H3N2 used the Hemagglutinin (HA) gene. We downloaded the datasets on February 25, 2026. This yielded 1228, 1035, 1428, 120 and 360 sequences for each SARS-CoV-2, H1N1pdm, H3N2, RSV–A and RSV-B, respectively. Similarly to above, Nextclade was used to align all the sequences to the corresponding reference genomes (*37*). For each dataset, we used IQ-Tree2 (*40*) to reconstruct a maximum likelihood genetic distance-based phylogeny, using the *--fast* tree search option, with a GTR+I+G substitution model. For each pathogen, we computed the pairwise distance matrix between all viruses using the *cophenetic.phylo* function in *ape*, rescaled by the sequence length of each virus (SARS-CoV-2: 29,903 base pairs (bp); H1N1pdm HA: 1752 bp; H3N2 HA: 1718 bp; RSV-A: 15225 bp; RSV-B: 15222 bp). The complete list per pathogen of isolates used and their accession numbers are available in Table S2-6.

**Model.** We used our pipeline to estimate the symmetric migration rate matrix between age groups for each virus. For SARS-CoV-2 and Influenza, we used 5 age groups: 0–19 years, 20–39 years, 40–59 years, 60–79 years and 80 years and above. For RSV-A and RSV-B, we used 3 age groups: 0–19 years, 20–59 years, 60 years and above. These age groups were chosen to maximise power given the data available. We computed the histograms on distances ranging from 0 to 5, in steps of 1, for pairs of sequences sampled within a small time window from each other, which we stratified by pairs of age groups. For the time windows, we used 1 month, 12 days, 12 days, 1 month and 1 month for SARS-CoV-2, H1N1pdm, H3N2, RSV-A and RSV-B, respectively (*i.e.* ~4 generation intervals for each pathogen). We ran our model on each pathogen datasets, on the histograms computed from the full dataset, using 3 chains of 2,000 warm-up and 2,000 sampling iterations each. We rescaled the transition rates obtained in units of time using each pathogen substitution rate (SARS-CoV-2: $5.10^{-4}\ subs/site/year$ (*26*, *27*).; H1N1pdm HA: $3.3.10^{-3}\ subs/site/year$ (*41*); H3N2 HA: $4.10^{-3}\ subs/site/year$ (*42*); RSV-A: $6.7.10^{-4}\ subs/site/year$ (*43*); RSV-B: $7.7.10^{-4}\ subs/site/year$ (*43*).

**Social contact rates.** We used social contact data from the SocialCov study, which has been performed during the pandemic (*28*). We used the matrix generated with data from May 2022, as it is the closest in time with our study period (2024-2025). We also used social contact data from the COMES-F study, published in 2015 (*44*), a large population-based contact survey in France. For the SocialCov matrix, we align the age group stratifications to our by distributing the contact of the SocialCov last age group (70+) between our 60-79 and 80+ groups, weighted by population size. For the COMES-F matrix, we used the R package *socialmixr* to compute the contact matrix for each of our age group stratifications. Each matrix $C$describes the per-capita daily number $C_{ij}$ of contacts for an individual of age i with someone of age j. Similarly as above, these rates are by default asymmetric. To compare with backward-in-time symmetric migration estimates from our method, we transformed the matrix to a symmetric one by replacing the raw rates by:

$$C_{ij}^{\ sym} = C_{ji}^{\ sym} = \frac{N_i C_{ij} + N_j C_{ji}}{N_i + N_j}$$

Where $N_i$ is the population size of age group $i$, extract from INSEE (*45*).

**Relationship between estimates and external data.** We fitted a linear regression of the log transition rates versus log-transformed external data (*i.e.* mobility, geographic distance or contact data) across all state pairs, for each posterior draw. We present the resulting regression median and 95% credible intervals across posterior draws. For asymmetric matrices, both directional entries were paired with the corresponding symmetric migration rate. As rates are not independent due to the matrix structure of the data, we assessed significance using a permutation test. We permuted the labels of the external matrix in all possible ways and calculated the median slope across 500 posterior draws for each permutation. The p value was the proportion of permutations with a median slope at least as large as observed (or at least as small for expected negative associations). We also tested associations with the total number of sequences sampled from each pair of states. For RSV, three age groups enabled only six possible label permutations, giving a minimum p value of 1/6 and therefore insufficient resolution to assess significance.

**MERS-CoV spread between hosts**

**Data.** We reanalysed a dataset comprised of an alignment of 274 MERS-CoV genomes, including 100 genomes from camel hosts and 174 from humans (*15*). We restricted the analysis to the time period during which both humans and camels were sampled, *i.e.* from 2013-11-07 to 2015-05-28. This yielded 178 genomes, listed in Table S7. We extracted evolutionary distances (in units of time) between isolates from the trees generated in a BDMM BEAST2 analysis (*7*).

**Model.** We used our model to estimate the transition matrix between camel and human hosts. As the data was sparse, we computed the histograms on distances ranging from 0 to 4 years, with bins every 0.1 year up to a distance of 1 year, and then bins of 0.5 years, for pairs of sequences sampled within 0.1 year from each other.

As our model does not capture directional transmission (Figure S11), we fitted the equal-rate model, which estimates the average of the two directional rate (see Supplementary text S1). We ran 3 chains of 2,000 warm-up and 2,000 sampling iterations each. To represent uncertainty in the observed ratios, we computed 200 bootstrap replicates of the histograms. Previous studies identified frequent camel-to-human transmission but little evidence of sustained human-to-camel transmission (*7*, *15*). Assuming no human-to-camel transmission, the symmetric rate estimated by our model corresponds to half of the backward migration rate from humans to camels, and we applied the same transformation to its prior. All rates are backward-in-time rates per lineage: the rate at which a lineage sampled in humans moves to camels corresponds to camel-to-human movement forward in time, but converting the backward rate into forward-in-time rate requires additional information about the relative sizes of the two host populations.

**Comparison with BDMM-prime.** We compared our estimates with the rate implied by the BDMM-prime analysis of the same dataset (*7*). The analysis was parameterised for each host by a reproduction

number ($R_c$, $R_h$) and a become-uninfectious rate ($\delta_c$, $\delta_h$), and by among-host reproduction numbers ($R_{ch}$, $R_{hc}$). $I_c$ and $I_h$ denote the number of infected individuals from each host. As BDMM-prime estimates forward-in-time rates, we converted them to the equivalent backward-in-time rate at which a lineage in humans moves to camels: $m_{h>c} = \frac{R_{ch}\delta_c I_c}{I_h}$.
(*46*). The resulting estimates were broadly comparable to those obtained with our framework (Figure S18C).

**Supplementary Text S1: Relationship between the co-location approximation and the structured coalescent**

In the methods, we approximate in Eqn 12:

$$P(ij \mid d, Q) \approx P(ij \mid A_d, Q)$$

where $A_d$ denotes the event that the two ancestral lineages are co-located at distance $d/2$ from the sampled sequences. This approximation separates the migration process on ancestral lineage locations from the demographic process driving coalescence.

Under the structured coalescent, migration and coalescence are coupled: ancestral lineages migrate between demes backwards in time, but can coalesce only when they occupy the same deme. Consequently, coalescence can itself alter the distribution of ancestral lineage locations among pairs that remain uncoalesced. Related structured-coalescent approximations similarly model ancestral lineage locations and coalescence conditional on deme membership (*29*, *47*).

In the general case of $K$ demes, let $q_{kk}(t|r)$ denote the probability that a pair sampled in configuration $r$ has not yet coalesced and that its two ancestral lineages occupy the same deme $k$ at distance $t$. Similarly, for $i \neq j$, let $q_{ij}(t|r)$ denote the probability that the pair has not yet coalesced and that its ancestral lineages occupy different demes $i$ and $j$. Let $Q$ denote the backward-in-time migration rate matrix, with $Q_{ij}$ the per-lineage rate of migration from deme $i$ to deme $j$, and let $c_k(t)$ denote the coalescence rate in deme $k$. In the general case, we have:

$$\frac{d\,q_{kk}}{dt} = (2Q_{kk} - c_k(t))q_{kk} + \sum_{l \neq k} Q_{lk} q_{kl}$$
$$\frac{d\,q_{ij}}{dt} = 2Q_{ij}q_{ii} + 2Q_{ji}q_{jj} + (Q_{ii} + Q_{jj})q_{ij} + \sum_{l \notin \{i,j\}} [Q_{lj}q_{il} + Q_{li}q_{jl}]$$

[Eqn. S1]

In this supplementary text, we derive $P(ij \mid d, m)$ under different structured coalescent models and compare it with the migration-only quantity $P(ij \mid A_d, m)$. We first consider a two-deme model with symmetric migration and equal effective population sizes, for which an analytical solution can be obtained. We then consider the case of unequal effective population sizes in two-deme and three-deme models. Finally, we explore a two-deme model with asymmetric migration and equal effective population sizes.

We investigate these models in the supplementary text:

1. Two-deme model with symmetric migration and equal effective population size
2. Two-deme model with different effective population sizes
3. Three-deme model with different effective population sizes
4. Asymmetric migration in a two-deme model

### 1. Two-deme model with symmetric migration and equal effective population size

We first consider two demes $i$ and $j$ with equal coalescence rate $c$. We assume symmetric migration: each ancestral lineage moves from one deme to the other backwards in time at rate $Q_{ij} = Q_{ji} = m/2$, and therefore the total migration rate between deme is $m$.

For simplicity, we measure time in the same units as branch length, such that two contemporaneously sampled sequences with a most recent common ancestor $t$ have pairwise genetic distance $d = 2t$.

Here, the sampling configurations are $r \in \{same, diff\}$, denoting whether the two sequences were sampled from the same demes ($ii$ and $jj$) or different demes ($ij$).

Therefore, simplifying Eqn. S1, we have the system:

$$\frac{d\,q_S}{dt} = -(m+c)q_S + mq_D$$
$$\frac{d\,q_D}{dt} = mq_S - mq_D$$

[Eqn. S2]

For a pair sampled from the same deme, *i.e.* both in $i$ or both in $j$:

$$q_S(t = 0|\,same) = 1, \quad \text{and} \quad q_D(t = 0|\,same) = 0$$

[Eqn. S3]

And for a pair sampled from different demes, *i.e.* one in $i$ and one in $j$:

$$q_S(t = 0|\,diff) = 0, \quad \text{and} \quad q_D(t = 0|\,diff) = 1$$

[Eqn. S4]

**Analytical solution**

We first solve the system in Eqn. S2 for $q_S(t)$, which we can write as:

$$\frac{d}{dt}\begin{pmatrix} q_S \\ q_D \end{pmatrix} = \begin{pmatrix} -(m+c) & m \\ m & -m \end{pmatrix}\begin{pmatrix} q_S \\ q_D \end{pmatrix}$$

[Eqn. S5]

The eigenvalues of $M = \begin{pmatrix} -(m+c) & m \\ m & -m \end{pmatrix}$ are:

$$\lambda_{1,2} = -(m+\frac{c}{2}) \pm \sqrt{m^2+\frac{c^2}{4}}$$

[Eqn. S6]

Defining

$$\delta = \sqrt{m^2+\frac{c^2}{4}}$$

[Eqn. S7]

the general solution is

$$q_S(t) = A\,e^{\lambda_1 t} + B\,e^{\lambda_2 t} \quad \text{and} \quad q_D(t) = A'\,e^{\lambda_1 t} + B'\,e^{\lambda_2 t}$$

[Eqn. S8]

Using the initial conditions in Eqn. S3 and S4 to compute $A$, $B$, $A'$ and $B'$, we obtain:

$$q_S(t|\,same) = e^{-(m+\frac{c}{2})t}\,[cosh(\delta\,t) - \frac{c}{2\delta}\,sinh(\delta\,t)]$$

[Eqn. S9]

and,

$$q_S(t|\,diff) = e^{-(m+\frac{c}{2})t}\,\frac{m}{\delta}\,sinh(\delta\,t)$$

[Eqn. S10]

**Exact distribution of coalescence times and genetic distances**

Coalescence can occur only when the two ancestral lineages occupy the same deme. Conditional on the initial sampling configuration r, the density of coalescence at depth $t$ is therefore:

$$f(t|r) = c\,q_S(t|r)$$

[Eqn. S11]

Because $d = 2t$, the corresponding density of pairwise genetic distances is:

$$g(d|r) = \frac{1}{2}f(\frac{d}{2}|\,r)$$

[Eqn. S12]

Where $g(d|\ same)$ and $g(d|\ diff)$ are the densities of genetic distance conditional on the sampling configuration:

$$g(d|\ same) = P_{SC}(d|same, m, c) \qquad \text{and} \qquad g(d|\ diff) = P_{SC}(d|diff, m, c)$$

[Eqn. S13]

The quantity relevant to the approximation in the main text is the reverse conditional probability,$P_{SC}(diff|d, m, c)$, that is, the probability, under the structured coalescent model, that a pair was sampled from different demes given its genetic distance. By Bayes' rule,

$$P_{SC}(diff|d, m, c) = \frac{P_{SC}(d|diff, m, c)\ P_{SC}(diff|\ m, c)}{P_{SC}(d|\ m, c)}$$

[Eqn. S14]

Using equal reference probabilities for same- and different-deme pair configurations:

$$P_{SC}(same|\ m, c) = P_{SC}(diff|\ m, c) \ = \frac{1}{2}$$

[Eqn. S15]

The marginal distance density is:

$$P_{SC}(d|\ m, c) = \frac{1}{2} g(d|\ same) + \frac{1}{2} g(d|\ diff)$$

[Eqn. S16]

It follows that:

$$P_{SC}(diff|d, m, c) = \frac{g(d|\ diff)}{g(d|\ same) + g(d|\ diff)}$$

[Eqn. S17]

As the coalescence rates are the same in both demes, this simplifies as:

$$P_{SC}(diff|d, m, c) = \frac{q_S(d/2|\ diff)}{q_S(d/2|\ same) + q_S(d/2|\ diff)}$$

[Eqn. S18]

Substituting Eqns. S9 and S10 gives:

$$P_{SC}(diff|d, m, c) = \frac{m\ sinh(\delta\ d/2)}{\delta\ cosh(\delta\ d/2) +\ (m - \frac{c}{2}) sinh(\delta\ d/2)}$$

[Eqn. S19]

And similarly for pairs sampled in the same demes:

$$P_{SC}(same|d, m, c) = \frac{\delta cosh(\delta\ d/2) - \frac{c}{2}\ sinh(\delta\ d/2)}{\delta cosh(\delta\ d/2) + (m - \frac{c}{2})\ sinh(\delta\ d/2)}$$

[Eqn. S20]

Equation S19 therefore gives the exact relationship between genetic distance and sampling configuration under the two-deme structured coalescent.

**Migration-only co-location probability**

The main-text approximation considers the movement of the two ancestral lineages independently of the coalescent process. Let $A_d$ denote the event that the two ancestral lineages are co-located at distance $d/2$.

For two lineages sampled from the same deme, migration alone gives (Eqn. 16 in two demes):

$$P(A_d|same, m) = \frac{1}{2}(1 + e^{-md})$$

For two lineages sampled from different demes,

$$P(A_d|diff, m) = \frac{1}{2}(1 - e^{-md})$$

Under equal reference probabilities for same- and different-deme pairs,

$$P(A_d|\, m) = \frac{1}{2}P(A_d|same, m) \; + \frac{1}{2}P(A_d|diff, m) \; = \frac{1}{2}$$

[Eqn. S21]

Applying Bayes' rule,

$$P(same \mid A_d, m) = \frac{P(A_d \mid same, m)P(same \mid m)}{P(A_d \mid m)}$$

[Eqn. S22]

Because both $P(same \mid m)$ and $P(A_d \mid m)$ equal $\frac{1}{2}$,

$$P(same \mid A_d, m) = \frac{1}{2}(1 + e^{-md})$$

[Eqn. S23]

Similarly,

$$P(diff \mid A_d, m) = \frac{1}{2}(1 - e^{-md})$$

[Eqn. S24]

The approximation in the main text therefore amounts to replacing the exact structured-coalescent quantity $P_{SC}(same|d, m, c)$ with the corresponding migration-only quantity $P(same \mid A_d, m)$.

**Condition of accuracy of the approximation**

Eqn. S19 allows us to determine analytically when the approximation is accurate. Consider the regime in which migration is fast relative to coalescence,

$$\frac{c}{m} \ll 1$$

In this limit,

$$\delta = \sqrt{m^2 + \frac{c^2}{4}} \rightarrow \; m + \frac{c^2}{8m} + o(\frac{c^2}{m})$$

[Eqn. S25]

Therefore,

$$P_{SC}(diff|d, m, c) = \frac{sinh(m\,d/2)}{cosh(m\,d/2) + \; sinh(m\,d/2)}$$

[Eqn. S26]

We obtain:

$$P_{SC}(diff|d, m, c) \approx \frac{1}{2}(1 - e^{-md})$$

[Eqn. S27]

Thus, for pairs of different demes, we have:

$$P_{SC}(ij\,|d, m, c) \approx P(ij \mid A_d, m), \qquad \text{for } \frac{c}{m} \ll 1$$

[Eqn. S28]

This result explains the timescale underlying the approximation. When migration occurs on a faster timescale than coalescence, ancestral lineages can redistribute between demes before coalescence substantially alters their location probabilities. The distribution of ancestral locations is therefore governed predominantly by migration, and the migration-only probability of co-location closely approximates the relationship between genetic distance and sampling configuration generated by the structured coalescent.

Conversely, when coalescence occurs on a timescale comparable to or faster than migration, pairs that become co-located can coalesce before migrating again. Such pairs are removed from the population of surviving ancestral lineages, so conditioning on survival increasingly alters the distribution of ancestral locations. The migration-only co-location probability can therefore deviate from the full structured-coalescent expectation.

### 2. Two-deme model with different effective population sizes

We consider the case in which the two demes have different effective population sizes, and therefore different coalescence rates $c_i$ and $c_j$.

Let $q_{ii}(t|r)$, $q_{ij}(t|r)$ and $q_{jj}(t|r)$ denote the probabilities that a pair sampled in configuration $r$ remains uncoalesced at distance $t$ and has both lineages in deme 1, the lineages in different demes, or both lineages in deme 2, respectively.

For Eqn. S1, the system defining the dynamics is:

$$\frac{d\,q_{ii}}{dt} = -(m + c_i)q_{ii} + \frac{m}{2}q_{ij}$$
$$\frac{d\,q_{jj}}{dt} = -(m + c_j)q_{jj} + \frac{m}{2}q_{ij}$$
$$\frac{d\,q_{ij}}{dt} = mq_{ii} + mq_{jj} - mq_{ij}$$

[Eqn. S29]

For same-deme pairs, using equal reference probabilities for the two demes,

$$q_{ii}(t = 0|\,same) = q_{jj}(t = 0|\,same) = \frac{1}{2}, \quad \text{and} \quad q_{ij}(t = 0|\,same) = 0$$

[Eqn. S30]

For different-deme pairs,

$$q_{ii}(t = 0|\,diff) = q_{jj}(t = 0|\,diff) = 0, \quad \text{and} \quad q_{ij}(t = 0|\,diff) = 1$$

[Eqn. S31]

The exact coalescence-time density is then:

$$f(t|r) = c_i\, q_{ii}(t|r) + c_j\, q_{jj}(t|r)$$

[Eqn. S32]

When $c_1 \neq c_2$, two effects can cause departure from the migration-only approximation. First, previous coalescence alters the distribution of ancestral locations among surviving pairs (same as above). Second, co-location in the two demes does not contribute equally to the probability of coalescence, because a co-located pair coalesces at rate $c_1$ in deme 1 but at rate $c_2$ in deme 2.

Nevertheless, when both coalescence rates are small relative to migration,

$$\frac{max(c_1, c_2)}{m} \ll 1$$

the coalescence terms in Eqns. S29 are small relative to the migration terms. Therefore, the system approaches the migration-only process, and we can write:

$$q_{ii}(t|r) \approx q_{jj}(t|r)$$

[Eqn. S33]

With specifically:

$$q_{ii}(t|\,same) \approx \frac{1+e^{-2mt}}{4} \text{ and } q_{ii}(t|\,diff) \approx \frac{1-e^{-2mt}}{4}$$

[Eqn. S34]

Therefore:

$$f(t|same) \approx (c_i + c_j)\frac{1+e^{-2mt}}{4} \text{ and } f(t|diff) \approx (c_i + c_j)\frac{1-e^{-2mt}}{4}$$

[Eqn. S35]

Replacing in Eqn. S16, the coalescence factors simplify, and we obtain:

$$P_{SC}(diff|d, m, c_i, c_j) = \frac{f(d/2|\,diff)}{f(d/2|\,same) + f(d/2|\,diff)} \approx \frac{1-e^{-md}}{2}$$

[Eqn. S36]

This approximation shows that when $\frac{max(c_1,c_2)}{m} \ll 1$, the probability $P_{SC}(diff|d, m, c_i, c_j)$ becomes independent of coalescence, even when the individual effective population sizes are different.

To compare our approximation with the exact structured-coalescent system, we solve Eqns. S29 numerically and compare the resulting probabilities and pairwise distance distributions across a range of migration rates, effective population sizes, and differences in effective population size between demes (Figure S24). These comparisons quantify the error introduced by treating migration and demography as separable and identify the parameter regimes in which the approximation closely reproduces the full structured-coalescent expectation.

#### 3. Three-deme model with different effective population sizes

We now extend this structured-coalescent system to three demes, $i$, $j$, and $k$, with different effective population sizes. The density of coalescent times for any starting configuration $r$ of the system is, in this case:

$$f(t|r) = c_i\, q_{ii}(t|r) + c_j\, q_{jj}(t|r) + c_k\, q_{kk}(t|r)$$

[Eqn. S37]

As above, when migration dominates coalescence, the probabilities $q_{ll}(t|r)$ can be approximated using the migration-only system. However, unlike the two-deme system, they are not all equal, because they depend on the stating configuration $r$, which by definition can be either $ii$, $jj$ or $ij$, but never $kk$.

Writing again $q_{ll}(t|r)$ in the case where coalescence is negligible compared to migration, we have:

$$q_{ll}(t|ab) \approx T_{a,l}(t)T_{b,l}(t)$$

[Eqn. S38]

Therefore:

$$f(t|ab) \approx \sum_{l\in\{i,j,k\}} c_l\; T_{a,l}(t)T_{b,l}(t)$$

[Eqn. S39]

For pair of demes $i$ and $j$, we have:

$$P_{SC}(diff|d, m, c_i, c_j, c_k) = \frac{f(d/2|ij)}{1/2\, f(d/2|ii) + 1/2\, f(d/2|jj) + f(d/2|ij)}$$

[Eqn. S40]

Which yields:

$$P_{SC}(diff|d,m,c_i,c_j,c_k) \approx \frac{2\sum_{l\in\{i,j,k\}} \; c_l \; T_{i,l}(\frac{d}{2})T_{j,l}(\frac{d}{2})}{\sum_{l\in\{i,j,k\}} \; c_l \; (T_{i,l}(\frac{d}{2}) + T_{j,l}(\frac{d}{2}))^2}$$

[Eqn. S41]

Therefore, in general, the coalescence rates cannot be factored out in a three deme model, and can affect our approximation of $P_{SC}(diff|d,m,c_i,c_j,c_k)$.

**Effect of a highly connected hub.**

Further, we can investigate the particular case when $k$ is a "hub", *i.e.* a deme that strongly connects demes $i$ and $j$. We assume here that:

(1) migration into the hub $k$ is similar from both sampled demes, *i.e.* $T_{i,k}(t) \approx T_{j,k}(t)$

(2) migration is symmetric, *i.e.* $T_{n,l}(t) = T_{l,n}(t)$ for all demes.

(3) equal probabilities of being in the deme of origin for $i$ and $j$, *i.e.* $T_{i,i}(t) \approx T_{j,j}(t)$

In that case, $P_{SC}(diff|d,m,c_i,c_j,c_k)$ in Eqn. S40 simplifies to:

$$P_{SC}(diff|d,m,c_i,c_j,c_k) \approx \frac{1}{2}(1 - \frac{(T_{i,i}(\frac{d}{2}) - T_{i,j}(\frac{d}{2}))^2}{(T_{i,i}(\frac{d}{2}) + T_{i,j}(\frac{d}{2}))^2 + 4\frac{c_k}{(c_i + c_j)}T_{i,k}(\frac{d}{2})^2}$$

[Eqn. S42]

Which shows that in this case, $P_{SC}(diff|d,m,c_i,c_j,c_k)$ is affected by the relative coalescent rate in the hub $k$ compared to the sum of the coalescent rates in the two sampled demes. The distribution is particularly affected when $c_k \gg c_i + c_j$, *i.e.* when the effective population size in deme $k$ is a lot smaller than that of demes $i$ and $j$.

We exemplify this phenomenon in Figure S9. Importantly, we show that if the structure of the migration matrix is not hub-like, the dependence on effective population sizes is minimal.

### 4. Asymmetric migration in a two-deme model

Finally, we explore the effect of asymmetric migration between two demes, with migration rates:

$$a = m_{ij} \text{ and } b = m_{ji}$$

To isolate the effect of this asymmetric migration, we assume initially that the coalescence rates are equal in both demes, *i.e.* $c_i = c_j = c$.

The dynamics are described by:

$$\frac{d\,q_{ii}}{dt} = -(a+c)q_{ii} + \frac{b}{2}q_{ij}$$

$$\frac{d\,q_{jj}}{dt} = -(b+c)q_{jj} + \frac{a}{2}q_{ij}$$

$$\frac{d\,q_{ij}}{dt} = a\,q_{ii} + b\,q_{jj} - (\frac{a+b}{2})q_{ij}$$

[Eqn. S43]

In this case, $q_{ii}(t|r)$ and $q_{jj}(t|r)$ are not generally equal, as migration is asymmetric. However, as coalescent rates are equal, the density of coalescence times simplifies to:

$$f(t|ab) = c(q_{ii}(t|ab) + q_{jj}(t|ab))$$

[Eqn. S44]

We have, under the assumption that coalescence is negligible, similarly as above:

$$q_{ll}(t|ab) \approx T_{a,l}(t)T_{b,l}(t)$$

[Eqn. S45]

Where $T_{a,l}(t)$ can be computed from a CTMC with migration rate matrix:

$$M = \begin{pmatrix} -a/2 & a/2 \\ b/2 & -b/2 \end{pmatrix}$$

[Eqn. S46]

Which gives:

$$T(t) = \frac{1}{a+b} \begin{pmatrix} b + a\, e^{-(a+b)t/2} & a - a\, e^{-(a+b)t/2} \\ b - b\, e^{-(a+b)t/2} & a + b\, e^{-(a+b)t/2} \end{pmatrix}$$

[Eqn. S47]

We define:

$$C_{uv}(t) = \sum_{l} T_{u,l}(t) T_{v,l}(t)$$

[Eqn. S48]

This yields:

$$P_{SC}(diff|d,a,b,c) \approx \frac{C_{ij}(d/2)}{1/2C_{ii}(d/2) + 1/2C_{jj}(d/2) + C_{ij}(d/2)}$$

[Eqn. S49]

Which simplifies to:

$$P_{SC}(diff|d,a,b,c) \approx \frac{1}{2}\left(1 - \frac{e^{-(a+b)d/2}}{1 + (\frac{a-b}{a+b})^2 (1 - e^{-(a+b)d/4})^2}\right)$$

[Eqn. S50]

This shows that $P_{SC}(diff|d,a,b,c)$ predominantly depends on the sum of migration rates $a + b$, and invariant with respect to the direction of migration (*i.e.* exchanging $a$ and $b$ yield the same expression).

If we neglect asymmetry, the expression becomes:

$$P_{SC}(diff|d,a,b,c) \approx \frac{1}{2}(1 - e^{-(a+b)d/2})$$

[Eqn. S51]

Which is the same expression as the symmetry model, with $m = \frac{a+b}{2}$. Therefore, it shows that fitting a symmetric model to data generated under asymmetric migration will predominantly recover the average migration $\frac{a+b}{2}$. This relationship is an approximation as $P_{SC}(diff|d,a,b,c)$ also depends on the magnitude of the asymmetry $\frac{a-b}{a+b}$. Nevertheless, it demonstrates that the dominant information is the sum rather than their individual directional contributions. We exemplify this phenomenon in Figure S11 and S12.

When effective population sizes also differ, additional interactions between migration asymmetry and coalescence rates can arise, further limiting the identifiability of individual directional migration rates.

**Supplementary Text S2: Details on the identifiability of asymmetric migration matrices**

To demonstrate that asymmetric migration matrices are not always identifiable, we show in the method that we can construct distinct $Q$ matrices that produce the same pairwise distances distributions. In general, all rate matrices $Q$ can be written as:

$$Q = S + K$$

where $S = \frac{Q + Q^\top}{2}$ is symmetric and $K = \frac{Q - Q^\top}{2}$ is anti-symmetric.

Here we provide the full derivation of equation 23 from the methods, in the case where we impose for $S$ and $K$ to commute:

$$T(d/2)T(d/2)^\top = e^{Qd/2}e^{Q^\top d/2} = e^{S\,d}$$

Indeed we can write::

$$\begin{aligned} T(d/2)T(d/2)^\top &= e^{Qd/2}e^{Q^\top d/2} \\ &= e^{(S+K)d/2}e^{(S+K)^\top d/2} \\ &= e^{Sd/2+Kd/2}e^{(S^\top+K^\top)d/2} \end{aligned}$$

As $S$ is symmetric, and K is anti-symmetric:

$$T(d/2)T(d/2)^\top = e^{Sd/2+Kd/2}e^{(S-K)d/2}$$

We note that for two commutating matrices $A$ and $B$, we have: $e^{A+B} = e^A e^B$

Therefore we have:

$$\begin{aligned} T(d/2)T(d/2)^\top &= e^{Sd/2}e^{Kd/2}e^{Sd/2}e^{-Kd/2} \\ &= e^{Sd/2}e^{(K+S)d/2}e^{-Kd/2} \\ &= e^{Sd/2}e^{Sd/2}e^{Kd/2}e^{-Kd/2} \end{aligned}$$

And as: $e^{Kd/2}e^{-Kd/2} = I$

Finally we have:

$$T(d/2)T(d/2)^\top = e^{Sd/2}e^{Sd/2}I = e^{S\,d}$$

**Example of matrices unidentifiable**

We take $S$, symmetric matrix, and $K$, anti-symmetric matrix, as:

$$S = \alpha \begin{bmatrix} -2 & 1 & 1 \\ 1 & -2 & 1 \\ 1 & 1 & -2 \end{bmatrix}$$

$$K = \beta \begin{bmatrix} 0 & 1 & -1 \\ -1 & 0 & 1 \\ 1 & -1 & 0 \end{bmatrix}$$

We note that $S$ and $K$ commute. These matrices yield a rate matrix $Q$:

$$Q = S + K = \begin{bmatrix} -2\alpha & \alpha+\beta & \alpha-\beta \\ \alpha-\beta & -2\alpha & \alpha+\beta \\ \alpha+\beta & \alpha-\beta & -2\alpha \end{bmatrix}$$

Which is a valid CTMC rate matrix under the condition if: $|\beta| \leq \alpha$ and $\alpha > 0$, ensuring that all off-diagonal entries are non-negative.

Therefore, directional movements encoded by $\beta$ are not identifiable from pairwise distance distributions: all matrices $Q$ with the same parameter $\alpha$ but different values of $\beta$, with the constraint $|\beta| \leq \alpha$, are indistinguishable under our framework.

**Supplementary Figures**

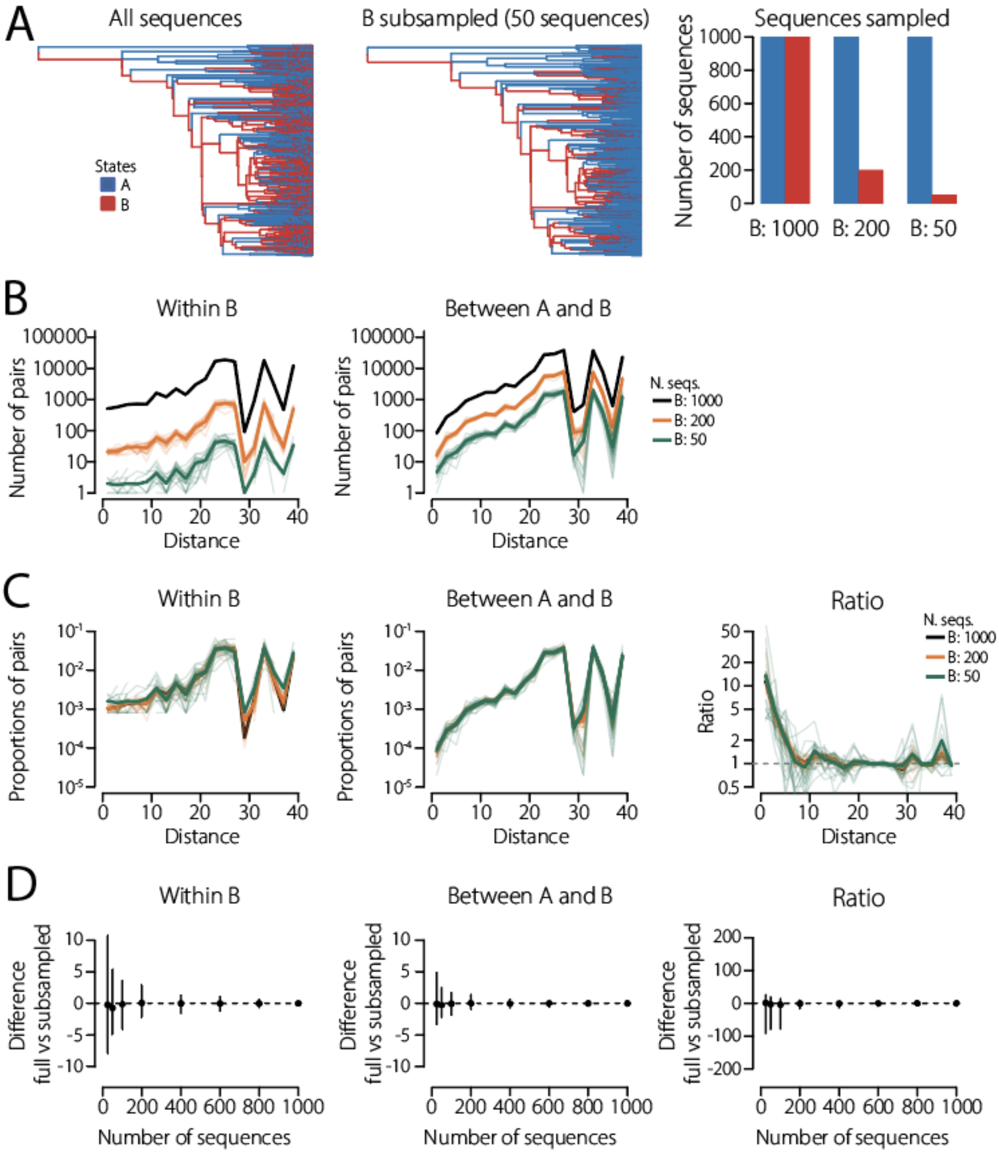


**Figure S1: Pairwise distance distribution are robust to subsampling**

We simulated one phylogeny with two demes A and B, and examined the impact of subsampling the population B on the pairwise distance distributions. **A.** (left) Example simulated phylogeny containing 1000 tips from each population. (center) Example of subsampled phylogeny: from the tree on the left, we subsampled the population B to only keep 50 tips. (right) Details of the number of samples used in the following figures. **B.** Pairwise distances distributions, for pairs of tips sampled both in population B (within B, left) or one in each population (between A and B, right). The distributions shows the raw number of pairs, for each subsampled tree (colours). Shaded lines indicate 20 different random subsamples. Solid lines indicate the median. **C.** Same as B, but we present normalised distributions, *i.e.* they sum up to 1. Additionally, we present the ratio of normalised distributions for each subsample (right). The dashed line denote a ratio of 1. **D.** Difference between the normalized distribution and ratio obtained on the full phylogeny versus the subsampled ones. Dots indicate median differences, bars indicate bars 95% intervals over 200 random subsamples. Dashes line denote no difference. Subsampling changes the number of pairs, but not the distribution of their distances or the ratio, which only become noisier at small sample sizes.

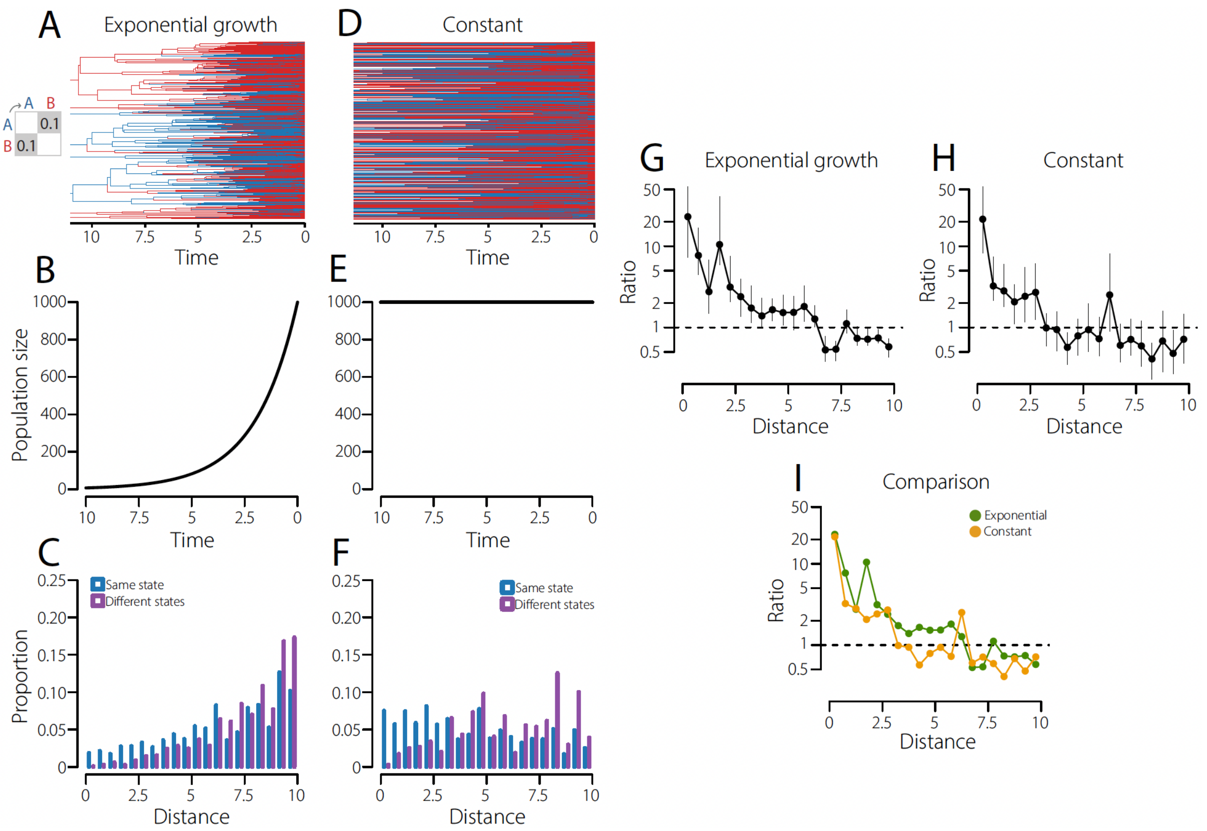


**Figure S2: Examples of the impact of population dynamics on pairwise distance ratios**

Two-state phylogenies were simulated under the same symmetric migration rate ($m = 0.1$) but different population dynamics. **A-C**. Exponentially growing population: simulated phylogeny coloured by state (A), effective population size trajectory (B), and pairwise genetic distance distributions for sequence pairs sampled from the same (blue) or different states (purple) (C). **D–F.** Same as A–C, but for a constant population size. **G and H.** Ratios of the same-state to different-state distance distributions under exponential growth (G) and constant population size (H), computed on all pairs of distances 0 to 10. Points show median ratios and error bars indicate 95% confidence intervals. **I.** Comparison of the ratios obtained under the two population scenarios. Dashed horizontal lines in G-I indicate a ratio of 1. Although the underlying migration process is unchanged across both simulations, variation in population dynamics alters the distributions of pairwise genetic distances, showcasing the need to account for overall transmission history when inferring migration rates from pairwise distance distributions.

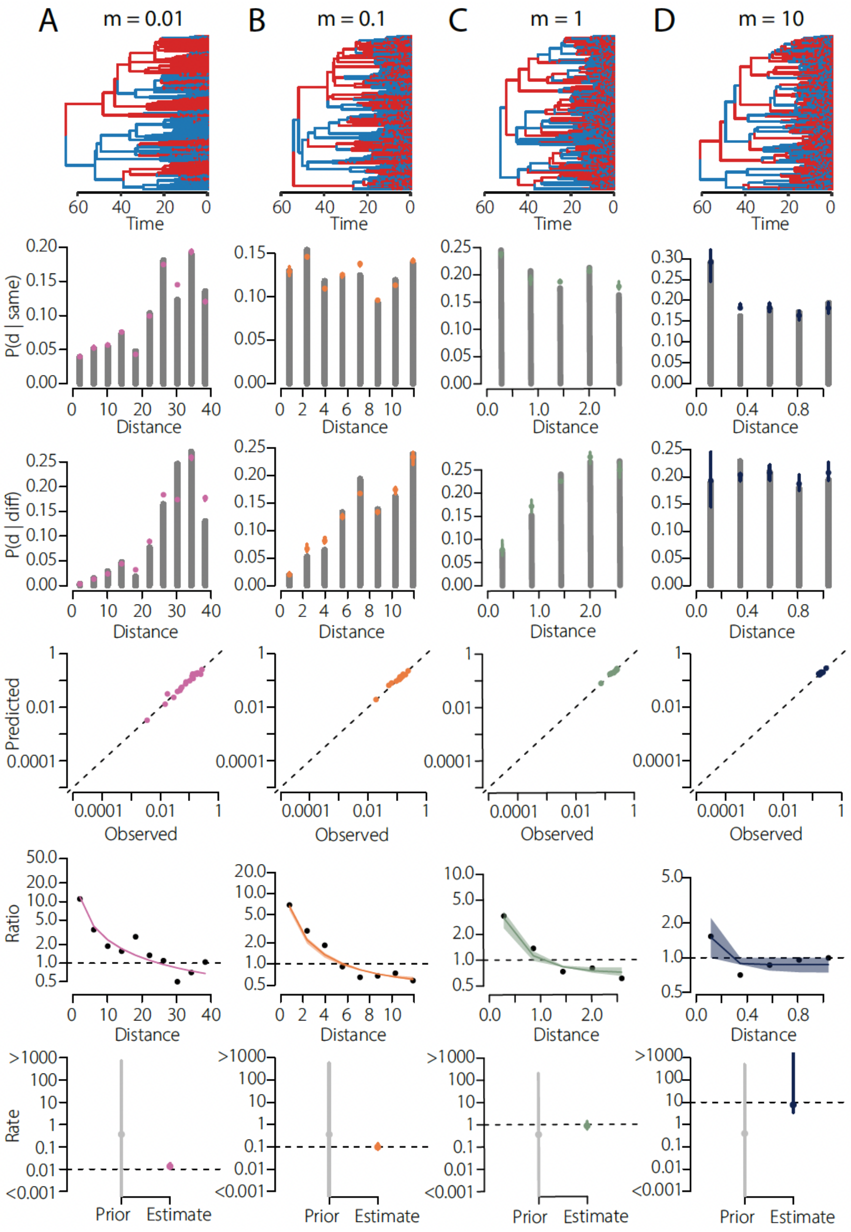


**Figure S3: Examples of model fits across four migration rates**

We show examples of model fits on simulated data generated under four migration rates, 0.01 **(A)**, 0.1 **(B)**, 1 **(C)**, and 10 **(D)**. In each column, the first row shows an example simulated phylogeny, with branches coloured by population (blue: A, red: B). The second and third rows show the observed pairwise genetic distance distributions for sequence pairs sampled within the same state and in different states. Grey bars indicate the observed distributions, and coloured points indicate the corresponding model fits. The fourth row present pairwise comparison of predicted versus observed probabilities for all distance bins. Bars denote the 95% credible intervals. Dashed lines denote identity. The fifth row shows the ratio of the two distance distributions as a function of genetic distance. Black dots denote the observed ratios, the solid coloured line the model fit, and the shaded area the 95% credible interval. The dashed horizontal line indicates a ratio of 1. The sixth row shows the prior and posterior distributions of the migration rate for each simulation. Grey denotes the prior, coloured points denote the posterior median, and bars denote the 95% credible interval. The dashed horizontal line indicates the true migration rate used in the simulation.

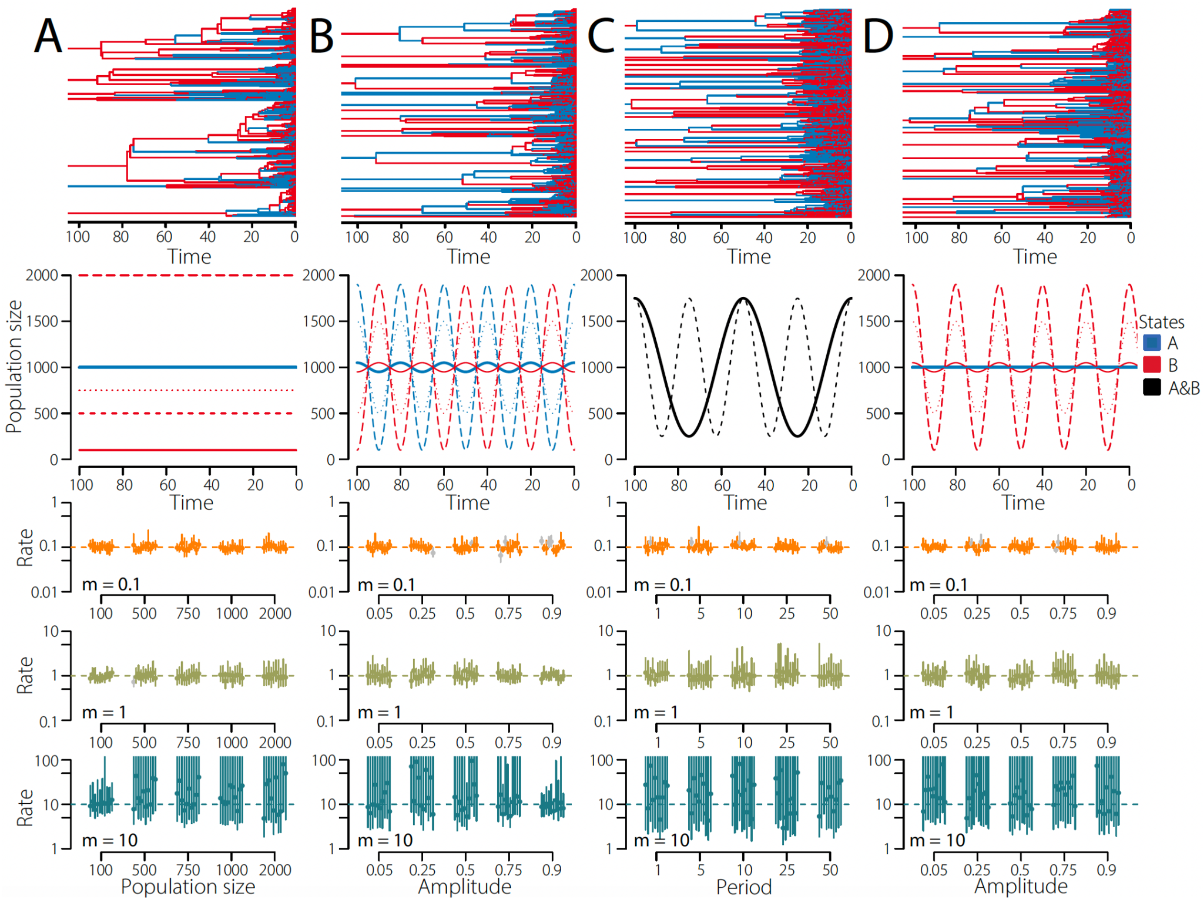


**Figure S4: Additional sensitivity analyses under different population dynamics**

Additional demographic scenarios used to assess robustness of our model estimates to variation in effective population size trajectories. **A.** Constant and unequal effective population sizes between states. **B.** Periodically varying effective population sizes in both states, out of time from each other, with varying amplitudes. **C.** Periodically varying effective population sizes with different periods. **D.** Periodically varying effective population sizes in population B while the population A remains constant. See methods for details on the simulations. The first row shows example of simulated phylogenies, coloured by state: state A (blue), state B (red). The second row shows the effective population size trajectories through time for state A (blue), state B (red), and both states combined (black). For each demographic scenario, the lower three rows show migration rates estimated by our approach for three true migration rates ($m = 0.1$, *1*, and *10*, from top to bottom). Dots indicate median posterior estimates and bars indicate 95% credible intervals. Dashed horizontal lines denote the true migration rates. The x axis indicates the demographic parameter varied across simulations: population size in state B, amplitude of periodic variation, or oscillation period, depending on the panel. Across these additional scenarios, estimates remained broadly unbiased, with wider credible intervals in settings where the phylogenetic signal was weaker. At high migration rates, uncertainty was lowest when population sizes were low (*i.e.* coalescent rates were high), as rapid coalescence generates many closely related sequence pairs that retain short-distance signal of structured transmission. By contrast, higher population sizes allow lineages to be well mixed before coalescing, diluting this signal and increasing uncertainty.

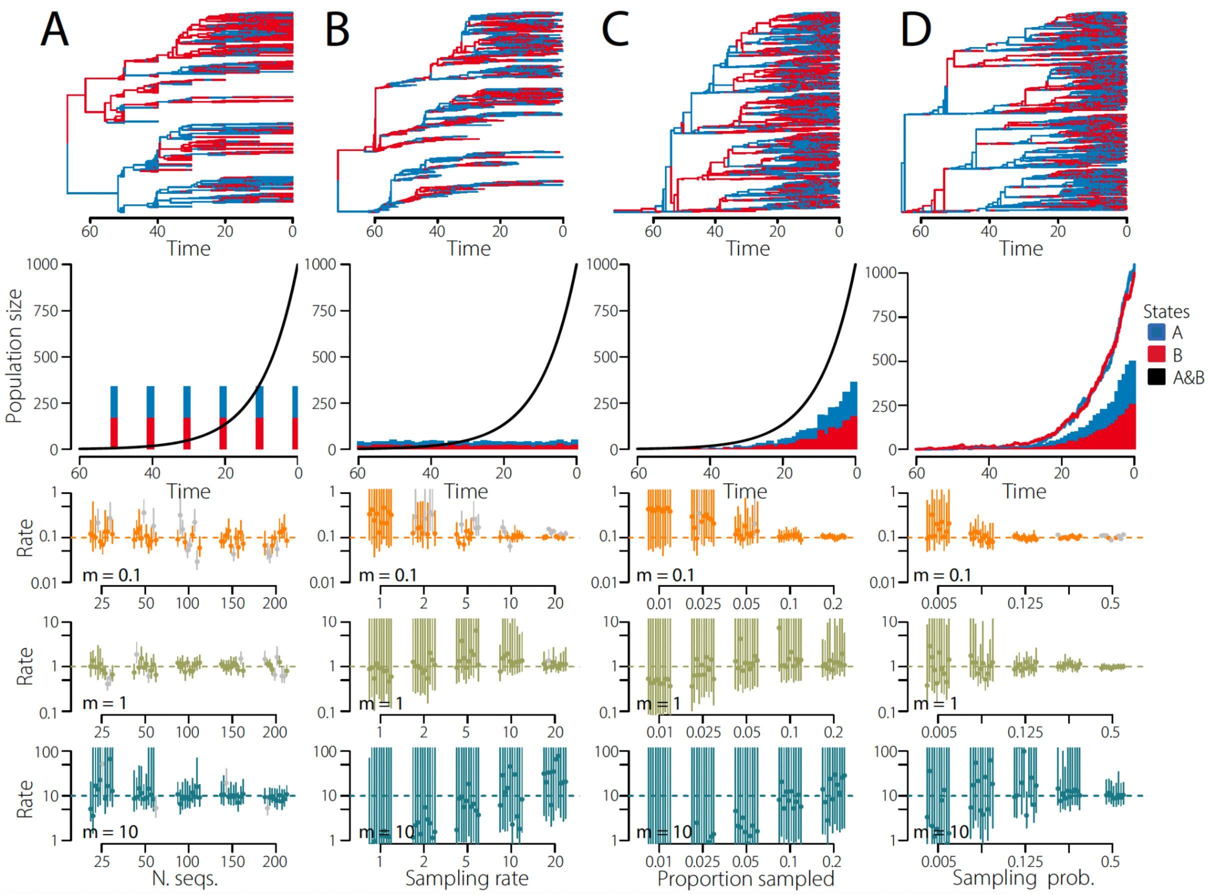


**Figure S5: Additional sensitivity analyses under different sampling schemes**

Additional sampling scenarios used to assess the robustness of our model estimates to variation in sampling schemes. **A.** Sampling at multiple discrete time points, with varying total number of sampled sequences. **B.** Continuous sampling through time at a constant rate, while varying the sampling rate. **C.** Sampling through time at a rate proportional to effective population size, while varying the proportion sampled. **D.** Forward-in-time simulation, using a birth-death model with sampling through time for each state, while varying the sampling probability in one state. In D we present the forward-in-time migration rate estimated, which assumes equal population size in each state. See Methods for details of the simulations. The first row shows examples of simulated phylogenies, coloured by state: state A (blue) and state B (red). The second row shows the effective population size trajectory through time for both states combined (black), together with the corresponding sampled sequences through time by state (blue and red). For each sampling scenario, the lower three rows show migration rates estimated by our approach for three true migration rates ($m = 0.1$, $1$, and $10$, from top to bottom). Dots indicate median posterior estimates and bars indicate 95% credible intervals. Dashed horizontal lines denote the true migration rates. The x axes indicate sampling parameters that varied across simulations: total number of sequences, sampling rate, proportion sampled, or sampling probability, depending on the panel. Across these additional scenarios, estimates remained broadly unbiased, with wider credible intervals in settings where the phylogenetic signal was weaker.

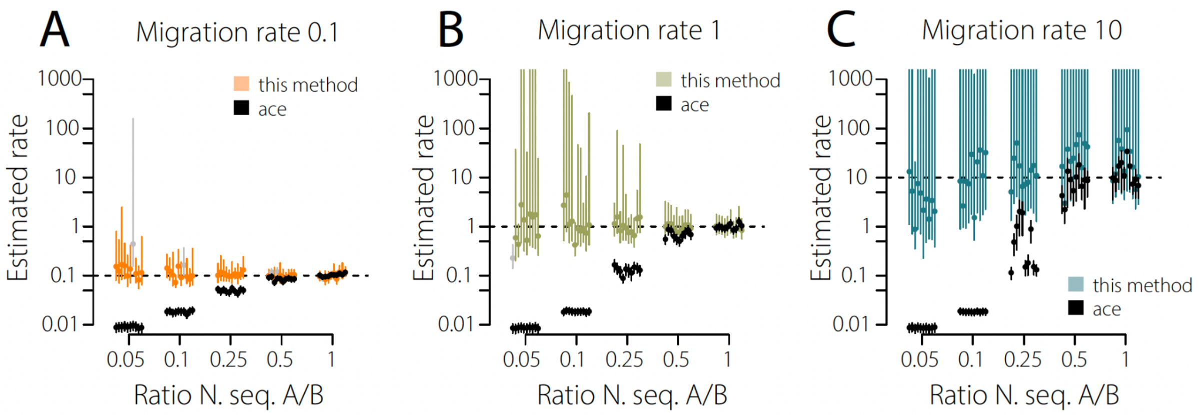


**Figure S6: Comparison of our method with estimates from ace**

Comparison between migration rates estimated by our approach and by the *ace* ancestral state reconstruction method (*36*) on simulated data under uneven sampling between two states. Panels show simulations with true migration rates per unit of time of 0.1 **(A)**, 1 **(B)**, and 10 **(C)**. The x axes show the ratio of the number of sampled sequences in states A and B. For example 0.05 indicates that state A has been sampled 20 times less than state B, while 1 indicates equal sampling. Dots indicate median posterior or point estimates for ace, and bars indicate 95% credible intervals for our method, or 95% confidence intervals for ace. The dashed horizontal line denotes the true migration rate. Estimates from our method are shown in colour (or grey if they don't cover the true rate), and estimates from ace are shown in black. Across all three migration regimes, our method remained broadly unbiased as the sampling imbalance increased, whereas ace estimates shifted systematically with the ratio of sampled sequences between states, consistent with sensitivity to uneven sampling intensity.

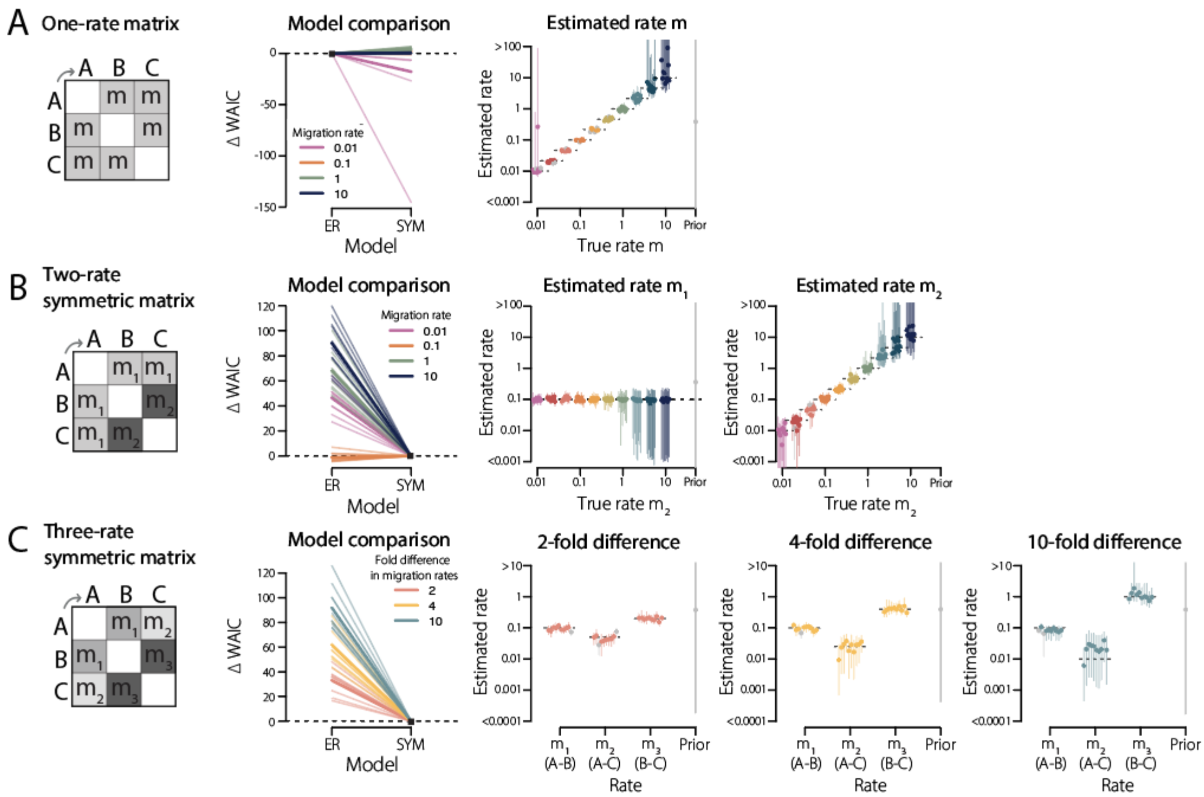


**Figure S7: Fitting symmetric three-state migration matrices**

Performance of our framework on simulated phylogenies under symmetric three-state migration matrices. Each state has an effective population size of 1,000 at present, growing exponentially at rate 0.1, with 1,000 sequences sampled per state and 10 trees per condition. **A.** One-rate matrix, in which all transitions between states occur at the same rate $m$. **B.** Two-rate symmetric matrix, with two distinct migration rates ($m_1$ and $m_2$). **C.** Three-rate symmetric matrix, with three distinct pairwise migration rates ($m_1$, $m_2$ and $m_3$). In each row, the left panel shows the true matrix used in simulations. The middle-left panel shows model comparison between alternative matrix structures using $\Delta\ WAIC$, relative to the true model (black square); lower values indicate better fit. ER: equal-rate model, SYM: symmetric model. Lines are coloured by the migration rate used in the simulation (A and B) or by the fold-difference between rates (C). Thin lines show individual trees and thick lines their mean. The remaining panels show estimated migration rates under the true model (ER in A, SYM in B and C), as a function of the true rates. Dots indicate median posterior estimates and bars indicate 95% credible intervals. Dashed lines denote the true values, and the grey distributions at the right indicate the prior.

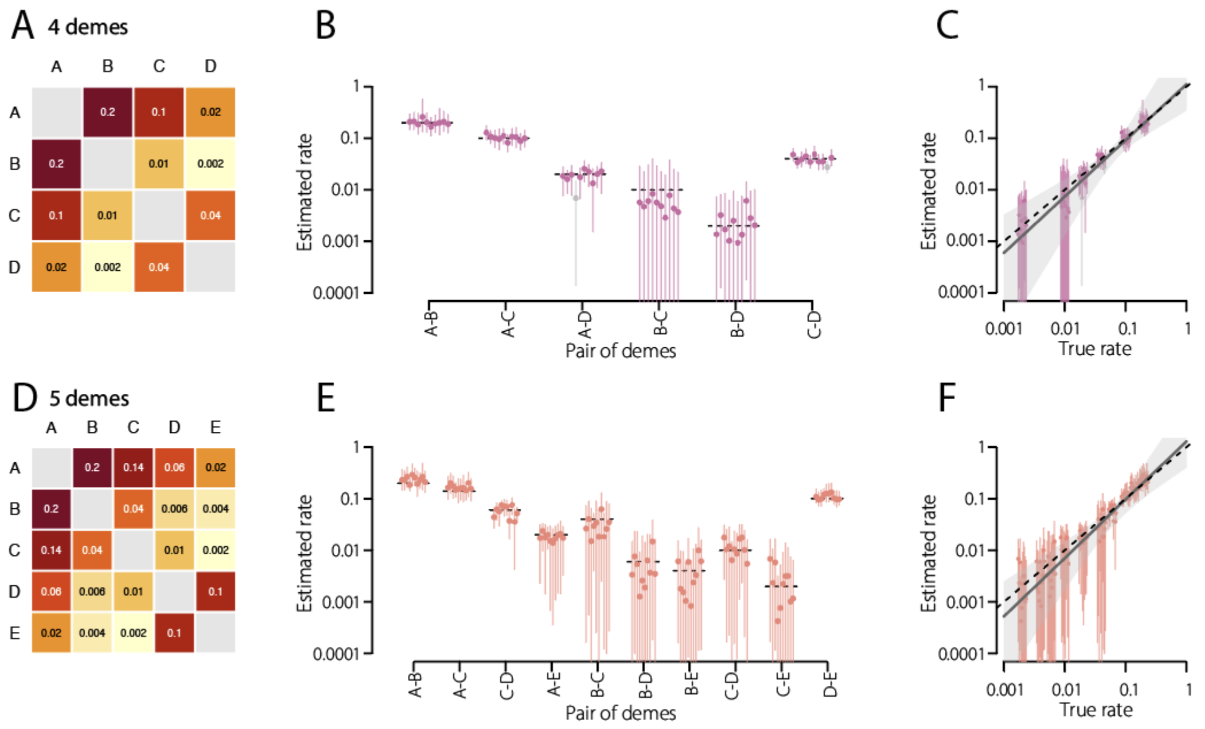


**Figure S8: Fitting four- and five- state migration matrices.**
Performance of the symmetric (SYM) model on phylogenies simulated under symmetric migration between four **(A–C)** and five **(D–F)** states, with distinct pairwise rates spanning 100-fold (0.002 to 0.2). Each state has an effective population size of 1,000 at present, growing exponentially at rate 0.1, with 1,000 sequences sampled per state and 10 trees per matrix. **A and D.** True migration matrices, coloured by rate (log scale). **B and E.** Estimated migration rate for each pair of states. Dots indicate posterior medians and bars 95% credible intervals for each tree. **C and F.** Estimated versus true migration rates across all pairs of states. The grey line and shaded area show the median and 95% interval of log-log regressions fitted to the posterior draws of each tree, accounting for the matrix structure. The dashed line denotes identity.

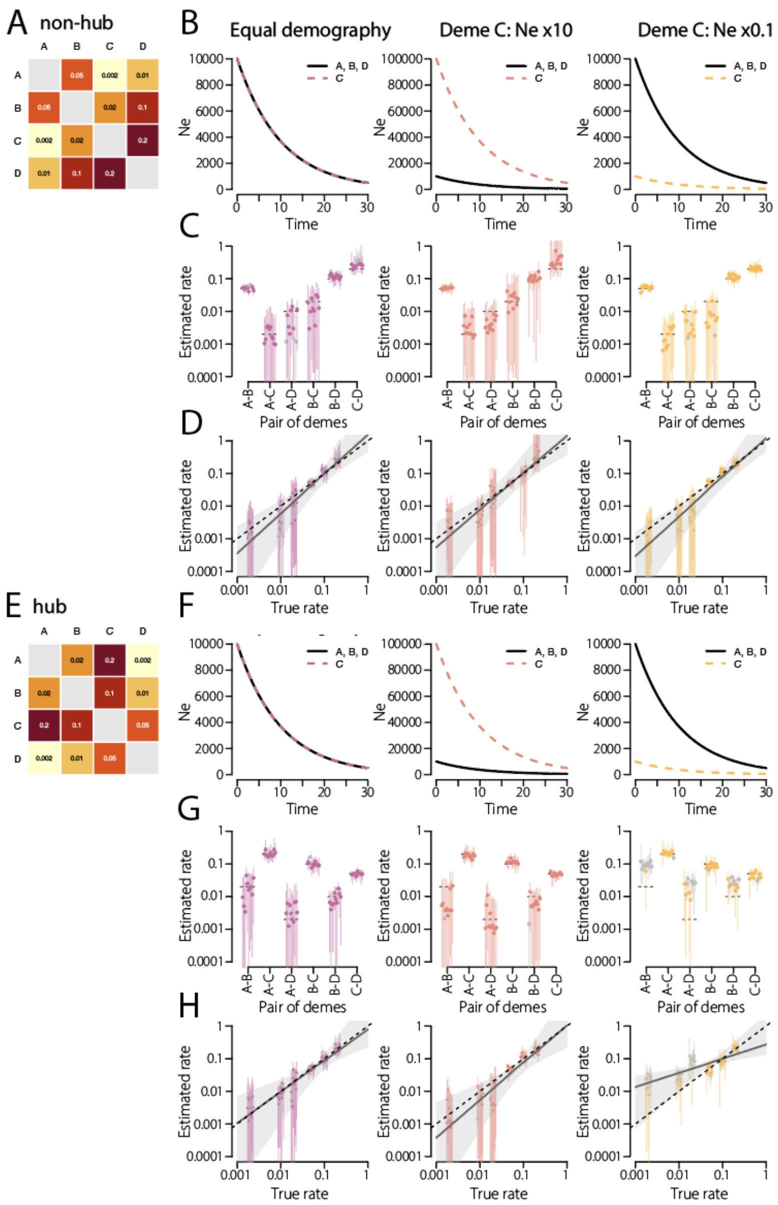


**Figure S9: Unequal demography in multi-deme systems**

Performance of the symmetric (SYM) model on phylogenies simulated under symmetric migration between four states, when the demography of one state, C, differs from the others. All other states have the same effective population sizes growing exponentially at rate 0.1. Ten trees were simulated per scenario, with 1,000 sequences sampled per state. The same six pairwise rates (0.002 to 0.2) are either distributed across pairs (A–D, non-hub) or assigned so that C has the three highest rates (E–H, hub). **A and E.** True migration matrices, coloured by rate (log scale). **B and F.** Effective population sizes through time of the different states. The initial population size of state C is equal to the others (10,000 at present; left), ten times larger (middle) or ten times smaller (right). **C and G.** Estimated migration rate for each pair of states. Dots indicate posterior medians and bars 95% credible intervals. **D and H.** Estimated versus true migration rates across all pairs of states. The grey line and shaded area show the median and 95% interval of log-log regressions fitted to the posterior draws of each tree, accounting for the matrix structure. The dashed line denotes identity. When the hub C is ten times smaller than the other states, the rates between the other, weakly connected states are overestimated.

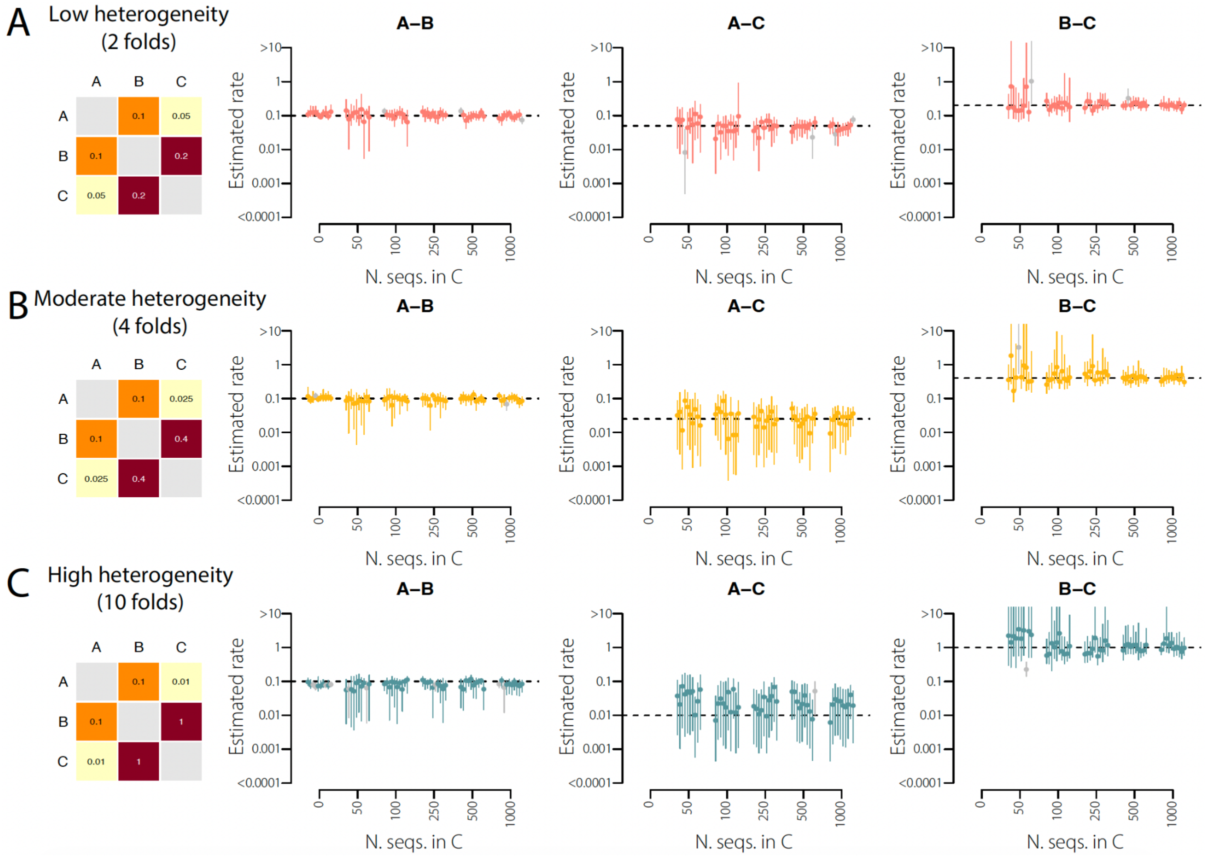


**Figure S10: Fitting a three deme matrix in the presence of uneven sampling**

Performance of our framework on phylogenies simulated under symmetric migration and when state C is undersampled. Each state has an effective population size. We sampled 1,000 sequences from A and B, and 0 (C unsampled), 50, 100, 250, 500 or 1,000 sequences from C, with 10 trees per scenario. **A.** Case of low heterogeneity between migration rates **B.** Moderate heterogeneity. **C.** High heterogeneity. Left: true migration matrices, coloured by rate (log scale). Right: estimated migration rate for each pair of states as a function of the number of sequences sampled from C. When C is not sampled, only the A–B rate is estimated, with a two-state model. Dots indicate medians and bars 95% credible intervals. Undersampling C widens the credible intervals of the rates involving C but does not bias the estimates, and the A–B connectivity is recovered even when C is not sampled (ghost deme).

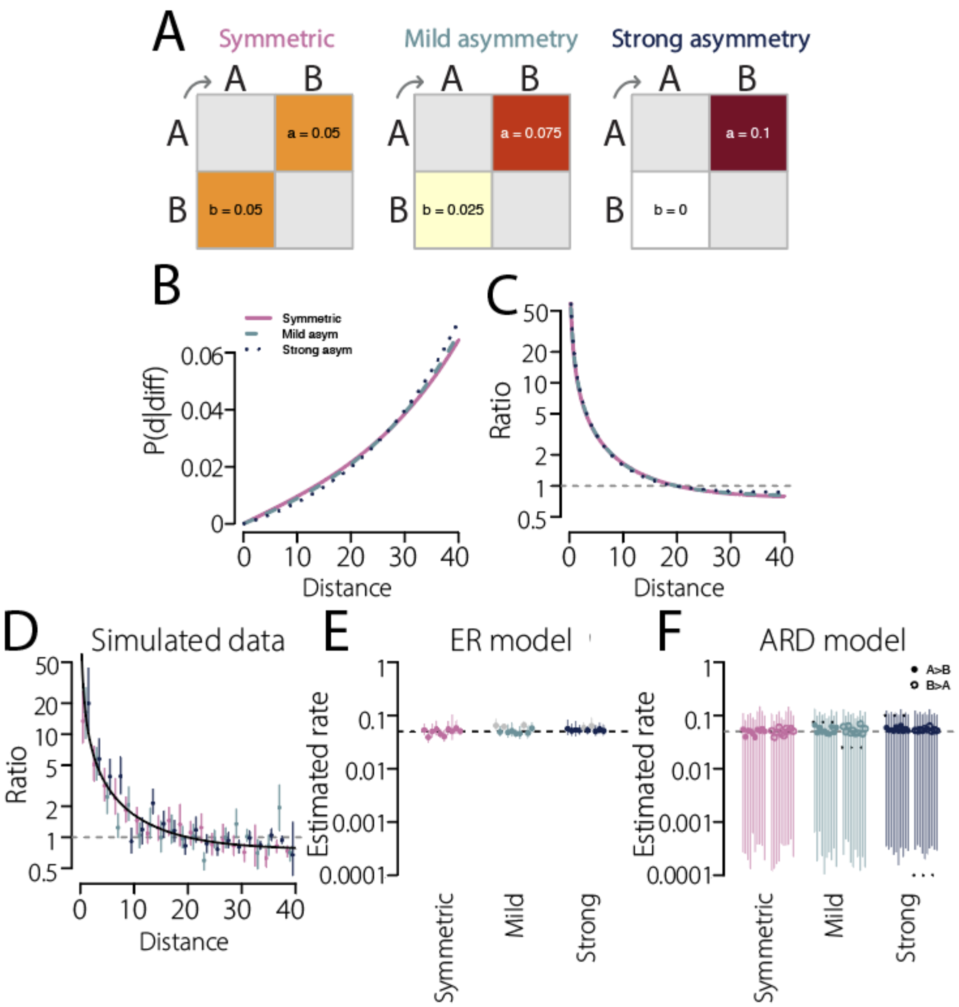


**Figure S11: Non-identifiability of asymmetric migration and estimation of the average migration rate in a two-deme population**

**A.** Migration matrices compared, all with the same total migration rate (a + b)/2 = 0.05, and increasing asymmetry: from left to right, symmetric (a = b = 0.05), mild asymmetry (a = 0.075, b = 0.025) and strong asymmetry (a = 0.1, b = 0). a is the migration rate from A to B and b from B to A. Cells are coloured by migration rate (log scale). **B.** Theoretical distribution of pairwise distance distributions for sequence pairs sampled in different states. Colours denote the different matrices. **C.** Theoretical ratio of the pairwise distance distributions for sequence pairs sampled in the same state versus different states ($Ne(t) = 1000 * exp(-0.1t)$). In B and C, distributions are computed with the structured coalescent, with each state having the same effective population size. **D.** Ratio computed on one simulated tree under each of the 3 different matrices. Dots indicate medians and bars indicate 95% intervals over 200 bootstrap replicates. The black line shows the theoretical ratio for the symmetric matrix. **E.** Migration rates estimated with a symmetric model (ER) for 10 simulated trees. Dots indicate median posterior estimates and bars indicate 95% credible intervals. Estimates that do not cover the average rate are shown in grey. **F.** Migration rates estimated with an asymmetric model (ARD), from A to B (solid dots) and B to A (empty dots). Dotted lines show the true rates. In E and F, the dashed line shows the average migration rate (a + b)/2 = 0.05. The distributions and ratios are nearly identical across the three matrices, so the direction and strength of the asymmetry are not captured by our model, whereas the average migration rate is consistently recovered.

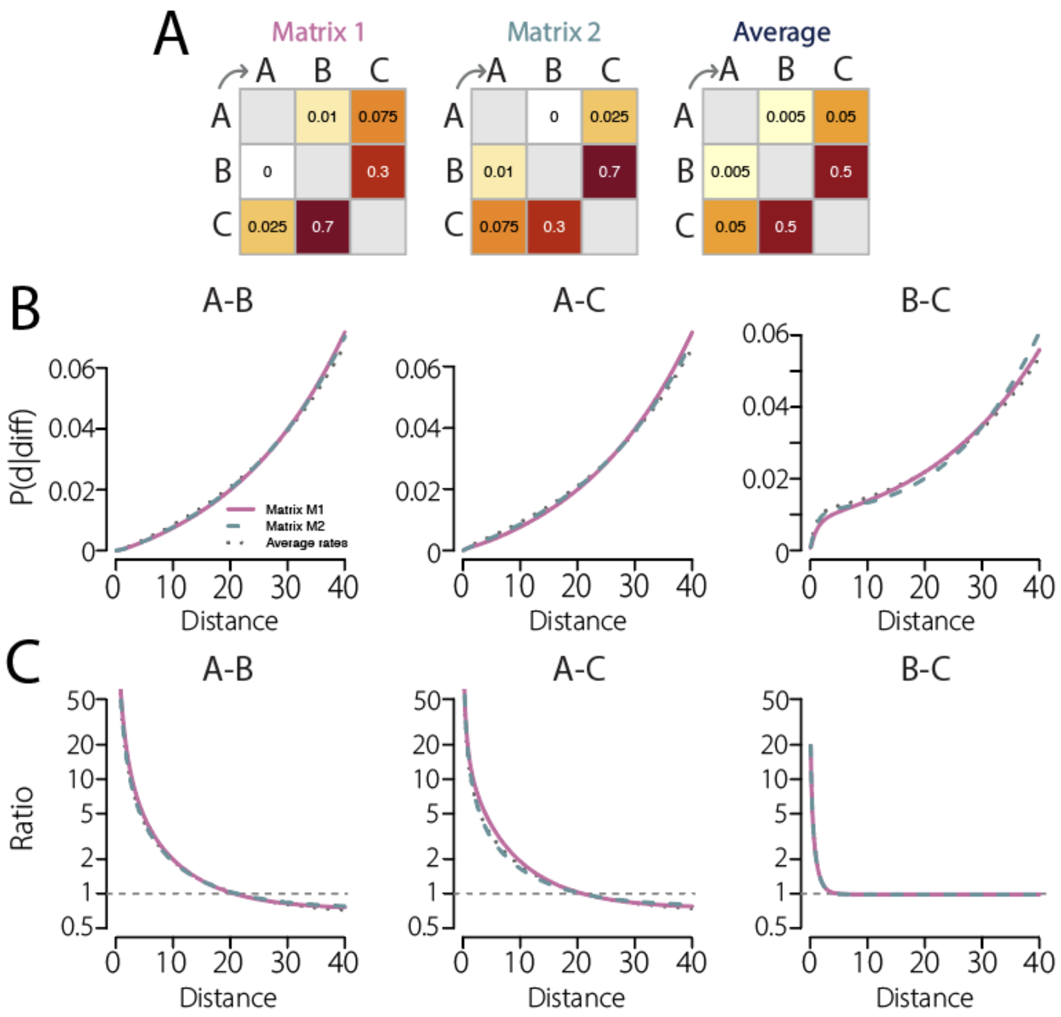


**Figure S12: Theoretical signal of asymmetric migration and non-identifiability of individual rates in a three-deme population**

Theoretical pairwise-distance signal under a three-state structured coalescent model with exponentially growing effective population size ($Ne(t) = 1000 * exp(-0.1t)$), and three distinct asymmetric backward-time migration matrices shown in **A**. **B.** Direct comparison of $P(d|diff)$ for each pair of demes. **C.** Direct comparison of the ratios of within- to between-deme pairwise distance distributions. The dashed horizontal line indicates a ratio of 1. In B and C, colours denote the different matrices used. Although the individual migration rates differ between the three matrices, they generate similar overall connectivity between deme pairs, making individual rates unidentifiable by our model.

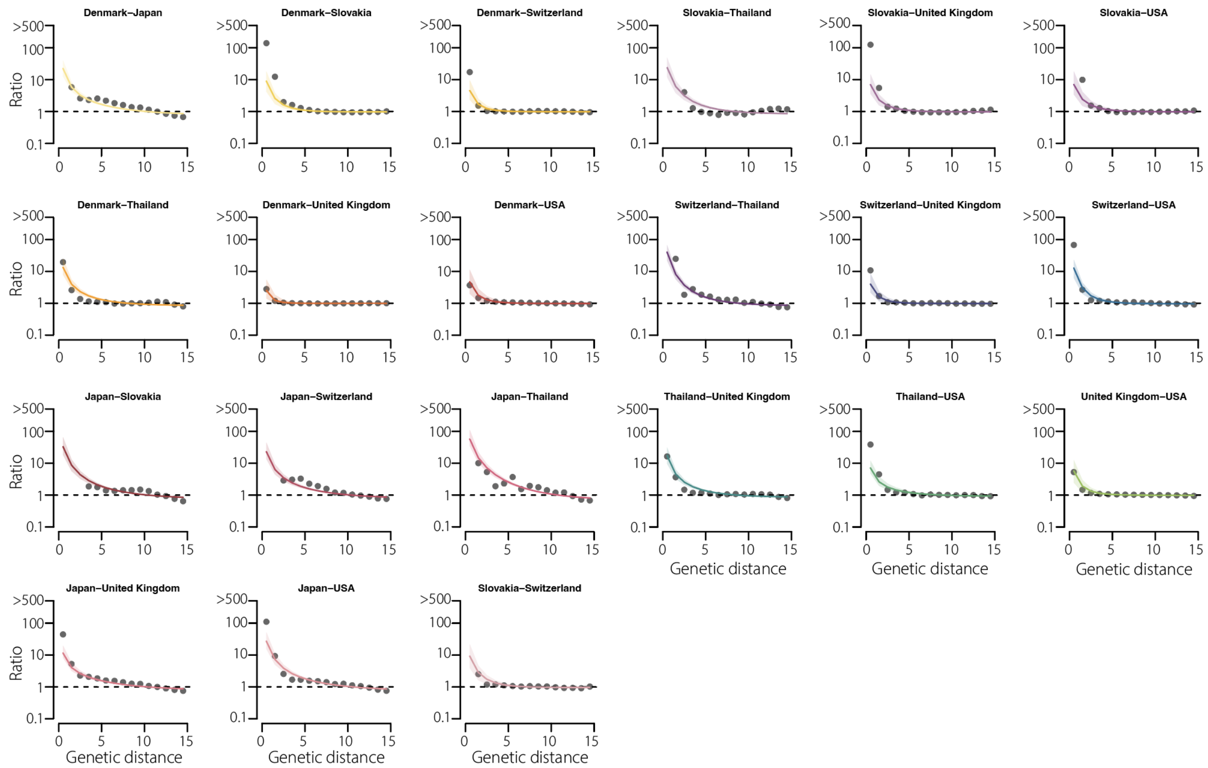


**Figure S13: Model fits by country pair for SARS-CoV-2 spread between countries**

Model fits of the ratio of pairwise genetic distance distributions for sequences sampled within versus between countries, shown separately for each unordered country pair. In each panel, black dots denote the observed ratio at each genetic distance bin. Coloured lines and shaded area show the fitted ratio for the corresponding country pair (median and 95% credible intervals, respectively). The dashed horizontal line indicates a ratio of 1.

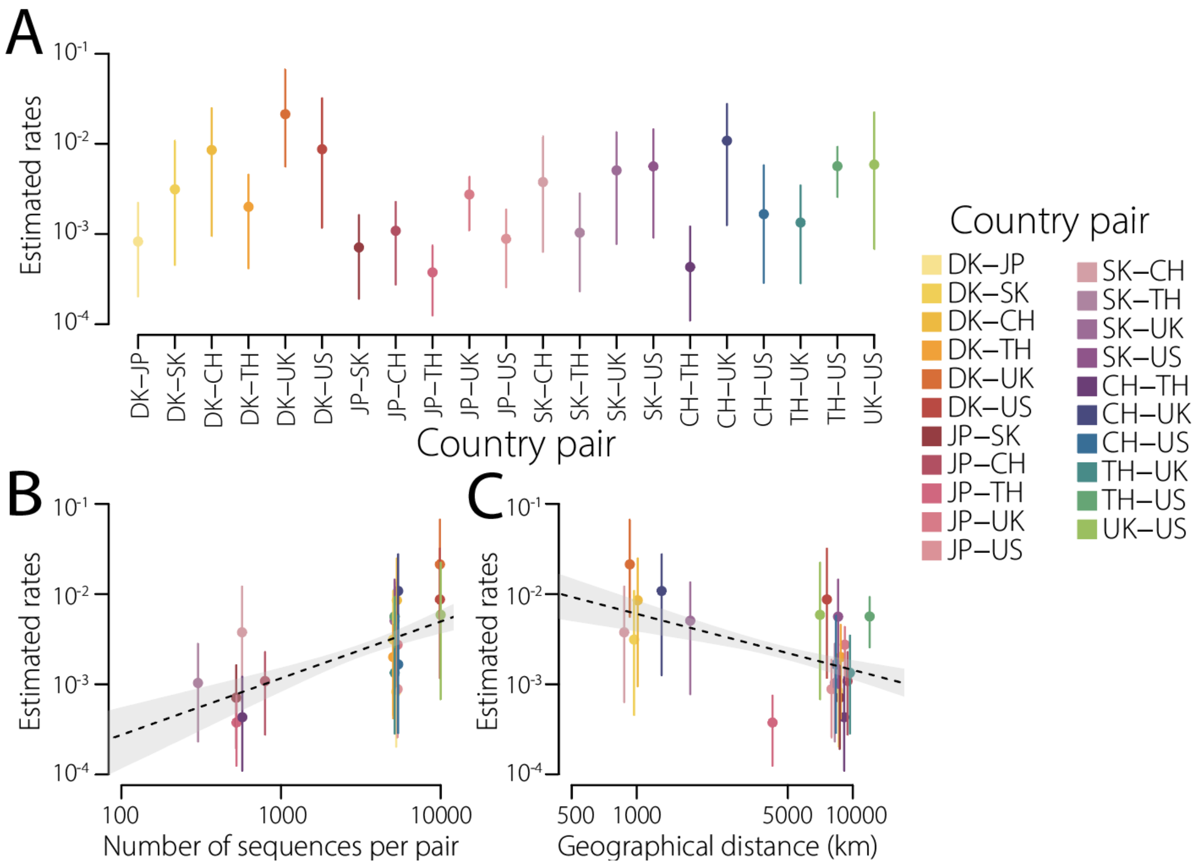


**Figure S14: Estimated SARS-CoV-2 transition rates between countries and comparison with covariates**

**A.** Estimated transition rates between all country pairs for SARS-CoV-2. Dots indicate median posterior estimates and bars indicate 95% credible intervals. Country pairs are coloured individually. **B.** Relationship between estimated transition rates (log scale) and the total number of sequences available for each country pair (log scale). **C.** Relationship between estimated transition rates (log scale) and geographical distance between countries (log scale). In B and C, the dashed line shows the fitted log-log regression and the shaded area the corresponding confidence region, accounting for the matrix structure (p value = 0.012 and 0.066, respectively).

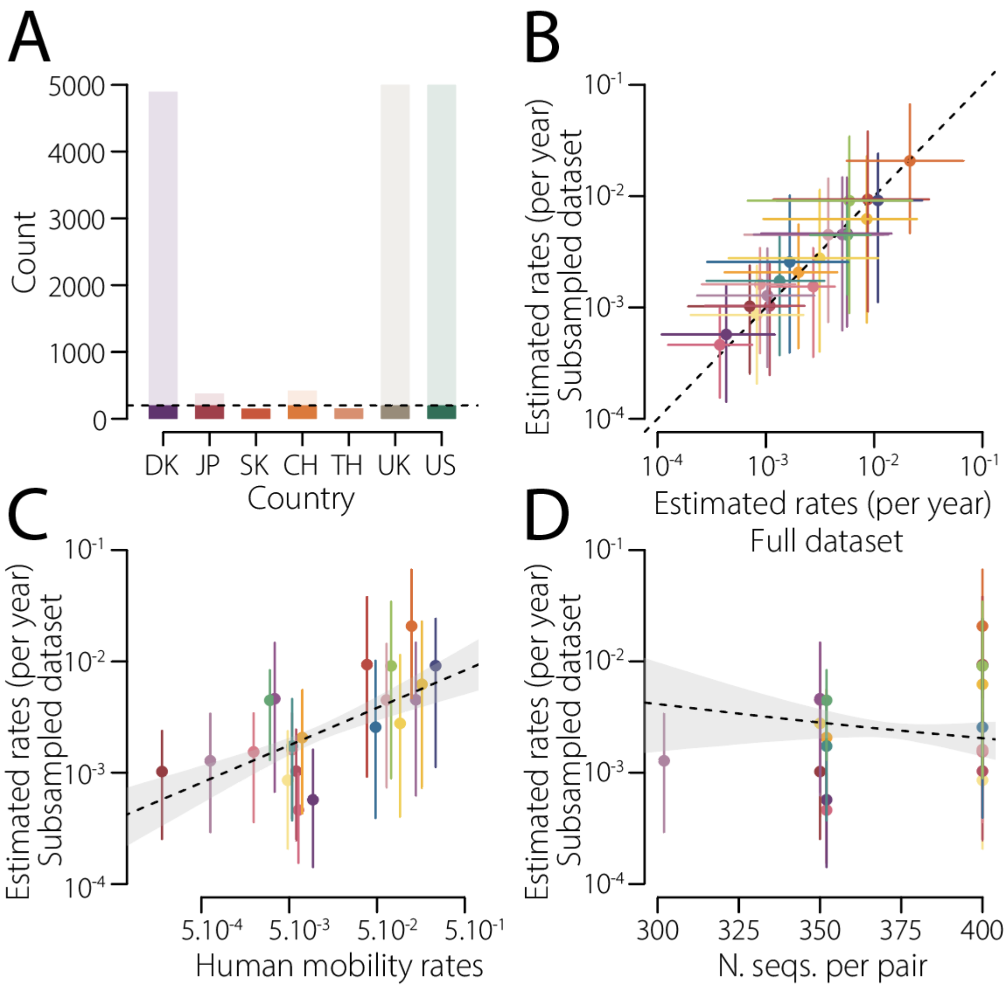


**Figure S15: Robustness of estimated transition rates between countries to subsampling of the SARS-CoV-2 dataset.**

**A.** Number of genomes available from each country in the full dataset (pale bars) and retained in the subsampled dataset (coloured bars). The dashed line represents the maximum of sequences retained per country: 200. DK, Denmark; JP, Japan; SK, Slovakia; CH, Switzerland; TH, Thailand; UK, United Kingdom; US, United States. **B.** Transition rates estimated from the subsampled dataset (y axis) compared with those estimated from the full dataset (x axis). Each point represents a country pair; error bars indicate 95% credible intervals, and the dashed line indicates identity. **C.** Relationship between estimated transition rates estimated from the subsampled dataset (y axis, log scale) and human mobility rates per year (x axis, log scale). **D.** Relationship between estimated transition rates (log scale) and the total number of sequences available for each country pair. In C and D, the dashed line shows the fitted regression and the shaded area the corresponding confidence region, accounting for the matrix structure. The estimated rates remain significantly associated with human mobility rates (p value = 0.023), while not being associated with the number of sequences per pair (p value = 0.238).

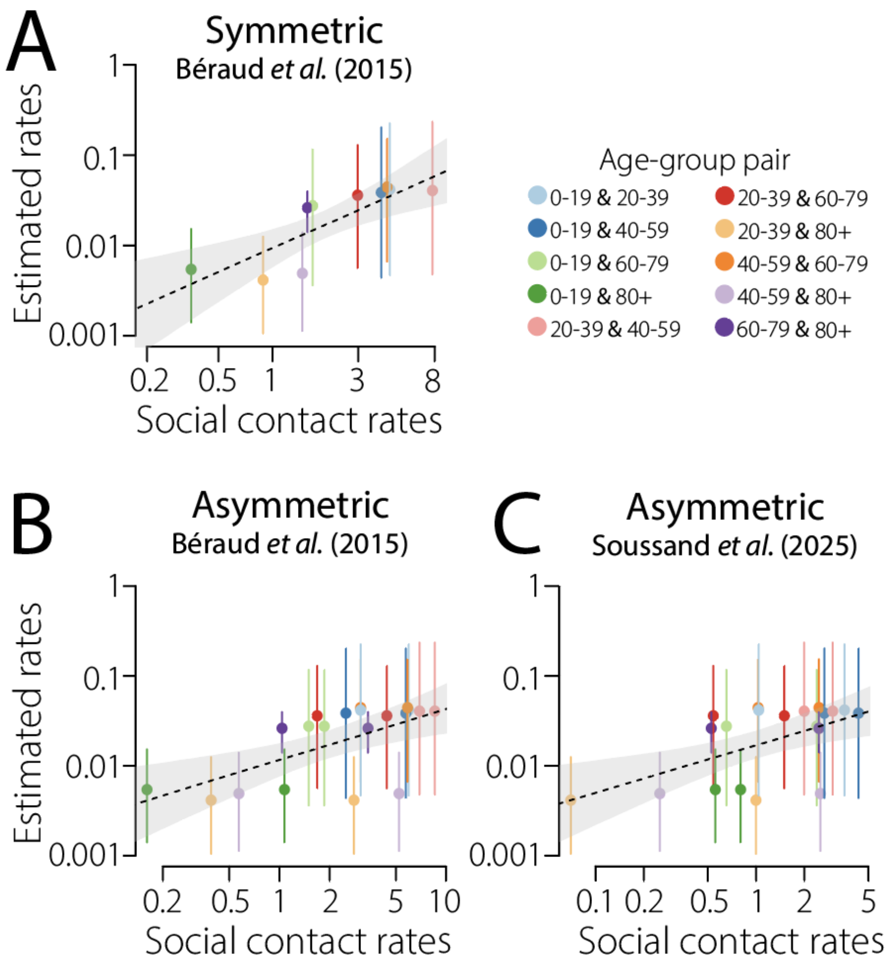


**Figure S16: Comparison of estimated SARS-CoV-2 transition rates across age groups using different contact matrices**

Comparison between our estimated transition rates across age-group pairs for SARS-CoV-2 and independent estimates of social contact rates under three alternative contact matrices. **A.** Symmetric contact matrix from Béraud et al. (2015) (*44*). **B.** Asymmetric contact matrix from Béraud et al. (2015) (*44*). **C.** Asymmetric contact matrix from SocialCov (*28*). In all panels, dots indicate median estimated transition rates for each age-group pair and bars indicate 95% credible intervals. Colours denote age-group pairs. The dashed line shows the fitted log-log regression and the shaded area the corresponding confidence region, accounting for the matrix structure. All regressions indicate a significant association between contact rates estimates and the contact patterns in the population (p values of 0.008, 0.008 and 0.008, in A, B and C, respectively).

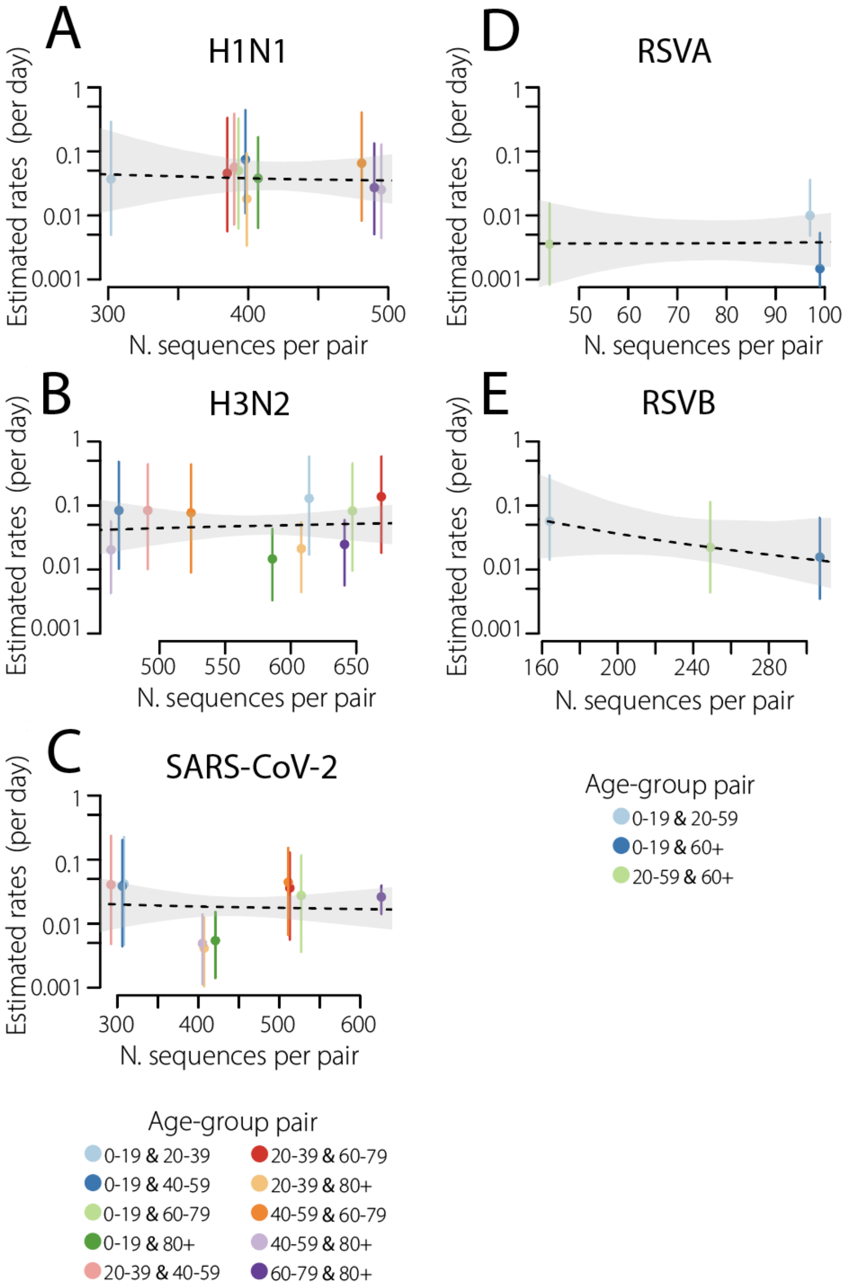


**Figure S17: Comparison of estimated transition rates across age groups with sampling intensity for SARS-CoV-2, H1N1, H3N2, RSV-A, and RSV-B**

Relationship between estimated transition rates and the number of sequences available for each age-group pair, shown separately for H1N1 **(A)**, H3N2 **(B)**, SARS-CoV-2 **(C)**, RSV-A **(D)**, and RSV-B **(E)**. Dots indicate median estimated transition rates and bars indicate 95% credible intervals. Colours denote age-group pairs. The dashed line shows the fitted regression and the shaded area the corresponding confidence region, accounting for the matrix structure.

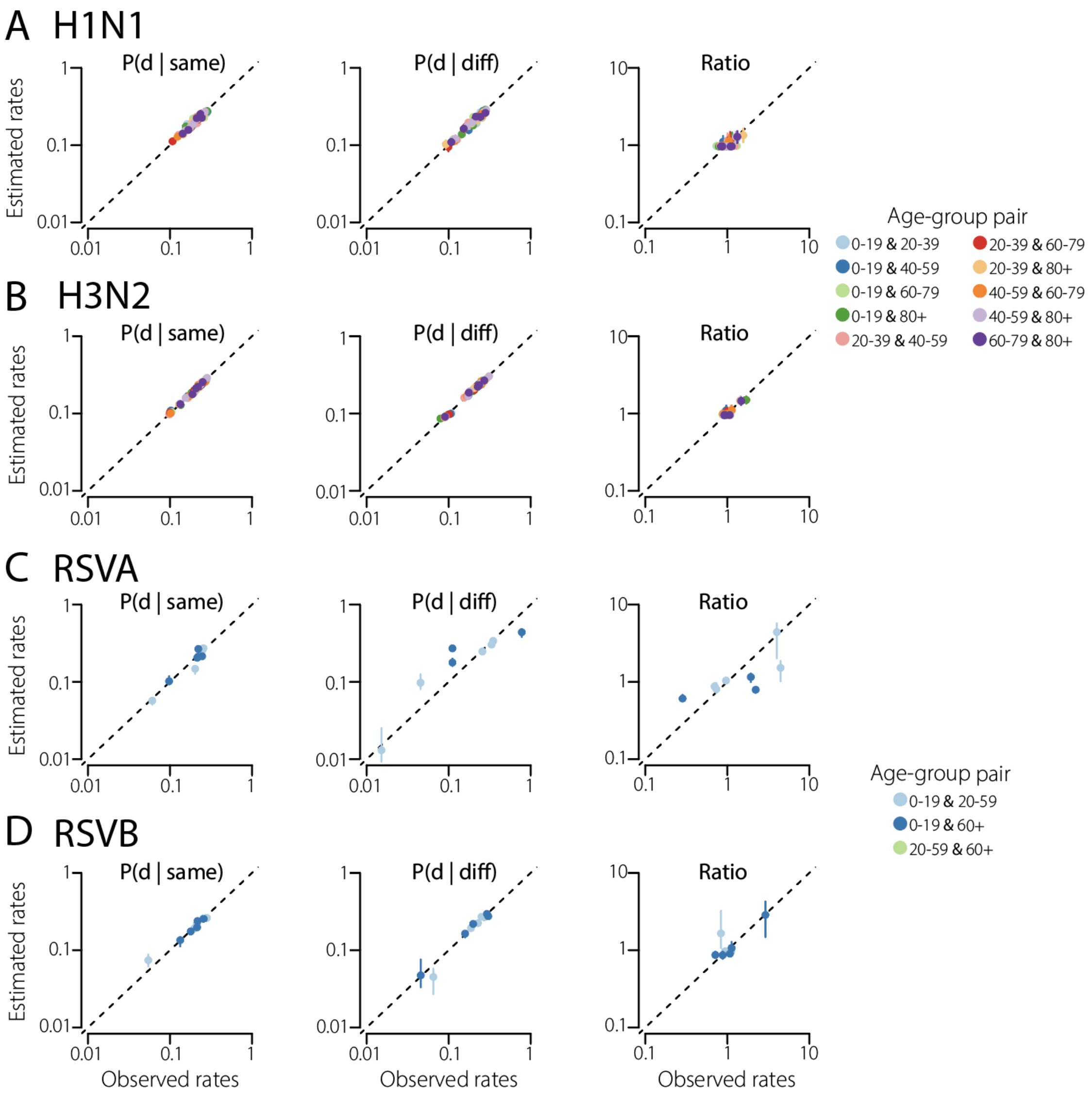


**Figure S18: Model fits for spread across age groups in H1N1, H3N2, RSV-A, and RSV-B**

Model fits of pairwise genetic distance distributions for transmission across age groups, shown separately for H1N1 **(A)**, H3N2 **(B)**, RSV-A **(C)**, and RSV-B **(D)**. For each pathogen, the three columns compare observed and predicted values for $\boldsymbol{P(d \mid same)}$, $\boldsymbol{P(d \mid diff)}$, and their ratio, respectively. Points are coloured by age-group pair. The dashed line denotes identity. Across pathogens, the model accurately reconstructed the observed pairwise distance distributions within and between age groups, although the ratio was more weakly captured for pathogens with limited age structure or greater uncertainty, particularly RSV-A and RSV-B.

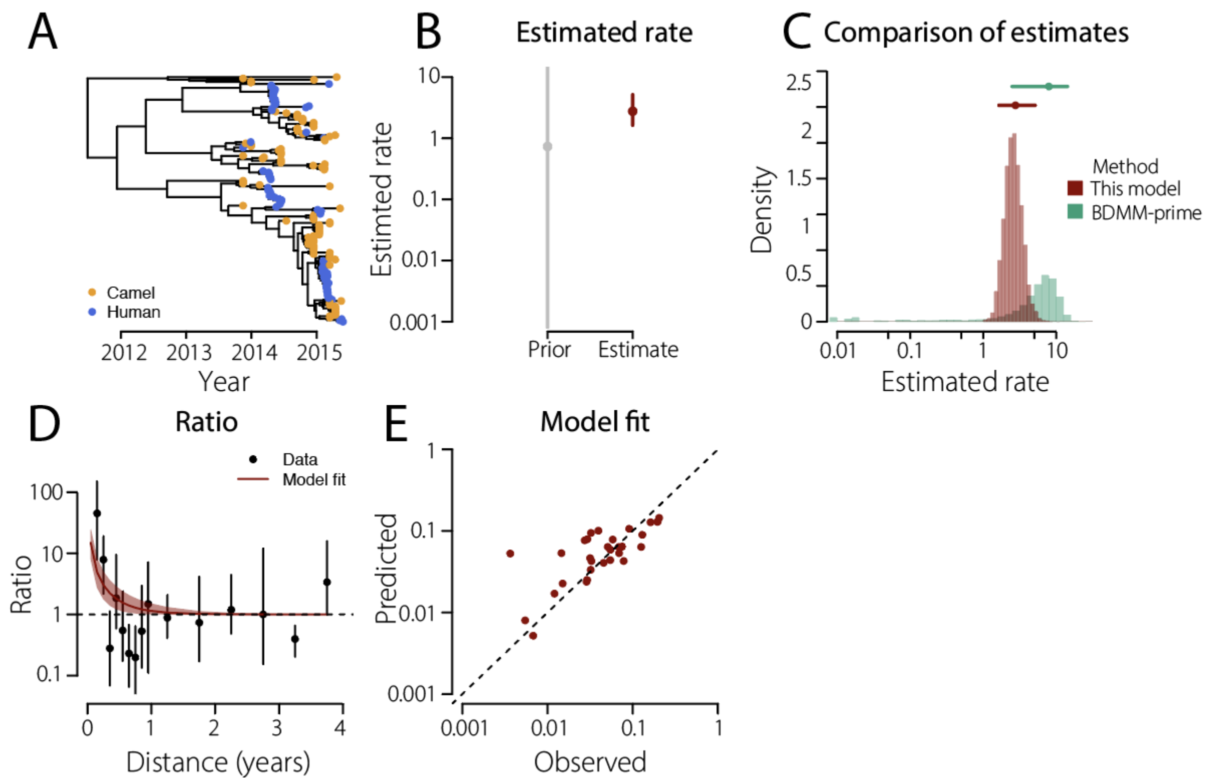


**Figure S19: Estimating the transmission rate between camels and humans in MERS-CoV**

Application of the equal-rate model to 178 MERS-CoV genomes sampled from camels and humans. As our model does not capture the direction of transmission, we assumed no human-to-camel transmission, as suggested by previous studies (*7*, *15*). Under this assumption, the camel-to-human rate corresponds to twice the rate estimated with the equal-rate model (Figure S11). **A.** Time-scaled phylogeny coloured by host, showing camel-derived (orange) and human-derived (blue) sequences. **B.** Estimated camel-to-human rate (per year) expressed as the backward-in-time rate at which a lineage sampled in humans moves to camels. The posterior median and 95% credible intervals are shown in red. The prior is shown in grey. **C.** Comparison of this rate estimated by our model (red, 2.76 (1.59–5.08)) with the equivalent one by BDMM-prime parameter estimates (*7*) (green, 7.91 (2.48–14.24), see Methods)**.** Histograms show the posterior distributions. Dots and bars represent the median and 95% credible intervals, respectively. **D.** Ratio of pairwise genetic distance distributions for sequences sampled from the same host versus different hosts. Data is shown in black, the model fit in red. Black dots denote medians and bars denote 95% confidence intervals over bootstrap replicates. The red line and shaded area denote the median and 95% credible interval of the model fit. The dashed horizontal line indicates a ratio of 1. **E.** Model fit comparing observed and predicted pairwise distance distributions. The dashed line denotes identity.

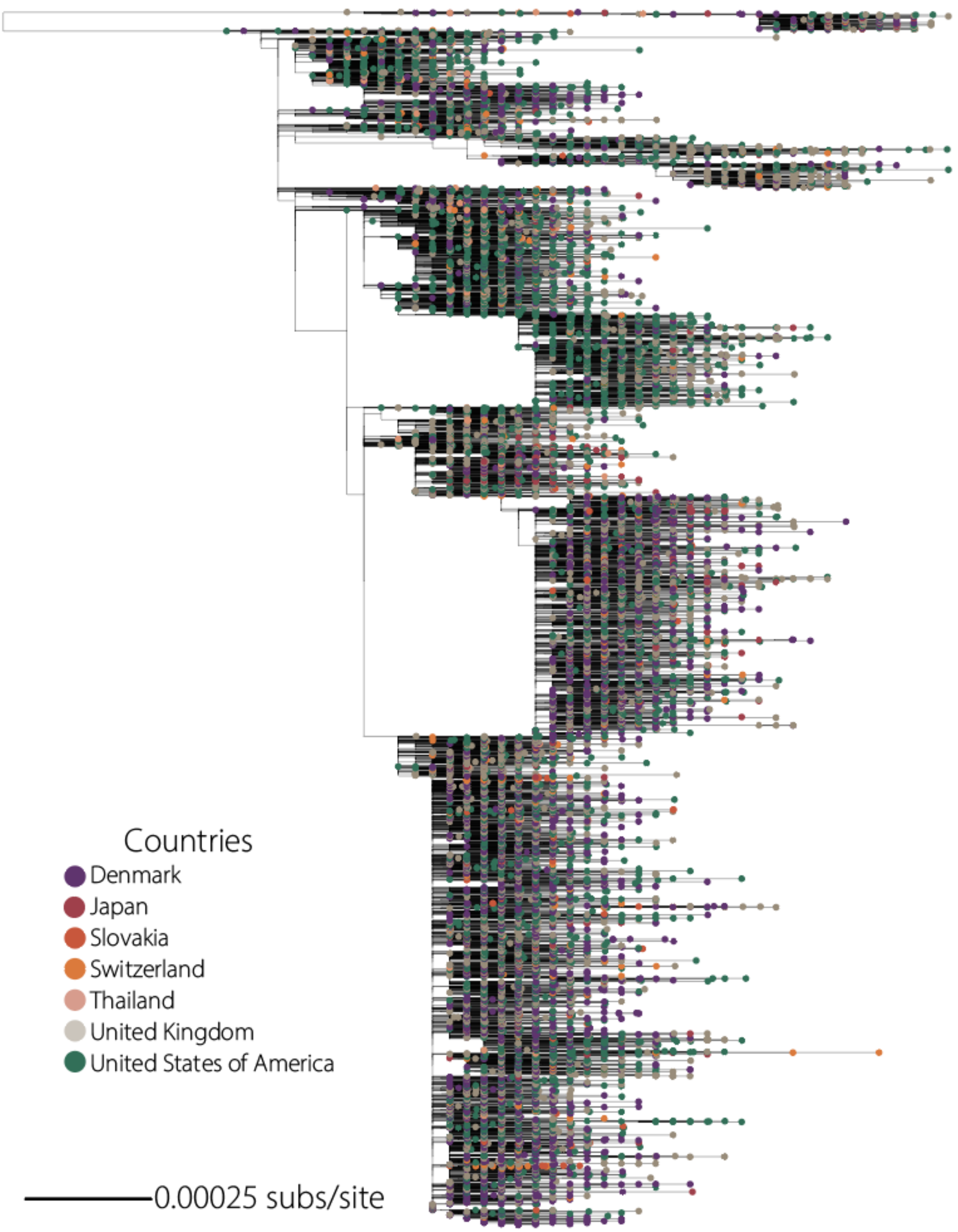


**Figure S20: SARS-CoV-2 phylogeny of sequences from seven countries, coloured by country**
Midpoint-rooted phylogeny of 15,991 SARS-CoV-2 genomes sampled between July 2024 and June 2025, coloured by country of sampling.

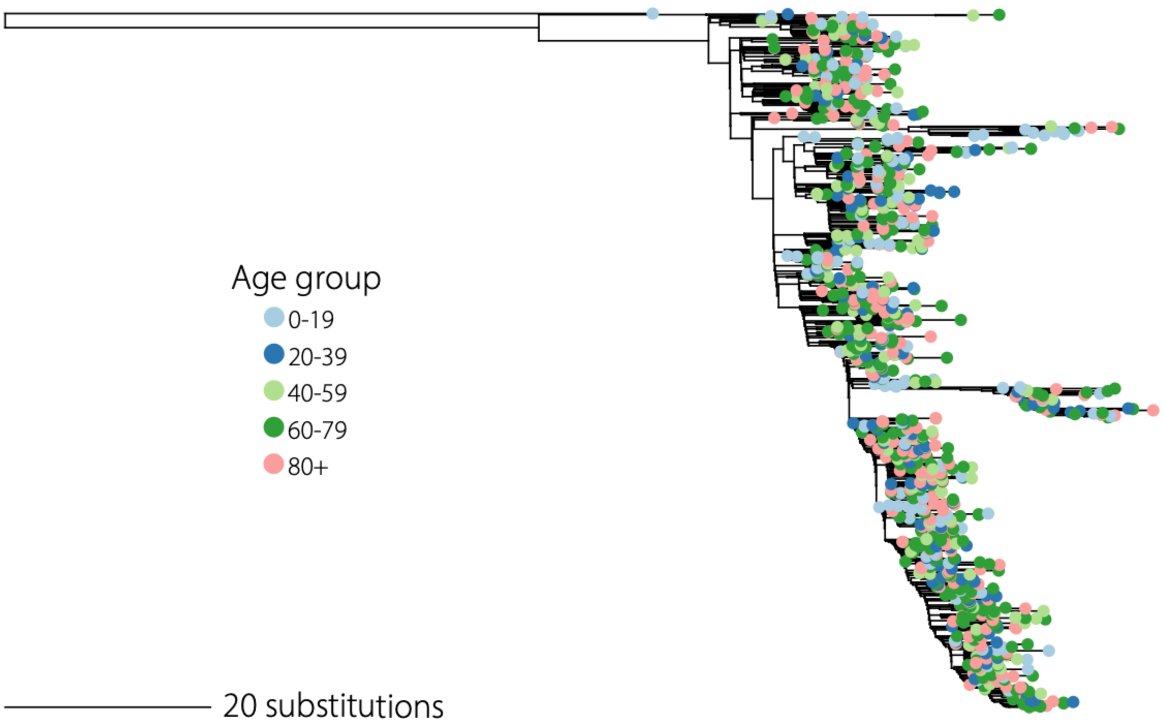


**Figure S21: SARS-CoV-2 phylogeny of sequences from Auvergne-Rhône-Alpes, France, coloured by age group**

Phylogeny of SARS-CoV-2 genomes sampled in the Auvergne-Rhône-Alpes region of France during the 2024–2025 season, with tips coloured by age group: 0–19, 20–39, 40–59, 60–79, and 80+ years.

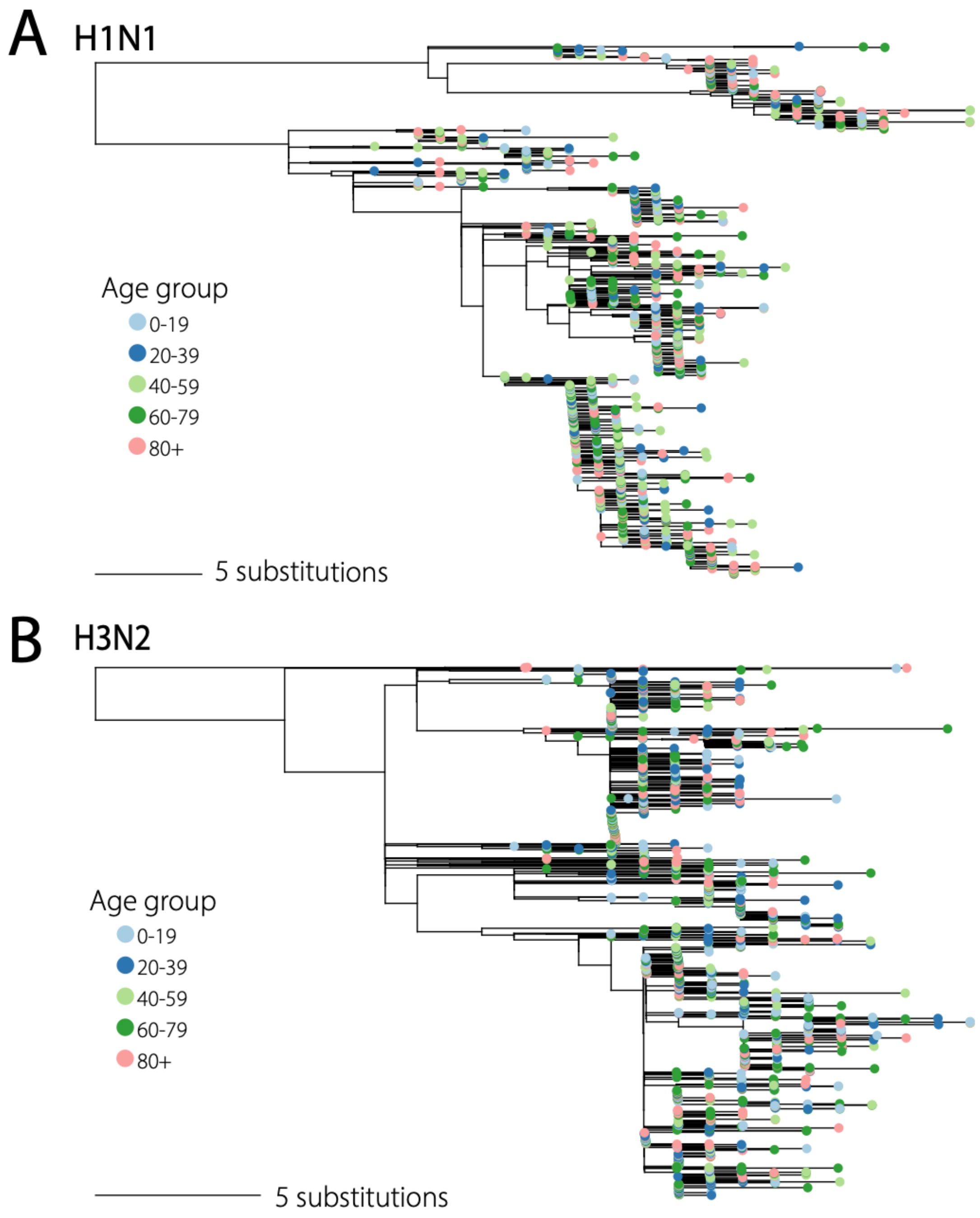


**Figure S22: Influenza A/H1N1 and H3N2 phylogenies of sequences from Auvergne-Rhône-Alpes, France, coloured by age group**

Phylogenies of influenza H1N1 **(A)** and H3N2 **(B)** genomes sampled in the Auvergne-Rhône-Alpes region of France, with tips coloured by age group: 0–19, 20–39, 40–59, 60–79, and 80+ years.

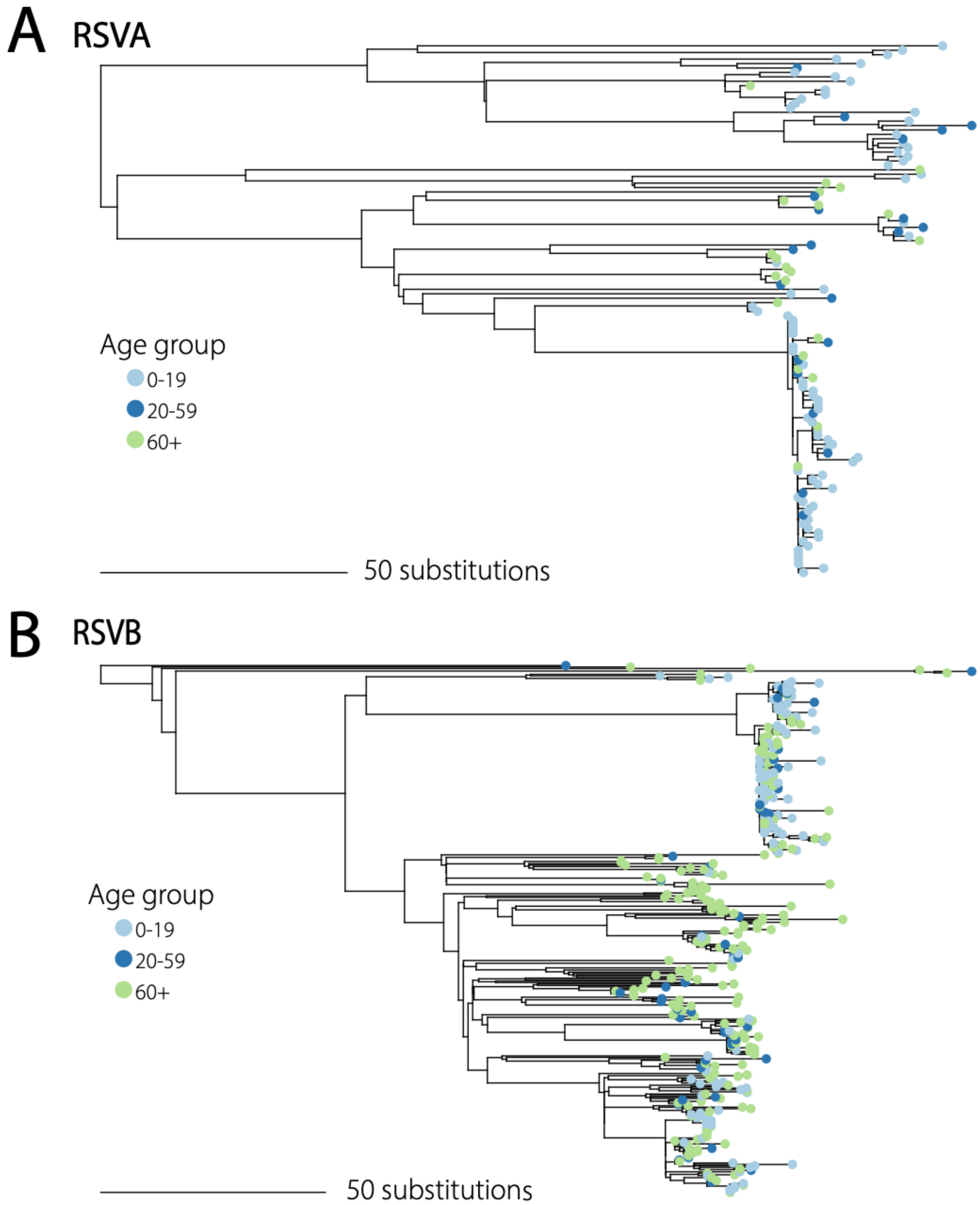


**Figure S23: RSV-A and RSV-B phylogenies of sequences from Auvergne-Rhône-Alpes, France, coloured by age group**

Phylogenies of RSV-A (**A**) and RSV-B (**B**) genomes sampled in the Auvergne-Rhône-Alpes region of France, with tips coloured by age group: 0–19, 20–59, and 60+ years.

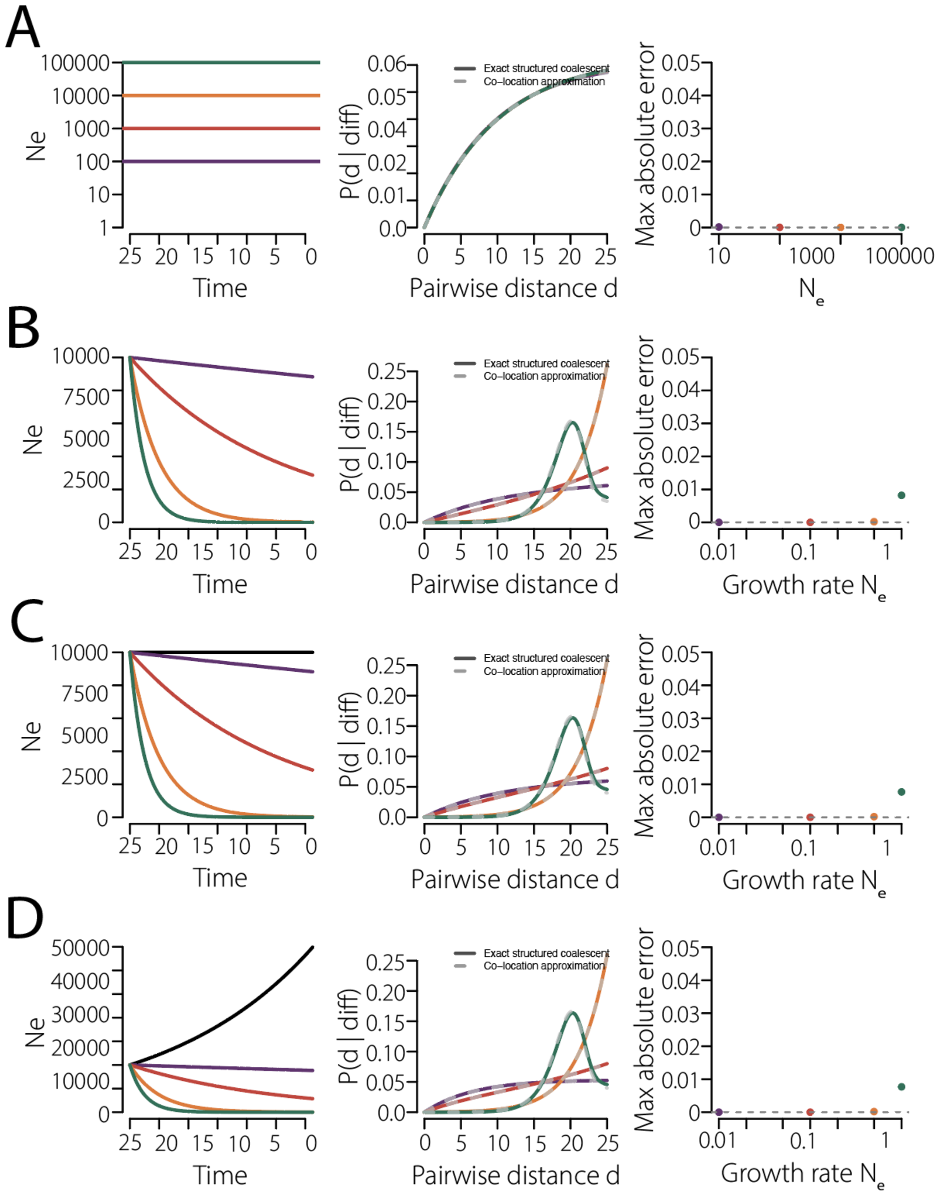


**Figure S24: Approximation versus true P(d | diff) distributions**

Comparison between the exact structured coalescent and our approximation for $\boldsymbol{P(d \mid diff)}$ under different effective population size scenarios. In each row, the left panel shows the effective population size trajectory through time, the middle panel shows the corresponding $\boldsymbol{P(d \mid diff)}$ distributions obtained under the exact structured coalescent (solid lines) and our approximation (dashed lines), and the right panel shows the maximum absolute error between the two across pairwise distances, for different population size trajectories. **A.** Constant effective population sizes across a range of values, with identical trajectories in the two demes. **B.** Exponentially growing effective population sizes, with identical trajectories in the two demes and varying growth rates. **C.** Exponentially growing effective population sizes in one deme, while the other deme remains constant (black line). **D.** Exponentially growing effective population sizes in one deme, while the other deme exponentially decreases (black line). Colours denote the different effective population size trajectories or growth rates shown in the left panel. Across all scenarios, our approximation closely matched the exact structured coalescent, with only small deviations, especially in the case of a low population size in one or both demes.

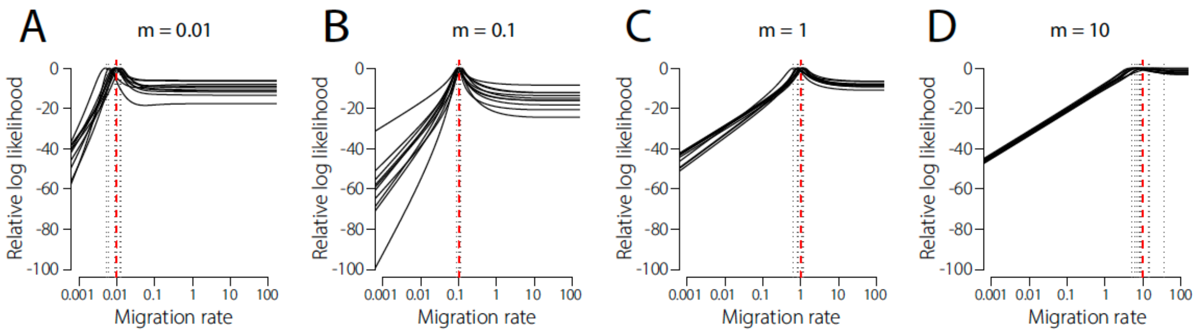


**Figure S25: Likelihood profiles across migration rates**

Likelihood profiles obtained for simulated datasets under four true migration rates: 0.01 **(A)**, 0.1 **(B)**, 1 **(C)** and 10 **(D)**. In each panel, black lines show the relative log-likelihood as a function of the migration rate for individual simulated phylogenies. The log-likelihood is plotted relative to its peak. The red dashed vertical line denotes the true migration rate used in the simulations, and the grey dotted vertical lines indicate the estimated migration rates for each profile.

### Supplementary Tables

**Table S1 (csv): Isolates of SARS-CoV-2 for the geographical spread analysis**
This table lists all 15,991 whole-genome SARS-CoV-2 sequences used in the geographical spread analysis. For each sequence, we list its NCBI accession number, country, collection date, release date, Pangolin lineage, Host, and Submitters.

**Table S2 (pdf): EPISET of isolates of SARS-CoV-2 for the age-group transmission analysis**
GISAID identifier of the whole-genome SARS-CoV-2 sequences used in the age-group transmission analysis (*20*).

**Table S3 (pdf): EPISET of isolates of H1N1 influenza for the age-group transmission analysis**
GISAID identifier of the HA H1N1 sequences used in the age-group transmission analysis (*21*).

**Table S4 (pdf): EPISET of isolates of H3N2 influenza for the age-group transmission analysis**
GISAID identifier of the HA H3N2 sequences used in the age-group transmission analysis (*22*).

**Table S5 (pdf): EPISET of isolates of RSV-A for the age-group transmission analysis**
GISAID identifier of the RSV-A whole-genome sequences used in the age-group transmission analysis(*23*).

**Table S6 (pdf): EPISET of isolates of RSV-B for the age-group transmission analysis**
GISAID identifier of the RSV-B whole-genome sequences used in the age-group transmission analysis (*24*).

**Table S7 (csv): Isolates of MERS-CoV**
This table lists all 178 MERS-CoV sequences used. For each sequence, we list its name (which includes its NCBI accession number), collection date, and host. This dataset is a subset of the one used in (*15*).
